# Selection shapes the evolution of genome size in a globally invasive plant

**DOI:** 10.64898/2026.01.27.702167

**Authors:** Byonkesh Nongthongbam, Paul Battlay, Katherine G. Maunder, Alexandre Fournier-Level, Michael D. Martin, John R. Stinchcombe, Kathryn A. Hodgins

## Abstract

- Biological invasions provide powerful natural experiments for understanding how genome architecture responds to novel climatic environments. Transposable elements (TEs) can rapidly restructure genomes, yet their role in adaptive genome size evolution during invasion remains poorly understood.
- Here, we examined genome size and TE abundance in 439 individuals of globally invasive common ragweed (*Ambrosia artemisiifolia* L.) across native (North American) and invasive (European, Australian) ranges. By integrating whole-genome resequencing, flow cytometry, and trait versus genetic differentiation comparison (*Q_ST_–F_ST_*), we tested whether genome size evolution is shaped by selection, climate, and life-history traits.
- Genome size was significantly larger in Australian genotypes, driven by increased TE and rRNA (ribosomal RNA) abundance. Crucially, trait versus genetic differentiation comparison provided evidence of divergent selection on genome size in North American and European populations, but not in Australia. Genome size was correlated with mean annual temperature (MAT) across all ranges, linking genomic traits to environmental variables.
- Genome size evolution during invasion can be rapid, adaptive, and range-specific, with TE-driven genome expansion emerging as a potential genomic response to the demographic and environmental pressures accompanying colonization of novel environments.

## Introduction

Genome size varies enormously across eukaryotes and is largely decoupled from gene number, reflecting extensive variation in non-coding and repetitive DNA (Elliott et al, 2015; Slijepcevic, 2018; Gregory, 2005). In angiosperms alone, genome size varies 2,400-fold (Pellicer et al, 2010), from the miniature genome of *Genlisea tuberosa* (1C = 63 Mbp; Zedek et al, 2024) to *Paris japonica* (1 C=148.89 Gbp), the largest eukaryotic genome currently known (Fernández et al, 2024). Once viewed as a passive consequence of duplications, deletions, and transposable element (TE) activity (Gregory et al, 2004; Adams et al, 2023), genome size is increasingly recognised as a form of structural variation with potential consequence for organismal performance and adaptation (Suda et al, 2015; Bilinski et al, 2018; Blommaert, 2020). In plants, genome size has been linked to cell division rates, development, and phenology, suggesting that it may mediate responses to environmental gradients (Blommaert, 2020; Robinson et al, 2018). At the same time, genome size can evolve rapidly through non-adaptive processes, including TE proliferation and genetic drift, particularly under demographic disequilibrium (Smith et al, 2016; Dazenière et al, 2022). Biological invasions, which combine strong demographic perturbations with exposure to novel environments, therefore provide powerful natural experiments for disentangling adaptive and non-adaptive drivers of genome size evolution. Here, we quantify genome size and repetitive element variation across native and invasive populations of the globally invasive common ragweed (*Ambrosia artemisiifolia* L.), integrating genomic, climatic, and phenotypic data to test how genome size evolves during rapid range expansion across broad climatic gradients.

Two non-mutually exclusive frameworks have been proposed to explain genome size variation within species. In one framework, genome size influences traits like cell cycle duration, growth rates, and phenology, such that selection acting on these traits indirectly shapes genome size along environmental gradients (Blommaert, 2020; Robinson et al, 2018). In the other, genome size evolves indirectly through the accumulation of TE insertions that are typically neutral or deleterious and persist under demographic regimes characterised by strong genetic drift and inefficient purifying selection, rather than through selection on genome size-associated traits (Smith et al, 2016; Arkhipova et al, 2018; Dazenière et al, 2022; Marino et al, 2024). Environmental stressors associated with habitat shifts may further promote TE activity by disrupting epigenetic silencing mechanisms, while associated demographic bottlenecks may reduce the efficacy of purifying selection, facilitating the persistence of TE insertions (Fedoroff, 2012; Lisch, 2013; Makarevitch et al, 2015; Lockton et al, 2008). These frameworks generate contrasting expectations. Genome size evolution shaped by divergent selection imposed by local environments should produce population differentiation exceeding neutral genetic divergence and correlations with the environmental gradient, whereas genome size evolution driven primarily by drift and reduced efficacy of purifying selection is expected to yield region-specific patterns reflecting invasion history.

Genome size is particularly relevant for invasion biology because it is linked to life-history characteristics that determine colonization success, establishment, and persistence in novel environments (Suda et al, 2015; Cang et al, 2024). Global surveys show that invasive plant species are disproportionately drawn from lineages with smaller genomes (Suda et al, 2015; Guo et al, 2024). Patterns of genome size variation among invasive plant species have been interpreted in the context of the Large Genome Constraint (LGC) hypothesis (Bennett, 1987; Knight et al, 2005), which proposes that larger genomes impose a minimum threshold on cell cycle duration and generation time. This produces a triangular relationship between genome size and developmental rate: species with large genomes are excluded from fast-growing strategies entirely, while species with small genomes can occupy both fast and slow ends of the spectrum (Šímová & Herben, 2012; Bhadra et al, 2023; Bures et al, 2024). However, genome size-invasion relationships are not uniformly directional: larger genomes have also been associated with slow-growing, resource-dominant strategies in invasive plants that establish through increased competitiveness rather than rapid colonisation (Grime, 1982,1998; Guo et al, 2024). These patterns are largely based on interspecific comparisons and therefore provide limited insight into how genome size evolves within species during invasion.

Evidence is growing that intraspecific genome size variation is not stochastic but can have significant ecological relevance, potentially shaped by environmental filtering and natural selection across environmental gradients (Kalendar et al, 2000; Smarda & Bures, 2010; Smarda et al, 2010; Bilinski et al, 2018). Intraspecific genome size variation may reflect a complex interplay between local selection on life-history traits and non-adaptive processes associated with invasion, including founder events and population bottlenecks that can reduce the efficacy of purifying selection and facilitate TE accumulation (Stapley et al, 2015; Hodgins et al, 2025; Merel et al, 2021; Schrader et al, 2014). The contrasting predictions based on cross-species patterns and within-species evolutionary dynamics highlights the need for intraspecific tests of genome size change during invasion, including explicit tests for selection on genome size divergence and triangular constraints predicted by the LGC hypothesis.

Common ragweed (Asteraceae) is an annual, wind-pollinated weed, native to North America that has rapidly expanded across Europe, Australia, and other regions since the nineteenth century (Knolmajer et al, 2024). Genomic studies indicate that the European invasion proceeded through multiple introductions, primarily from the West and Mideast North American genetic clusters, with little evidence for a reduction in effective population size (Bieker et al, 2022; Battlay et al, 2025a; Battlay et al, 2025b). In contrast, the more recent Australian invasion, first recorded in the early twentieth century, was primarily derived from South and Mideast North American genetic clusters and experienced a substantial reduction in effective population size consistent with a strong founder event, with a secondary restricted introduction from New England (Battlay et al, 2025a; Battlay et al, 2025b). Ragweed’s invasion has exposed populations to broad climatic gradients and substantial demographic perturbations associated with range expansion, while retaining considerable standing genetic variation (Bieker et al, 2022; Battlay et al, 2025a). Remarkably, latitudinal clines in several important traits such as flowering time and plant size, initially observed in the native range, have evolved in Europe and Australia, indicating rapid and parallel post-invasion adaptation (van Boheemen et al, 2019; McGoey et al, 2020). While previous population genomic studies have identified SNPs and structural variants (e.g., inversions, copy number variants) involved in local adaptation in this species (Battlay et al, 2023; Wilson et al, 2025), the role of genome size and TE content in shaping adaptive responses across its global range remains underexplored. Previous studies have reported genome size variation in common ragweed (Kubešová et al, 2010; Bai et al, 2012; Hrabovský et al, 2024b; Battlay et al, 2023), but the evolutionary drivers and ecological consequences of this variation remain unresolved, particularly across the species’ native and invasive ranges.

We use common ragweed introductions as a replicated natural experiment to examine how genome size and repetitive DNA evolve during rapid range expansion across broad climatic gradients. By integrating whole-genome resequencing, flow cytometry-based genome size estimates, climatic data, and life-history traits across native and invasive ranges, we address the following specific questions: (1) How is repetitive DNA, particularly TEs, distributed across the genome, and which components contribute most to genome size variation? (2) How do genome size and TE content vary across the native and introduced ranges, and are these patterns associated with climate? (3) Does genome size divergence among populations exceed neutral genetic divergence, consistent with selection acting on genome architecture rather than drift alone? and (4) does genome size impose a lower boundary constraint on developmental speed (flowering time), as predicted by the LGC hypothesis? Together, these questions allow us to evaluate whether genome size evolution during invasion is more consistent with adaptive responses to novel environments, non-adaptive demographic processes, or their interaction.

## Materials and Methods

### Samples and alignment

The primary haplotype (haplotype 1) of the diploid chromosome-level reference genome of common ragweed, originally sequenced and assembled from a sample collected in Novi Sad, Serbia (45.25°N, 19.91°E) was used as reference (Battlay et al, 2023). To ensure accurate mapping to the nuclear genome, reads were aligned to a refined version of this reference by removing 1) scaffolds shorter than 100 kbp, 2) scaffolds with more than 20% of annotated genes of mitochondrial or chloroplast origin, and 3) scaffolds containing fewer than one annotated gene per megabasepair (Mbp). The mitochondrial and chloroplast genomes were independently assembled (Terzer et al, in prep.) and added back to the reference. The resulting reference comprised two organellar genomes, 18 nuclear chromosomes, and 31 additional nuclear scaffolds with total nuclear genome size of 1.06 Gbp, compared to the unfiltered assembly of 1.1 Gbp.

We used a subset of the whole-genome resequencing data described in Bieker et al (2022) and Battlay et al (2025a). The leaf tissues from 443 individuals were sampled between 2007 and 2019 and sequenced on Illumina platforms to an average depth of ∼7.1X. Sampling spanned the native range (North America; n=179) and invasive ranges in Europe (n=169) and Australia (n=96). Reads were aligned to the filtered reference using the RepAdapt pipeline (https://github.com/RepAdapt/snp_calling_simple) incorporating fastp v.0.20.1 (Chen et al, 2018; https://github.com/OpenGene/fastp) for read trimming, bwa-mem v.0.7.17-r1188 (Li et al, 2013; https://github.com/lh3/bwa) and samtools v.1.16.1 (Li et al, 2009; https://github.com/samtools/samtools) for alignment, Picard Tools v.2.26.3 (Picard Toolkit, 2019; https://github.com/broadinstitute/picard) for duplicate removal and GATK v.3.8 (McKenna et al, 2010; https://github.com/broadgsa/gatk) for local realignment of reads around indels. Because we were interested in reads mapping to repetitive regions of the genome, we set the samtools parameter from `-q 10` to `-q 0` to relax the mapping quality filter and retain more mapped reads.

### Genome size estimation (sequence-based method)

We used average read-depth to estimate an effective haploid genome size (1C, Gbp) for each sample (Desvillechabrol et al, 2016; Pflug et al, 2020; Pfenninger et al, 2022; Natarajan et al, 2025). Although sequencing reads originated from diploid individuals, these were mapped to a primary haplotype assembly, allowing us to infer the size corresponding to a single, haploid set of chromosomes. First, the mean genome-wide coverage depth was calculated from aligned BAM files using the `samtools depth -aa` command in samtools v.1.16.1, with organelle scaffolds excluded. We scaled genome-wide depth by the mean depth over a set of putatively single-copy loci from the Benchmarking Universal Single-Copy Orthologs (BUSCO) eukaryota *odb10* dataset (Simão et al, 2015). Nevertheless, our reference genome contained 72 apparent duplications of BUSCO genes (Battlay et al, 2023), which we removed, leaving 183 genes for this analysis. The mean gene depth across these genes was calculated using `samtools depth` on a BED file of their genomic coordinates generated from the genome’s gene annotation, documented in Battlay et al, 2023. Genome size for each sample was then estimated by dividing the mean genome-wide depth by the mean BUSCO depth, reflecting coverage over single-copy regions. We used a coverage-based adaptation of the Lander-Waterman equation (Lander et al, 1988). In our case, genome size (G) was calculated as the ratio of the mean whole genome depth 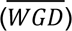 to the mean BUSCO depth 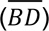:

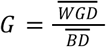

We also assessed several sequencing quality metrics including sequencing coverage depth, insert size, read length, mapping quality thresholds, callable base depth, BUSCO gene depth variance, and PCR clonality, for their potential effect on genome size estimation (Suppl. Material SM1; Suppl. Fig. S3; S4). We then removed samples with genome sizes more than three standard deviations from the mean genome size (Suppl. Fig. S1), resulting in 439 samples retained for downstream analyses (Suppl. Table S9).

### Flow cytometry

Sequence-based genome size estimates were independently validated using flow cytometry (n = 40) with *Manihot esculenta* as an internal control. Detailed protocols for sample preparation, calibration, and fluorescence analysis are provided in the Suppl. Material (SM2).

### TE annotation

TEs were annotated on the primary haplotype of the filtered reference genome (Battlay et al, 2023) using the EDTA (Extensive *de-novo* TE Annotator) pipeline (Ou et al, 2019; https://github.com/oushujun/EDTA), with the `-anno` function and default parameters. These elements were then processed to generate a non-redundant, curated TE library. This step clusters TEs into families while reducing redundancy, based on a modified 80-80-80 Wicker rule (Wicker et al, 2007), which groups elements that share ≥80% identity over ≥80% of their length and belong to the same superfamily. The curated TE library was then used to mask the genome using RepeatMasker v.4.1.1 (Smit, AFA, Hubley, R & Green, P. RepeatMasker Open-4.0.2013-2015; http://repeatmasker.org). Subsequently, the masked genome was re-annotated with the same TE library using EDTA’s `-anno` function to produce a comprehensive TE annotation. For downstream analyses, we used the split annotation BED file, which retains only non-overlapping TE annotations, ensuring that any particular genomic region is assigned to only a single TE feature.

### Ribosomal RNA annotation

RNAmmer version 1.2 (Lagesen et al, 2007) was used to annotate eukaryotic ribosomal RNA (rRNA) on the reference genome, including subunits 28S, 18S, and 8S. RNAmmer annotates rRNA using hidden Markov models (HMMs) (Rabiner et al, 1986), capturing structural features from multiple alignments of rRNA database sequences.

### Non-overlapping TE annotation and abundance estimation

A non-overlapping annotation file was generated by integrating annotations for TEs, rRNA, simple repeats, and low-complexity regions, with the following prioritization: rRNA > known TEs > simple repeats > low-complexity regions (adapted from Kreiner et al, 2023). Since EDTA did not identify simple and low-complexity repeats, their corresponding annotation BED files were obtained from the RepeatModeler output generated in our previous study (Battlay et al, 2023). Sequences shorter than or equal to 20 bp were excluded to minimize false positives. The resulting curated annotation file was used for downstream analyses.

To estimate TE abundance, we employed a read-depth-based approach analogous to that used for genome size estimation. Specifically, the abundance of each TE family was calculated as the mean depth of reads mapping to each TE annotation, normalized by the mean read depth over BUSCO genes. This provides an estimate of the relative copy number for each TE family, expressed as:

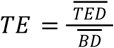

where (*TE*) is the abundance/copy number of a TE family, 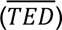 is the mean read depth over TE annotations and 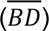 is the mean read depth over single-copy BUSCO genes. We then multiplied this normalized copy number by the length of each TE family (Suppl. Table S1) to calculate the per-individual total repeat abundance in base pairs (bp). We also assessed the effect of sequencing coverage depth on TE or rRNA abundance estimates by regressing whole-genome average depth against the estimated abundance of each TE or rRNA family across all samples (Suppl. Material SM1; Suppl. Fig. S8).

The size and quantity of TEs were directly visualised using non-overlapping annotations based on counts and base pair ranges for each superfamily. To determine the distance to the nearest gene, bedtools closest 2.31.0 (Quinlan et al, 2010; https://github.com/arq5x/bedtools2/releases) was employed, identifying the closest gene to each repeat element and reporting the distance using the `-d` option. The fraction of each 1 Mbp genomic window occupied by different types of repeats was computed using the `genomicDensity()` function from the R package circlize v0.4.16 (Gu et al, 2021; https://jokergoo.github.io/circlize_book/book/). Variance and coefficient of variation of repeat abundance were calculated in R.

### Comparison of trait versus genetic differentiation

To understand which evolutionary forces drive variation in genome size among populations, we performed a *Q_ST_–F_ST_* comparison as the primary analysis (Spitze, 1993; Prout & Barker, 1993; Whitlock, 2008). This analysis utilized whole-genome resequencing data from 289 individuals (representing all populations with n≥2) based on the variant call format (VCF) provided by Battlay et al. (2025a). Two complementary approaches, a *P_ST_–F_ST_* comparison (Sæther et al, 2007; Whitlock, 2008), which assesses phenotypic differentiation relative to neutral expectations, and *Q_PC_* analysis (Josephs et al, 2019), which tests for adaptive divergence along axes of genetic relatedness without requiring predefined populations, are described in Supplementary Material SM4.

To estimate *Q_ST_*, we first constructed a genomic relatedness matrix (GRM) from 50,000 LD-pruned (r^2^ = 0.5), MAF-filtered (≥0.01) SNPs excluding genic regions and known haploblocks, using the VanRaden (2008) method implemented in R\AGHmatrix (Amadeu et al, 2023). We then modelled genome size using a Bayesian animal model implemented in R\MCMCglmm (Hadfield, 2010), with the GRM used to parameterize the structure of the covariance of the genotypic effect. Individual genomic breeding values were extracted from each of the posterior distributions from 1,000 MCMC samples and used to calculate a posterior distribution of *Q_ST_* directly, without collapsing to a single point estimate. Within each geographical range (North America, Europe, Australia), breeding values were partitioned by population and *Q_ST_* was calculated as (Spitze, 1993; Lande, 1992; Whitlock, 2008):

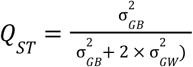

Where 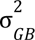is the additive genetic variance among-population and 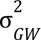 the additive genetic variance within-population. We report the mean of the *Q_ST_* posterior distribution and 95% credible interval. Full MCMC settings, convergence diagnostics, and heritability estimation details are provided in Suppl. Material SM4a.

For each range, observed *Q_ST_* was compared to a null distribution of *F_ST_* values generated from 10,000 putatively neutral, LD-pruned sites, randomly sampled from non-genic and non-structural variant regions of the VCF mentioned above; these SNPs were explicitly excluded from those used in GRM construction, ensuring independence between the neutral reference distribution and the markers used to estimate quantitative genetic parameters. For each range, SNPs were filtered for minor allele frequency ≥0.05 and LD-pruned (50-kbp window, 5-SNP step, *r*² = 0.5) using PLINK v1.9. Overlapping SNPs across all ranges were then identified using `bcftools isec` (Danecek et al, 2021; http://github.com/samtools/bcftools), and 10,000 SNPs were randomly selected for the calculation of pairwise Weir and Cockerham *F_ST_* for each range in `vcftools` (Weir et al, 1984; https://github.com/vcftools/vcftools). *Q_ST_* values exceeding the 99^th^ percentile of the neutral *F_ST_* distribution were interpreted as evidence of trait divergence exceeding neutral expectations, consistent with divergent selection. Values falling between the 1^st^ and 99^th^ percentiles were considered consistent with genetic drift alone. Values in the lower 1^st^ percentile of the neutral *F_ST_* distribution indicate reduced among-population divergence relative to neutral expectations and may reflect stabilizing or parallel selection, high within-population variance, or limited statistical power, and were therefore interpreted cautiously (Whitlock et al, 2009).

### Principal component analysis

We performed principal component analysis (PCA) on genome wide SNPs (VCF provided in Battlay et al, 2025a) to capture neutral population structure and account for its effects in the analyses of genome size and life history traits. Separate PCAs were conducted for the overall analysis (439 samples) and the phenotypic analysis (221 samples). The detailed method on PCA analyses is provided in Suppl. Material (SM3).

### Polygenic value estimation of flowering time

Polygenic values (PGV) for flowering time were estimated using flowering time-associated loci and effect sizes derived from a previous genome-wide association study (GWAS; Battlay et al, 2025a), with detailed methodology provided in Suppl. Material (SM5).

### Statistical analysis

We conducted all statistical analyses in R (v4.4.2). Most figures utilised the ggplot2 package (Wickham et al, 2016) for visualisation.

To explore differences in genome size and TE abundance across the native and two invasive ranges, we fitted linear-mixed models (LMMs), implemented in the R\lme4 package v1.1-36 (Bates et al, 2015; https://github.com/lme4/lme4):

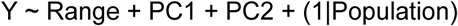

where Y is the trait (genome size or abundance of different TE families). Range was included as a fixed effect, and Population was included as a random intercept. Principal components (PC1 and PC2) derived from genome-wide SNP data were included as fixed effects to account for background population structure in analyses where SNP data were available. As a result, these terms were not included for analyses based solely on flow-cytometry–based genome size measurements. Model assumptions were evaluated by visual inspection of residual plots. The significance of fixed effects was assessed using Type III Wald F-tests implemented with the car package (Fox & Weisberg, 2018) applied to linear mixed-effects models fitted with `lme4`. Given a significant association with geographic range, post-hoc tests on estimated marginal means (EMMs) were conducted using the `emmeans` package (v1.10.7; Lenth et al, 2023; https://cran.r-project.org/package=emmeans) to examine pairwise differences. EMMs were calculated with the `emmeans()` function, and pairwise comparisons were performed using the `pairs()` function, applying Tukey Honest Significant Difference (HSD) test.

To distinguish the roles of ancestry (putative source population) from that of post-invasion changes in genome size, we compared the genome size of invasive range populations against each of the four spatio-genetic clusters in the native range. We used the same LMM framework, ANOVA, and post-hoc pairwise comparisons described above.

We further investigated the drivers of genome size, specifically the contributions of TE content and environment, and explored its relationship with phenotypic traits. Given the notably higher genome size and TE content in Australian populations, we tested the contribution of specific TE families and rRNA abundance to intraspecific genome size variation. To do so, we fitted linear mixed-effects models using the following R model formula:

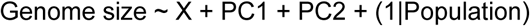

where Genome size is the response variable, X is the abundance of different TE families and Population represents a random intercept for each population. PC1 and PC2 were included as fixed effects as previously described.

To understand the relationship between climate and genome size, we focused on WorldClim (Fick & Hijmans, 2017) variable mean annual temperature (MAT) because it has been linked with genome size variation in previous studies (Guo et al, 2024; Meyerson et al, 2024; Hrabovský et al, 2024b). Bioclimatic variables at our sampling locations were highly collinear, with the first principal component (PC1) explaining approximately 48% of the total climatic variance and showing a strong correlation with MAT (r = 0.84; Suppl. Material SM6, Suppl. Fig. S13). Accordingly, MAT was used as a biologically interpretable proxy for the dominant climatic gradient. We fitted linear mixed-effects models using the following R model formulas:

*Base model*,

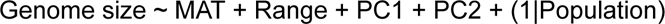

*Interaction model*,

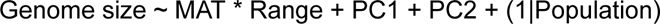

where MAT, Range, PC1, and PC2 were treated as fixed effects and Population was included as a random intercept. The base model included geographic range as a fixed effect to control for range-specific differences, while the interaction model tested whether the effect of MAT on genome size differed among ranges.

To test the LGC hypothesis, that larger genomes impose a lower boundary on developmental speed rather than shifting the mean, we first fitted a linear mixed model of flowering time on genome size, which revealed no significant effect on the mean. To test boundary constraints at tails of the distribution, we also fitted Linear Quantile Mixed Models (LQMM; `lqmm` R package) at five quantiles (*τ* = 0.10, 0.25, 0.50, 0.75, 0.90; Suppl. Material SM7).

## Results

### Transposable element landscape and abundance

TEs constituted 68.5% of the common ragweed genome, consistent with prior estimates (Battlay et al, 2023). The TE landscape was dominated by Long Terminal Repeat (LTR) retrotransposons (42.3%; 448 Mbp), primarily comprising LTR/Copia (17.5%), LTR/Gypsy (12.7%), and LTR/Unknown (12.1%) elements (Fig. 1a, panel 1). Other abundant families included Helitrons (16.4%) and TIR transposons such as TIR/Mutator (4.8%), TIR/hAT (2.8%), and TIR/CACTA (1.4%), with less abundant families detailed in Suppl. Table S1 (Fig. 1a, panel 1). The genomic distribution of TEs revealed distinct family-specific trends with respect to distance to genes, copy number, and element size (Fig. 1a, panels 2–4; Suppl. Table S1). TIR/Tc1_Mariner elements were located closest to genes (mean distance = 15,620 ± 458 bp) while TIR/Mutator elements were the most distal (mean distance = 192,314 ± 861 bp), suggesting differential impacts on gene regulation (Fig. 1a, panels 2; Suppl. Table S1). In terms of copy number, Helitrons were the most abundant (473,277 insertions) while TIR/Tc1_Mariner were the least frequent (4,859 copies; Fig. 1a, panel 3; Suppl. Table S1). LTR/Gypsy and LTR/Copia were the longest elements on average (1,277 ± 6 bp and 1,197 ± 5 bp respectively), while TIR/Tc1_Mariner were the shortest (265 ± 4 bp; Fig. 1a, panel 4; Suppl. Table S1). TE density was markedly heterogeneous across chromosomes, higher in pericentromeric and central regions (Fig. 1b). Ribosomal RNA elements showed local enrichment on chromosomes 1, 5, and 15, with a notable expansion on chromosome 5, and slightly depleted from regions with high TE density (Spearman ρ = −0.11, P = 0.0002; Fig. 1c). TE-rich regions were consistently gene-poor (Spearman ρ = −0.78, P < 2.2 x 10^-16^; Fig. 1d).

**Fig. 1:**
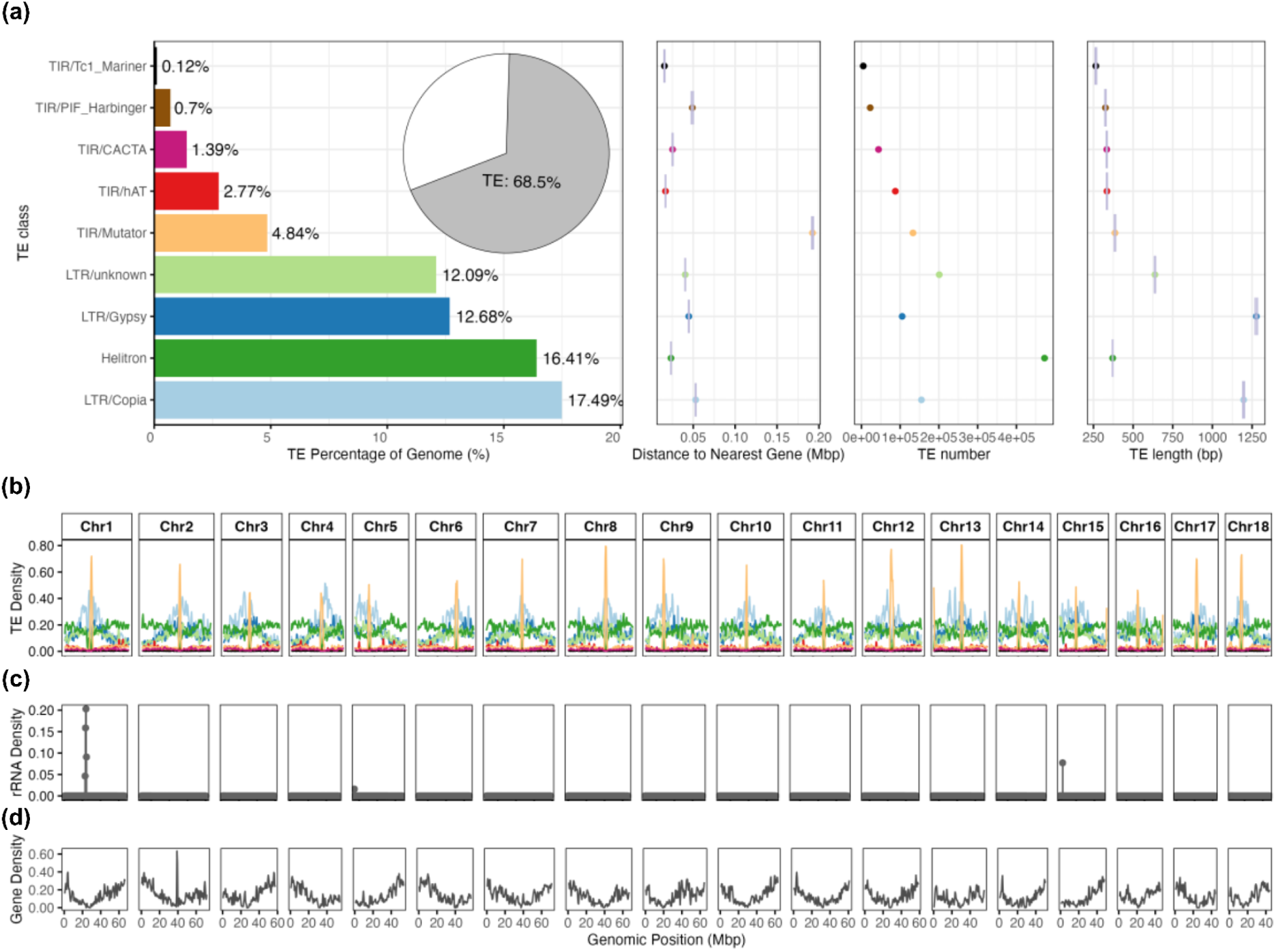
Variation in TE composition across common ragweed reference genome. (a) Proportion of the common ragweed reference genome made up by various TE classes, highlighting differences in distance to genes, number, and size across the genome. Error bars represent standard errors. (b) Distribution of different TE density across the 18 chromosomes (top; colour codes from (a), calculated using sliding 1Mbp windows. (c) & (d) Distribution of rRNA and gene density across the genome respectively, calculated using sliding 100-kbp windows and plotted alongside the corresponding TE density windows.

### Geographic variation in genome size

From our sequence-based estimates, the haploid genome size of common ragweed ranged from 0.88 to 1.23 Gbp, with a mean ± SE of 1.01 ± 0.0034 Gbp (Fig. 2a). Common ragweed exhibited a pronounced regional difference in genome size across its native and its two introduced ranges (linear mixed-effects model: F_2,138.8_ = 25.39, P = 4.018 x 10^-10^, R^2^ = 0.61; Table 1). Post-hoc tests confirmed that Australian genomes were significantly larger than European and North American genomes (Tukey’s HSD; diff_AUSvsEUR = 100 Mbp, P < 0.0001; diff_AUSvsNAM = 120 Mbp, P < 0.0001; estimated marginal means ± SE: AUS = 1.1 ± 0.015 Gbp, EUR = 1.00 ± 0.007 Gbp, NAM = 0.98 ± 0.006 Gbp; Fig. 2b, left panel). Conversely, genome sizes between Europe and North America did not differ significantly (diff_EURvsNAM = 20 Mbp, P = 0.26) (Fig. 2b).

**Fig. 2:**
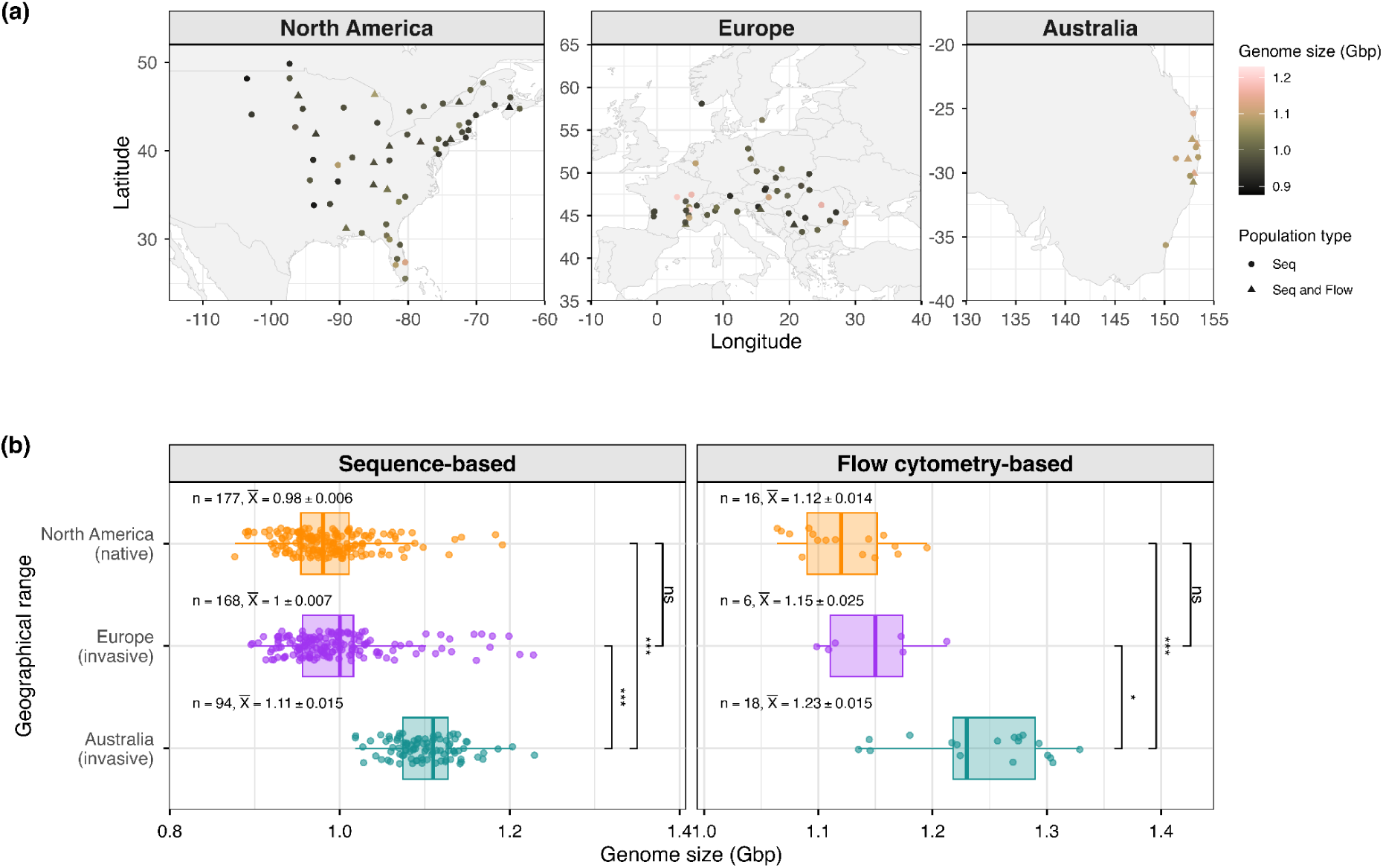
Pattern of genome size variation across native and invasive ranges. a) Geographical locations of populations used in this study and their genome size estimates. Circles represent populations with sequence-based estimates, and triangles indicate populations with both sequence-and flow-based estimates. b) Genome size variation among geographical ranges; native (North America) and invasive (Europe and Australia) from two independent analyses: genomic read depth-based (left panel) flow cytometry-based (right panel). Above each box, we report sample size (n) and estimated marginal mean (*X̄*) ± standard error. Significance codes: *** < 0.001, ** < 0.01, * < 0.05, ns = not significant.

**Table 1.** Linear mixed model for genome size with range, PC1, PC2 as fixed effects and population as a random intercept.

| Fixed effect | F | Df | Df.res | P |
| --- | --- | --- | --- | --- |
| Range | 25.3871 | 2 | 138.794 | 4.018 x 10<sup>-10</sup> |
| PC1 | 0.0027 | 1 | 210.657 | 0.9590 |
| PC2 | 2.5931 | 1 | 240.659 | 0.1086 |

### Validation of sequence-based genome size variation with flow cytometry

Sequencing-based estimates of haploid genome size were independently validated using flow cytometry, with calibration details provided in Suppl. Material SM2 (also see Suppl. Fig. S5). The resulting haploid genome size estimates for common ragweed were approximately normally distributed (Suppl. Fig. S6), with a mean of 1.18 ± 0.01 Gbp, ranging from 1.06 to 1.33 Gbp (Suppl. Table S2). At the population level, flow cytometry and sequencing-based genome size estimates were highly correlated (absolute Pearson’s r = 0.938, Suppl. Fig. S7). Flow cytometry revealed significant differences in genome size across geographic ranges (F_2,22.627_ = 14.62, P = 8.4 x 10^-5^), consistent with the sequencing-based findings (Fig. 2b, right panel).

### Genome size difference between the invasive ranges and their corresponding native sources populations

As previously described, genomic analysis suggested that the two invasion ranges were introduced from distinct native source clusters. We therefore compared genome size of invasive ranges against their likely native source populations. Significantly larger genome sizes were observed in the Australian invasive range compared to its putative native sources (North American South and Mid-East) and other native North American populations (Fig. 3, Suppl. Table S3). In contrast, the European invasive range showed overall similar genome sizes to native North American populations and their putative native sources. These patterns in genome size mirrored similar observations for the abundance of most TE and rRNA families (Suppl. Fig. S9).

**Fig. 3:**
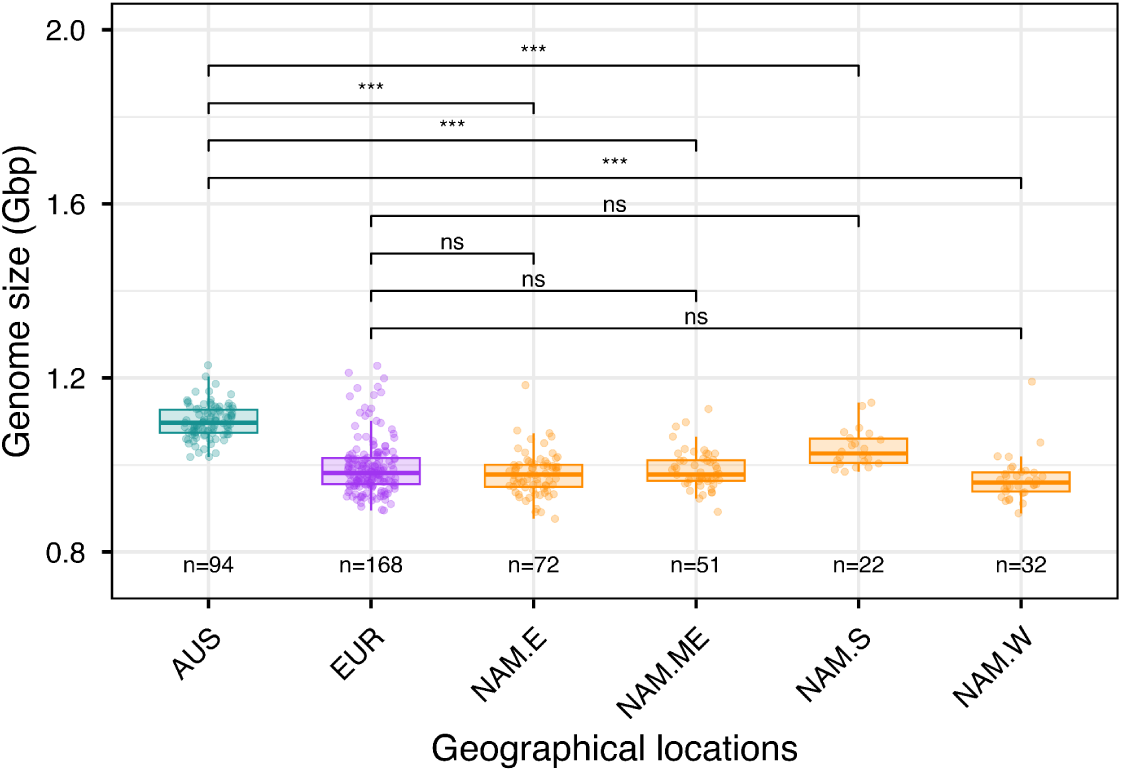
Genome size differences between invasive ranges and their native populations. Haploid genome size (Gbp) estimates using sequence data for Australia (AUS), Europe (EUR), North America spatio-genetic clusters mid-east (NAM.ME), south (NAM.S), west (NAM.W) and east (NAM.E). Sample sizes (n) are provided for each cluster. Asterisks denote statistical significance from pairwise comparisons (e.g., \**P* < 0.05, \*\**P* < 0.01, \*\*\**P* < 0.001), and ‘ns’ indicates non-significant differences.

### Geographical Pattern of Class-Specific TE Abundance

To assess the potential for TE and rRNA variation to contribute to intraspecific adaptation in common ragweed, we quantified the abundance of TE and rRNA families across ranges. The mean ± standard error length of genome represented by each TE family across individuals mirrored the rank order observed in our reference genome: LTR/Copia (208 ± 1.05 Mbp), Helitron (159 ± 0.43 Mbp), LTR/Gypsy (156 ± 1.06 Mbp), LTR/unknown (133 ± 0.51 Mbp), TIR/Mutator (55.6 ± 0.39 Mbp), TIR/hAT (38.2 ± 0.17 Mbp), TIR/CACTA (13.1 ± 0.03 Mbp), TIR/PIF_Harbinger (6.8 ± 0.02 Mbp), Simple Repeats (5.27 ± 0.03 Mbp), TIR/Tc1_Mariner (1.25 ± 0.01 Mbp), and Low Complexity (0.88 ± 0.01 Mbp) (Suppl. Fig. S11). Despite this compositional consistency, Australian samples exhibited higher abundances of TEs across most families compared to European and North American samples (Fig. 4a, Suppl. Fig. S12). Linear mixed models confirmed a significant effect of geographic range on most TE and rRNA families (excluding TIR/hAT and TIR/PIF_Harbinger; P < 0.05; Suppl. Table S4). Pairwise comparisons revealed that most TE and rRNA families, particularly LTR/Copia, LTR/Gypsy, and TIR/Mutator, were significantly more abundant in Australia than in either Europe or North America (Suppl. Table S4). Among all repeat types, rRNA families (8S, 18S, 28S) exhibited the highest coefficient of variation (CV), likely reflecting their confinement to a few clustered loci where small absolute changes yielded large proportional variance (Fig. 4b). By contrast, highly abundant TEs such as LTR/Copia, LTR/unknown and Helitron showed lower relative variance but substantial absolute copy-number variation, suggesting these elements contribute notably to genomic differences among individuals.

**Fig. 4:**
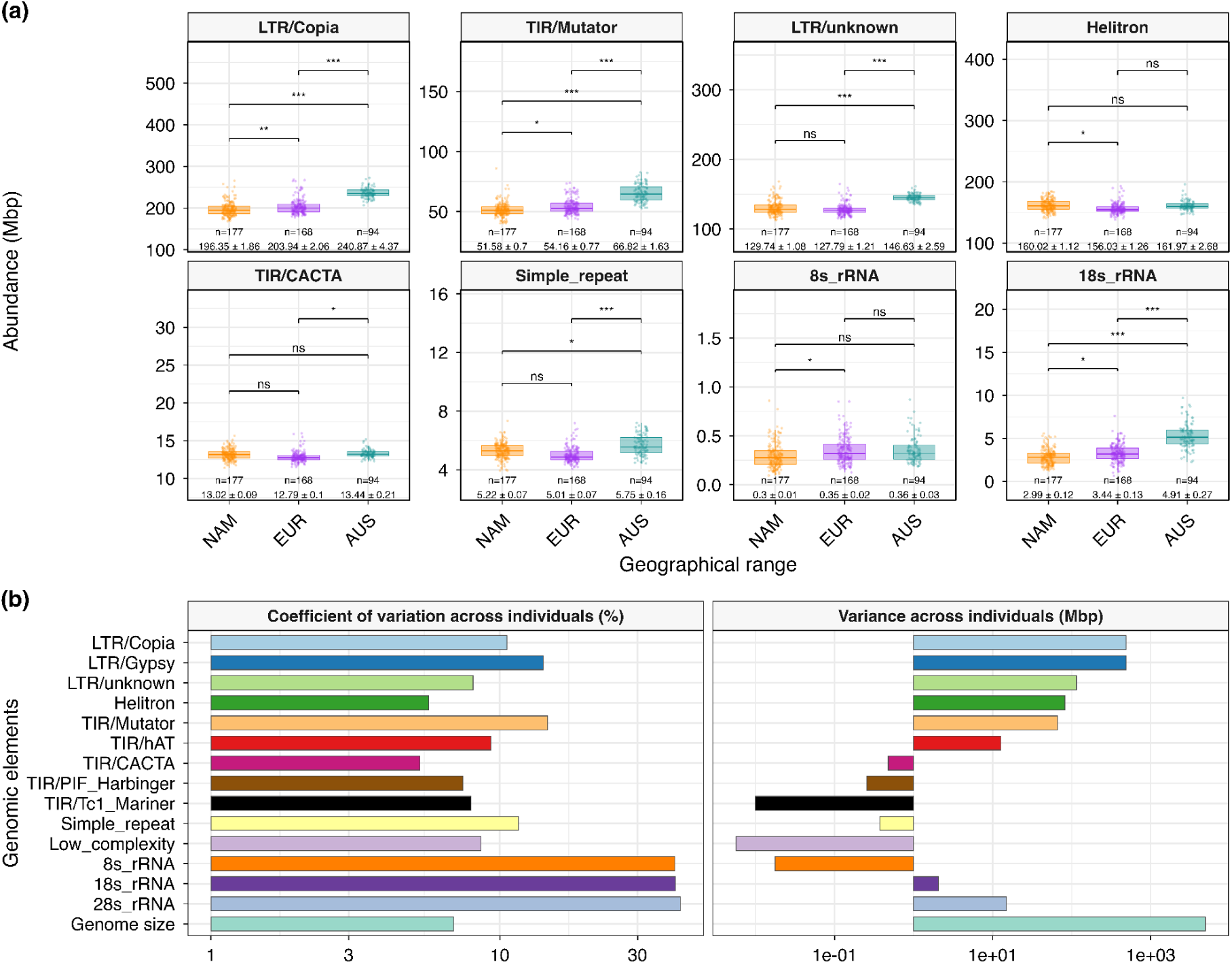
TE and rRNA abundance variation across individuals. a) Variation in TEs and rRNAs abundance among individuals from different geographical ranges; North America (NAM), Europe (EUR) and Australia (AUS). b) Measures of variability in TE and rRNA abundance: left, coefficient of variation (CV); right, Variance (V, calculated on log-transformed values). Below each box, we report sample size (n) and estimated marginal mean (*X̄*) ± standard error. Asterisks denote statistical significance for pairwise comparisons (e.g., \**P* < 0.05, \*\**P* < 0.01, \*\*\**P* < 0.001), and ‘ns’ indicates non-significant differences.

### TE and rRNA abundance significantly contribute to genome size variation

The abundance of specific TEs and rRNA predicted genome size well (Suppl. Table S5). Our linear mixed models revealed strong and significant associations between genome size and TEs abundance, particularly LTR and TIR families (Suppl. Table S5). Furthermore, consistent with TE abundance, rRNA subunit abundance, except 18S-rRNA, was likewise significantly correlated with genome size (Suppl. Table S5).

### Genome size selection analysis

To test whether divergent selection shaped genome-size variation, we performed *Q_ST_-F_ST_* comparisons. We first confirmed that genome size variation has a strong additive genetic basis with a posterior mean SNP-based heritability of ^2^ = 0.886 (95% CI: 0.772–0.953).

In North America and Europe, additive genetic differentiation (*Q_ST_*) significantly exceeded neutral expectations. In North America, the posterior mean *Q_ST_* (0.362) far exceeded the 99^th^ percentile of the neutral *F_ST_*distribution (0.26; Fig. 5a). Similarly, in Europe, *Q_ST_* (0.456) exceeded the 99^th^ percentile of the neutral *F_ST_* distribution (0.28; Fig. 5b), providing strong evidence that divergent selection is driving genome size differences among populations in both ranges. In contrast, populations in Australia exhibited almost no differentiation; the posterior mean *Q_ST_*was 0.035, falling well below the upper neutral *F_ST_* thresholds (0.27; Fig. 5c), providing no evidence that divergent selection has contributed to genome size differentiation within this range. Although *Q_ST_* was low, *Q_ST_* also exceeded the 1^st^ percentile *F_ST_* (0.0009), precluding inference of stabilising selection.

**Fig. 5:**
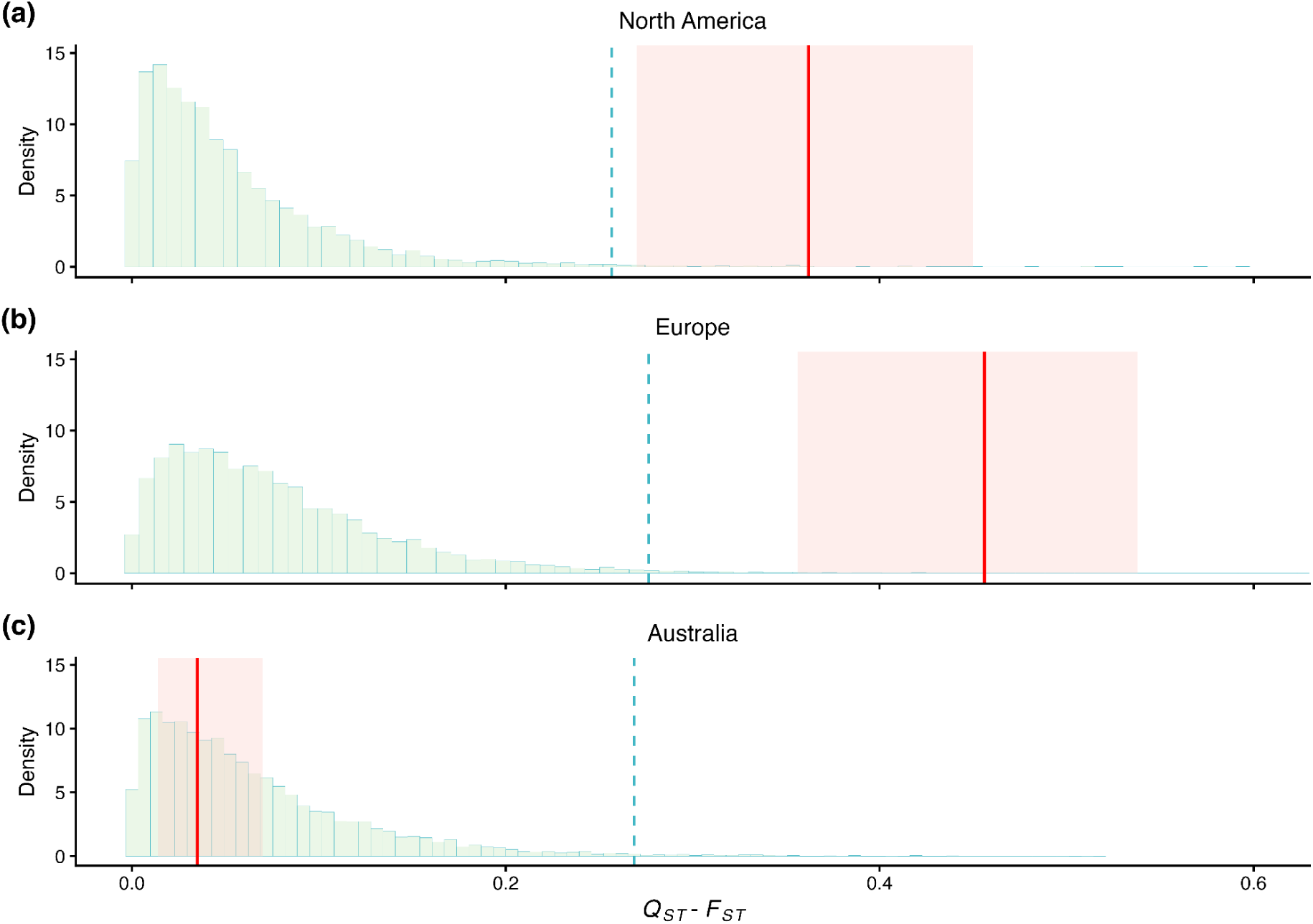
Quantitative genetic differentiation comparison of genome size differentiation across geographic ranges. Quantitative genetic differentiation (*Q_ST_*) for genome size derived from genomic breeding values. The background green histograms show the null distribution of neutral *F_ST_* values, with the teal dashed line indicating its 99^th^ percentile. The shaded light red boxes represent the 95% credible interval of the *Q_ST_* posterior distribution, and the solid red lines represent the posterior *Q_ST_* mean. Posterior mean *Q_ST_* values exceeding the 99^th^ percentile of the neutral *F_ST_* distribution indicate divergent selection on genome size.

These findings were further corroborated by two complementary approaches: *P_ST_–F_ST_* comparison and *Q_PC_* analysis, both of which yielded consistent conclusions across all three ranges (Suppl. Material SM4). Together, these results provide robust evidence for the divergence of genome size during colonisation of North America and Europe, while variation in Australia remains inconsistent with divergent selection.

### Mean annual temperature predicts genome size

Our models revealed strong among-range differences in genome size and support for an overall association with temperature. In the main-effects model, both range and MAT significantly predicted genome size (range: F_2,129.73_ = 18.79, P = 6.81 x 10^-8^; MAT: F_1,144.86_ = 8.2482, P = 0.0047; Suppl. Table S6), with warmer climates associated with larger genomes across ranges. Consistent with this, Australian populations, occurring in warm environments, had the largest genomes on average, whereas European populations, sampled from cooler climates, had the smallest. The interaction between MAT and range was not significant (F_2,83.09_ = 0.38, P = 0.68; Suppl. Table S6), indicating that the positive association between temperature and genome size did not differ detectably among ranges.

### Genome size does not act as a lower-boundary constraint on phenology

LQMM analysis across all ranges and within individual ranges revealed no significant associations at lower quantiles (*τ* = 0.10-0.25; all P > 0.34), inconsistent with a lower-boundary constraint on development. While a significant effect was detected at the median (*τ* = 0.50, P = 0.049) across all ranges combined, range-specific models showed no significance in North America or Europe, and signals were restricted to the upper tail (*τ* = 0.90, P = 0.027) in Australia (Suppl. Table S7-8). Given that only two of twenty quantile tests were nominally significant, these results provide little evidence for a consistent effect of genome size on flowering time and do not support a lower-boundary constraint on flowering time.

## Discussion

The remarkable success of invasive species often involves rapid evolutionary adaptations (Prentis et al, 2008; Suda et al, 2015; Wu et al, 2019; Hodgins et al, 2025) and our study of common ragweed genome size evolution reveals a particularly intriguing case. We found that Australian invasive populations possess significantly larger genomes compared to both their native North American and invasive European counterparts, a difference largely attributable to the increased abundance of TEs. *Q_ST_–F_ST_* comparisons with the support from *P_ST_–F_ST_* and *Q_PC_* analyses in North America and Europe further revealed patterns consistent with divergent selection on genome size, suggesting that genome size divergence is unlikely to be explained by neutral processes alone. Genome size showed evidence of climate structuring, consistent with a role in climate adaptation. Below, we discuss these results in light of the genetic mechanisms underlying TE landscapes, evaluate our findings in the context of hypotheses linking genome size to invasion success, and examine the relative roles of local selection and demographic history in shaping genome size variation across native and invasive ranges.

### The common ragweed TE landscape: a dynamic interplay of composition, context, and organization

We provide a comprehensive analysis of the TE landscape in common ragweed, revealing the pattern of composition, genomic context, and population-level abundance variation. The common ragweed genome is replete with TEs, dominated by LTR retrotransposons (particularly *Copia* and *Gypsy*), consistent with patterns observed in other plant genomes (e.g., *Arabidopsis* [Lian et al, 2024], maize [Chen et al, 2023], sunflower [Badouin et al, 2017], and *Amaranthus* species [Kreiner et al, 2023]). The non-random distribution of TEs, particularly the enrichment of LTR elements in chromosome centers that are relatively gene poor, is likely a consequence of these regions serving as ‘safe havens’ where selection against TE insertions is less effective due to inherently low recombination rates (Kent et al, 2017; Huang et al, 2025). Furthermore, our analysis reveals a clear non-random genomic distance of TE insertions to genes: elements like TIR/Tc1_Mariner are found nearest to genes, while TIR/Mutator elements are typically located farthest from them (Fig. 1a). This differential localization suggests varying selective pressures depending on their impact on nearby genes and their functional importance (Hirsch et al, 2017; Schrader et al, 2019). Together, these TE-specific patterns of accumulation and retention provide a mechanistic basis for the substantial genome size differences we observe among ranges and reinforce the idea that TE dynamics can contribute to rapid genome evolution during invasion (Schrader et al, 2014; Stapley et al, 2015; Goubert et al, 2017; Dennenmoser et al, 2017; Niu et al, 2019). Future work combining population-level TE polymorphism data and environmental associations will be needed to test whether changes in specific TE families are directly linked to range expansion or climate adaptation.

### Does the large genome constraint hypothesis extend to intraspecific genome size evolution during invasion?

Although intraspecific genome size variation is well documented in flowering plants (Greilhuber, 2005), including in Asteraceae (Suda et al, 2007; Slovák et al, 2009; Dirkse et al, 2014), the ecological and evolutionary drivers of this variation, particularly in invasive species, remain poorly understood. Drivers of genome size include polyploidy, demography and mating system shifts, which can change effective population size (*N_e_*) and thus the efficacy of selection, and selection on cell size or growth rate, which can alter genome size and influence life history traits (Glémin et al, 2007; Lefébure et al, 2017; Jiang et al, 2024). Invasive species tend to have smaller genomes (Suda et al, 2015; Pyšek et al, 2023; Guo et al, 2024), a pattern often interpreted as advantageous through faster cell cycles, earlier germination, and higher growth rates which in turn enable their rapid spread during invasions (Kubešová et al, 2010; Suda et al, 2015; Knight et al, 2005). This negative correlation between genome size and invasiveness has been interpreted in the context of the LGC hypothesis (Knight et al, 2005; Kubešová et al, 2010; Pyšek et al, 2023), which proposes that larger genomes impose physiological and developmental constraints through their effects on cell size, cell cycle duration, and generation time (Bennett, 1972; Cavalier-Smith, 1985), indirectly limiting the rapid growth and reproduction that favour establishment and spread in novel habitats.

By substantially expanding population-level estimates of genome size across native and invasive ranges, our study provides a nearly global perspective on common ragweed genome size. We estimated a haploid genome size of 1.01 ± 0.0034 Gbp (range: 0.88 to 1.23 Gbp), a mean value consistent with previous reports (Bai et al, 2012; Kubešová et al, 2010; Battlay et al, 2023; Hrabovský et al, 2024b), but with an expanded upper range exceeding prior sequencing (1.06 Gbp) and flow cytometry (1.13*–*1.16 Gbp) limits due to the inclusion of larger Australian genomes. In contrast to the relatively subtle differences between the European and North American ranges, Australian populations exhibit uniformly larger genomes relative to both regions, including their putative source genetic clusters (Fig. 3), pointing to a pronounced region-specific expansion in genome size following introduction. These findings suggest that the relationship between genome size and invasion success is more context-dependent than implied by broad interspecific patterns linking small genomes with invasiveness (Pyšek et al, 2018,2023; Suda et al, 2015); our results align with a growing body of evidence indicating that genome size alone is not a consistent predictor of invasiveness. Similar findings have been reported for other successful invaders, such as certain *Acacia* species (Gallagher et al, 2011) and various Cactaceae species (Lopes et al, 2021), where no consistent change in genome size was observed in invasive populations. Instead, the adaptive significance of genome size is highly context-specific and contingent upon the particular selective pressures of the introduced environment or the genomic architecture of adaptive traits, including the potential contribution of TE dynamics to standing genetic variation (Richardson et al, 2006; Potapenko et al, 2025). Notably, our quantile analysis provided little support for the lower-boundary developmental constraint predicted by recent formulations of the LGC hypothesis (Bhadra et al, 2023; Bures et al, 2024). Genome size showed no association with flowering time at lower quantiles, where developmental constraints should be the most apparent, and the only marginal signal was detected at the upper flowering time quantile in Australian populations. Nevertheless, Australian populations possess both the largest genomes and the latest flowering times observed across the species’ range (van Boheemen et al, 2019), suggesting that genome size variation may still be associated with life-history divergence. More broadly, while nucleotypic theory predicts developmental constraints associated with increased DNA content, it also predicts effects on cellular dimensions and physiological function (Šímová & Herben, 2012; Roddy et al, 2020), raising the possibility that genome size variation contributes to ecological divergence through multiple mechanisms.

### Evidence of adaptive genome size variation in common ragweed

Across the native North American range, genome size shows greater among-population differentiation than expected under neutrality (Fig. 5), consistent with divergent selection rather than neutral processes. Moreover, genome size correlates with MAT (Fig. 6) across ranges, consistent with climate-mediated selection contributing to genome size differentiation. Environmentally structured genome-size patterns occur in other systems including maize, where genome size correlates with altitude and temperature and tracks cell size and developmental rate (Díez et al, 2013; Tenaillon et al, 2016). In common ragweed, genome size shows climatic structuring with MAT across all ranges yet no significant association with flowering time in our common garden, indicating that genome size is unlikely to be a major determinant of this life-history trait, in contrast to findings in some other species (e.g., Kreiner et al 2023). Instead, the elevated *Q_ST_ and P_ST_*values, together with significant *Q_PC_* results (Suppl. Material SM4) collectively indicate that selection has shaped genome size variation potentially through pathways beyond flowering time, such as other cellular, physiological, or developmental pathways not captured in our common garden. Together, these results suggest that genome size could be responding to spatially variable selection while remaining largely decoupled from the key adaptive life-history trait of flowering time. We note, however, that the lack of correlation between genome size and flowering time in our common garden experiment may partly reflect the benign, resource-rich conditions under which plants were grown. Phenotypic correlations with genome size may disappear in benign experimental gardens but emerge under physiological stress or intense intraspecific competition typical of natural stands (Xiao et al, 2025; Smarda et al, 2010; Li et al, 2026), suggesting that the adaptive significance of genome size variation may be underestimated in common garden settings.

**Fig. 6:**
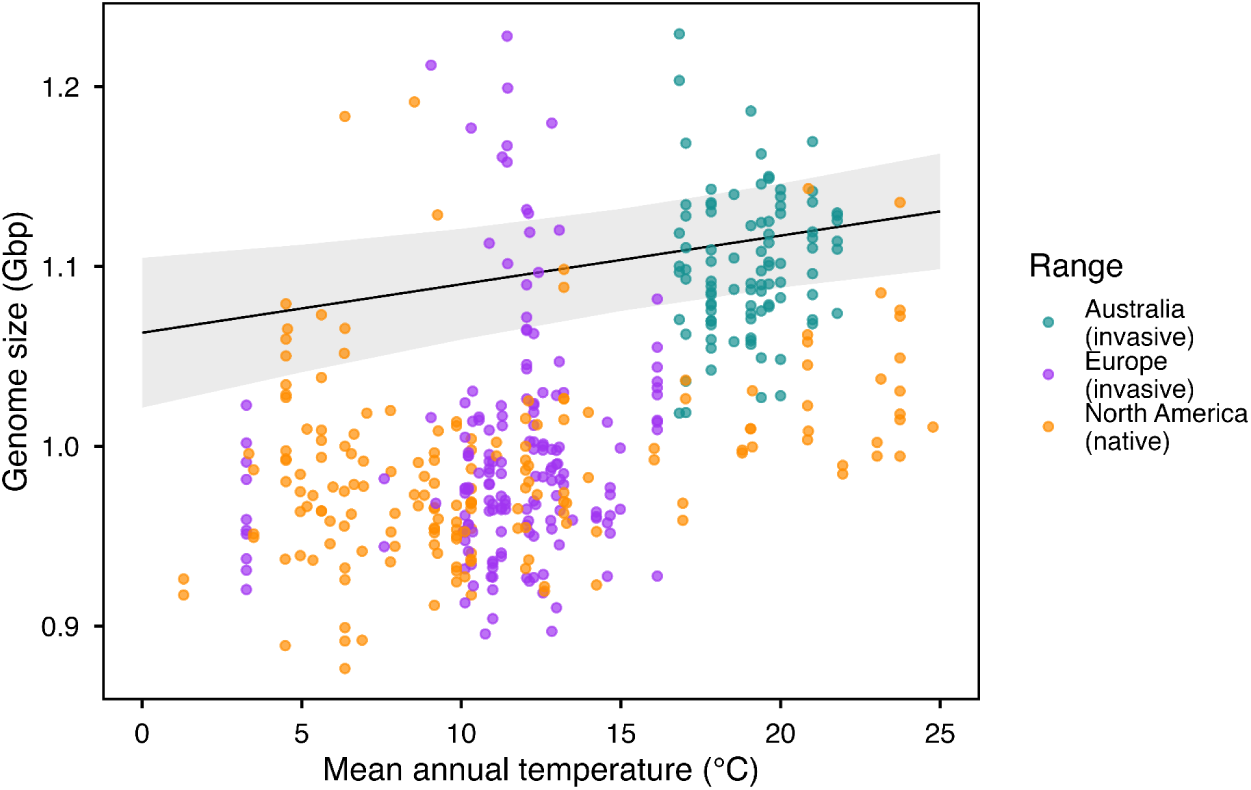
Relationship between genome size and MAT across the ranges. Scatter plot showing the estimated genome size (Gbp) of individual samples as a function of their corresponding MAT (°C) at the sampling site. The black line represents the estimated linear relationship from a mixed-effects model (model R² = 0.61, P = 0.0047), which included MAT, genetic principal components (PC1 and PC2), and range as fixed effects, and population as a random effect. The grey shaded area indicates the 95% confidence interval around the regression line.

Similar to North American populations, European common ragweed exhibits elevated *Q_ST_* and *P_ST_* relative to *F_ST_*, supporting range-wide divergent selection on genome size. While *Q_PC_*analysis provided weaker support for selection in Europe than in North America (Suppl. Material 4b), likely reflecting lower power or the distribution of adaptive divergence across multiple axes of genetic structure, both *Q_ST_–F_ST_*and *P_ST_–F_ST_* analyses consistently indicated that genome size differentiation exceeds neutral expectations. To our knowledge, neither *Q_ST_* nor *P_ST_* has been estimated for genome size in plants, so it remains unclear how often genome-wide structural traits diverge from neutral expectations. *Q_ST_–F_ST_* comparisons for gene/genomic copy-number variation in this species similarly revealed signatures of divergent selection on genomic variation in both Europe and North America (Wilson et al, 2025). Together, these results suggest that genome size differentiation has repeatedly evolved beyond neutral expectations across continents, although the relative importance of the underlying evolutionary drivers may vary among regions.

The Australian invasion some 100 years ago relied on two distinct introductions: a main genetic cluster that has experienced a bottleneck (Putra et al, 2024; Battlay et al, 2025a), originating from high-MAT sources in the southern and mideastern North American range, and a rarer genetic cluster with ancestry from eastern North America (Battlay et al, 2025a). Despite these different origins, both Australian clusters consistently exhibit larger genome sizes than their respective native source populations (Suppl. Fig. S10). Within-Australia *P_ST_*fell below neutral expectations, while *Q_ST_* was only marginally lower than expected under neutrality (Fig. 5; Suppl. Material SM4b), consistent with a relatively homogeneous genome size increase across populations. The absence of a significant *Q_PC_* signal in Australia (Suppl. Material SM4c) is consistent with this pattern, suggesting no divergent selection within the range. Introduction bottlenecks likely reduced effective population size, facilitating an increase in TE copy number through relaxed purifying selection, as observed in other invasive species (Merel et al, 2021), with TE remobilisation in novel environments potentially contributing further (Schrader et al, 2014). Although reduced efficacy of purifying selection following parallel bottlenecks could facilitate directional increases in TE load and genome size, such a mechanism alone would be expected to generate more heterogeneous outcomes among independently founded populations. The repeated increase in genome size across two independent introductions is therefore difficult to reconcile by drift alone, and is more consistent with colonisation filtering of large-genome genotypes and/or parallel responses to shared environmental conditions following introduction (Cang et al, 2024; Suda et al, 2015; Stapley et al, 2015; Prentis et al, 2008). Pangenomic analyses of TE insertion age spectra, combined with expression profiling and reciprocal-transplant assays, will help discriminate between demography-driven relaxation of constraint and environment-mediated selection.

## Conclusion

Genome size variation in common ragweed likely arises from the combined effects of demographic history, repetitive-element dynamics, and environmental context. Across North America and Europe, genome size differentiation is consistent with spatially varying selection, with climate-associated differentiation evident across all ranges. In contrast, the uniformly larger genomes of Australian plants are potentially explained by demographic processes associated with invasion, likely shaped or reinforced by shared environmental conditions. Together, these findings reframe genome size from a static genomic property into a dynamic, evolutionarily labile trait that responds to demographic and ecological pressures accompanying range expansion. A key frontier now lies in resolving the mechanistic links between specific transposable element families, climate adaptation, and fitness; particularly whether TE-driven genome expansion represents a recurrent evolutionary response to novel environments, or a largely neutral outcome of founder dynamics that persists until purifying selection erodes it. Addressing these questions across multiple invasive taxa will be essential for understanding how genome architecture *–* shapes or is shaped by *–* the colonisation of new ranges.

## Acknowledgements

We thank Naomi Drego for assistance with preliminary aspects of this project. We thank A/Prof. Tim Connallon for suggesting the *P_ST_–F_ST_* analysis. We also acknowledge the FlowCore facility at Monash University for access to their flow cytometry platforms, and A/Prof. Michal Hrabovský for assistance with the flow cytometry protocol for this species. This work was supported by the ARC Discovery Grant (DP220102362), NSERC Discovery Grants (JRS), and Graduate Fellowships (KGM).

## Authors contributions

KAH, PB and BN conceptualized the study. BN and PB developed the methodology and software. Resources were provided by KAH, MDM, AFL and KGM, and data curation was performed by PB. BN conducted the formal analyses, with support from KAH, PB and JRS, and created the visualizations. BN drafted the manuscript, with review and editing by all authors. Supervision and project administration were provided by KAH, PB and JRS. Funding was acquired by KAH, MDM, JRS and AFL. All authors contributed and approved the final version of the manuscript.

## Data availability

No new sequencing data were generated in this study. The whole-genome resequencing data used were obtained from Bieker et al. (2022; https://doi.org/10.1126/sciadv.abo5115) and Battlay et al. (2025; https://doi.org/10.1093/molbev/msae270). Flow cytometry genome size estimates generated in this study are provided in Suppl. Table S2. All code used in this study is available at the following github repository: https://github.com/Byonkesh/Ragweed_Genomesize_Evolution.

## References

1. Adams, P. E. et al. Genome Size Changes by Duplication, Divergence, and Insertion in Caenorhabditis Worms. Mol Biol Evol 40 (2023). 10.1093/molbev/msad039

2. Ajay, S. S., Parker, S. C., Abaan, H. O., Fajardo, K. V. & Margulies, E. H. Accurate and comprehensive sequencing of personal genomes. Genome Res 21, 1498–1505 (2011). 10.1101/gr.123638.111

3. Amadeu, R. R., Garcia, A. A. F., Munoz, P. R. & Ferrão, L. F. V. AGHmatrix: genetic relationship matrices in R. Bioinformatics 39 (2023). 10.1093/bioinformatics/btad445

4. Arkhipova, I. R. Neutral Theory, Transposable Elements, and Eukaryotic Genome Evolution. Mol Biol Evol 35, 1332–1337 (2018). 10.1093/molbev/msy083

5. Awoleye, F., Van Duren, M., Dolezel, J. & Novak, F. Nuclear DNA content and in vitro induced somatic polyploidization cassava (Manihot esculenta Crantz) breeding. Euphytica 76, 195–202 (1994). 10.1007/BF00022164

6. Badouin, H. et al. The sunflower genome provides insights into oil metabolism, flowering and Asterid evolution. Nature 546, 148–152 (2017). 10.1038/nature22380

7. Bai, C., Alverson, W. S., Follansbee, A. & Waller, D. M. New reports of nuclear DNA content for 407 vascular plant taxa from the United States. Annals of botany 110, 1623–1629 (2012). 10.1093/aob/mcs222

8. Bates, D., Mächler, M., Bolker, B. & Walker, S. Fitting Linear Mixed-Effects Models Using lme4. Journal of Statistical Software 67, 1–48 (2015). 10.18637/jss.v067.i01

9. Battlay, P. et al. Rapid Parallel Adaptation in Distinct Invasions of Ambrosia Artemisiifolia Is Driven by Large-Effect Structural Variants. Molecular Biology and Evolution 42 (2025a). 10.1093/molbev/msae270

10. Battlay, P., Bieker, V. C., Wu, T., Martin, M. D. & Hodgins, K. A. How DNA Sequencing of Herbarium Specimens Can Elucidate Our Understanding of Plant Invasions: Insights from Common Ragweed. CABI Books, 103–117 (2025b). 10.1079/9781800626263.0008

11. Battlay, P. et al. Large haploblocks underlie rapid adaptation in the invasive weed Ambrosia artemisiifolia. Nat Commun 14, 1717 (2023). 10.1038/s41467-023-37303-4

12. Bennett, M. D. Nuclear DNA content and minimum generation time in herbaceous plants. Proceedings of the Royal Society of London. Series B. Biological Sciences 181, 109–135 (1972). 10.1098/rspb.1972.0042

13. Bennett, M. D. Variation in genomic form in plants and its ecological implications. New Phytologist 106, 177–200 (1987). /10.1111/j.1469-8137.1987.tb04689.x

14. Bhadra, S., Leitch, I. J. & Onstein, R. E. From genome size to trait evolution during angiosperm radiation. Trends in Genetics 39, 728–735 (2023). 10.1016/j.tig.2023.07.006

15. Bieker, V. C. et al. Uncovering the genomic basis of an extraordinary plant invasion. Science Advances 8, eabo5115 (2022). doi:10.1126/sciadv.abo5115

16. Bilinski, P. et al. Parallel altitudinal clines reveal trends in adaptive evolution of genome size in Zea mays. PLoS genetics 14, e1007162 (2018). 10.1371/journal.pgen.1007162

17. Blommaert, J. Genome size evolution: towards new model systems for old questions. Proceedings of the Royal Society B: Biological Sciences 287, 20201441 (2020). doi:10.1098/rspb.2020.1441

18. Bredeson, J. V. et al. Sequencing wild and cultivated cassava and related species reveals extensive interspecific hybridization and genetic diversity. Nature biotechnology 34, 562–570 (2016). 10.1038/nbt.3535

19. Brommer, J. E. Whither Pst? The approximation of Qst by Pst in evolutionary and conservation biology. Journal of Evolutionary Biology 24, 1160–1168 (2011). 10.1111/j.1420-9101.2011.02268.x

20. Bureš, P. et al. The global distribution of angiosperm genome size is shaped by climate. New Phytologist 242, 744–759 (2024). 10.1111/nph.19544

21. Cang, F. A., Welles, S. R., Wong, J., Ziaee, M. & Dlugosch, K. M. Genome size variation and evolution during invasive range expansion in an introduced plant. Evolutionary Applications 17, e13624 (2024). 10.1111/eva.13624

22. Cavaller-Smith, T. The evolution of genome size. (1985).

23. Cerca, J. et al. The genomic basis of the plant island syndrome in Darwin’s giant daisies. Nature Communications 13, 3729 (2022). 10.1038/s41467-022-31280-w

24. Chen, J. et al. A complete telomere-to-telomere assembly of the maize genome. Nature Genetics 55, 1221–1231 (2023). 10.1038/s41588-023-01419-6

25. Chen, S., Zhou, Y., Chen, Y. & Gu, J. fastp: an ultra-fast all-in-one FASTQ preprocessor. Bioinformatics 34, i884–i890 (2018). 10.1093/bioinformatics/bty560

26. Danecek, P. et al. Twelve years of SAMtools and BCFtools. Gigascience 10 (2021). 10.1093/gigascience/giab008

27. Dazenière, J., Bousios, A. & Eyre-Walker, A. Patterns of selection in the evolution of a transposable element. G3 (Bethesda) 12 (2022). 10.1093/g3journal/jkac056

28. de Villemereuil, P. Quantitative genetic methods depending on the nature of the phenotypic trait. Annals of the New York Academy of Sciences 1422, 29–47 (2018). 10.1111/nyas.13571

29. Dennenmoser, S. et al. Copy number increases of transposable elements and protein-coding genes in an invasive fish of hybrid origin. Mol Ecol 26, 4712–4724 (2017). 10.1111/mec.14134

30. Desvillechabrol, D., Bouchier, C., Kennedy, S. & Cokelaer, T. Detection and characterization of low and high genome coverage regions using an efficient running median and a double threshold approach. bioRxiv, 092478 (2016). 10.1101/092478

31. Díez, C. M. et al. Genome size variation in wild and cultivated maize along altitudinal gradients. New Phytologist 199, 264–276 (2013). 10.1111/nph.12247

32. Dirkse, G. M., H., D. & and Zonneveld, B. J. M. Morphology and genome weight of Symphyotrichum species (Asteraceae) along rivers in The Netherlands. New Journal of Botany 4, 134–142 (2014). 10.1179/2042349714Y.0000000049

33. Doležel, J., Sgorbati, S. & Lucretti, S. Comparison of three DNA fluorochromes for flow cytometric estimation of nuclear DNA content in plants. Physiologia plantarum 85, 625–631 (1992). 10.1111/j.1399-3054.1992.tb04764.x

34. Elliott, T. A. & Gregory, T. R. What’s in a genome? The C-value enigma and the evolution of eukaryotic genome content. Philosophical Transactions of the Royal Society B: Biological Sciences 370, 20140331 (2015). doi:10.1098/rstb.2014.0331

35. Falconer, D. S. Introduction to quantitative genetics. (Pearson Education India, 1996). 10.1093/genetics/167.4.1529

36. Fedoroff, N. V. Transposable Elements, Epigenetics, and Genome Evolution. Science 338, 758–767 (2012). doi:10.1126/science.338.6108.758

37. Fernández, P. et al. A 160 Gbp fork fern genome shatters size record for eukaryotes. iScience 27, 109889 (2024). 10.1016/j.isci.2024.109889

38. Fick, S. E. & Hijmans, R. J. WorldClim 2: new 1-km spatial resolution climate surfaces for global land areas. International Journal of Climatology 37, 4302–4315 (2017). 10.1002/joc.5086

39. Fox, J. & Weisberg, S. An R companion to applied regression. (Sage publications, 2018). https://www.john-fox.ca/Companion/

40. Gallagher, R. V. et al. Invasiveness in introduced Australian acacias: the role of species traits and genome size. Diversity and Distributions 17, 884–897 (2011). 10.1111/j.1472-4642.2011.00805.x

41. Glémin, S. Mating systems and the efficacy of selection at the molecular level. Genetics 177, 905–916 (2007). 10.1534/genetics.107.073601

42. Goubert, C. et al. High-throughput sequencing of transposable element insertions suggests adaptive evolution of the invasive Asian tiger mosquito towards temperate environments. Mol Ecol 26, 3968–3981 (2017). 10.1111/mec.14184

43. Gregory, T. R. Insertion–deletion biases and the evolution of genome size. Gene 324, 15–34 (2004). 10.1016/j.gene.2003.09.030

44. Gregory, T. R. The C-value Enigma in Plants and Animals: A Review of Parallels and an Appeal for Partnership. Annals of Botany 95, 133–146 (2005). 10.1093/aob/mci009

45. Greilhuber, J. Intraspecific Variation in Genome Size in Angiosperms: Identifying its Existence. Annals of Botany 95, 91–98 (2005). 10.1093/aob/mci004

46. Grime, J. & Mowforth, M. Variation in genome size—an ecological interpretation. Nature 299, 151–153 (1982). 10.1038/299151a0

47. Grime, J. P. Plant Classification for Ecological Purposes: is there a Role for Genome Size? Annals of Botany 82, 117–120 (1998). 10.1006/anbo.1998.0723

48. Gu, Z., Gu, L., Eils, R., Schlesner, M. & Brors, B. “Circlize” implements and enhances circular visualization in R. (2014). 10.1093/bioinformatics/btu393

49. Guo, K. et al. Plant invasion and naturalization are influenced by genome size, ecology and economic use globally. Nature Communications 15, 1330 (2024). 10.1038/s41467-024-45667-4

50. Hadfield, J. D. MCMC Methods for Multi-Response Generalized Linear Mixed Models: The MCMCglmm R Package. Journal of Statistical Software 33, 1–22 (2010). 10.18637/jss.v033.i02

51. Hirsch, C. D. & Springer, N. M. Transposable element influences on gene expression in plants. Biochim Biophys Acta Gene Regul Mech 1860, 157–165 (2017). 10.1016/j.bbagrm.2016.05.010

52. Hodgins, K. A., Battlay, P. & Bock, D. G. The genomic secrets of invasive plants. New Phytologist 245, 1846–1863 (2025). 10.1111/nph.20368

53. Hrabovský, M., Kubalová, S. & Kanka, R. The impact of changing climate on the spread of the widely expanding species Ambrosia artemisiifolia in Slovakia. Theoretical and Applied Climatology 155, 6137–6150 (2024). 10.1007/s00704-024-05006-5

54. Hrabovský, M., Kubalová, S., Mičieta, K. & Ščevková, J. Environmental impacts on intraspecific variation in Ambrosia artemisiifolia genome size in Slovakia, Central Europe. Environmental Science and Pollution Research 31, 33960–33974 (2024). 10.1007/s11356-024-33410-x

55. Huang, Y. et al. Polymorphic transposable elements contribute to variation in recombination landscapes. Proceedings of the National Academy of Sciences 122, e2427312122 (2025). doi:10.1073/pnas.2427312122

56. Institute, B. Picard toolkit. Broad Institute, GitHub repository (2019). https://broadinstitute.github.io/picard/

57. Jiang, J. et al. Forces driving transposable element load variation during Arabidopsis range expansion. The Plant Cell 36, 840–862 (2024). 10.1093/plcell/koad296

58. Josephs, E. B., Berg, J. J., Ross-Ibarra, J. & Coop, G. Detecting adaptive differentiation in structured populations with genomic data and common gardens. Genetics 211, 989–1004 (2019). 10.1534/genetics.118.301786

59. Kent, T. V., Uzunović, J. & Wright, S. I. Coevolution between transposable elements and recombination. Philos Trans R Soc Lond B Biol Sci 372 (2017). 10.1098/rstb.2016.0458

60. Knight, C. A., Molinari, N. A. & Petrov, D. A. The large genome constraint hypothesis: evolution, ecology and phenotype. Annals of botany 95, 177–190 (2005). 10.1093/aob/mci011

61. Knolmajer, B., Jócsák, I., Taller, J., Keszthelyi, S. & Kazinczi, G. Common Ragweed—Ambrosia artemisiifolia L.: A Review with Special Regards to the Latest Results in Biology and Ecology. Agronomy 14, 497 (2024). 10.3390/agronomy14030497

62. Korneliussen, T. S., Albrechtsen, A. & Nielsen, R. ANGSD: Analysis of Next Generation Sequencing Data. BMC Bioinformatics 15, 356 (2014). 10.1186/s12859-014-0356-4

63. Kreiner, J. M., Hnatovska, S., Stinchcombe, J. R. & Wright, S. I. Quantifying the role of genome size and repeat content in adaptive variation and the architecture of flowering time in Amaranthus tuberculatus. PLoS genetics 19, e1010865 (2023). 10.1371/journal.pgen.1010865

64. Kubešová, M., Moravcova, L., Suda, J., Jarošík, V. & Pyšek, P. Naturalized plants have smaller genomes than their non-invading relatives: a flow cytometric analysis of the Czech alien flora. Preslia 82, 81–96 (2010). https://www.preslia.cz/article/225

65. Lagesen, K. et al. RNAmmer: consistent and rapid annotation of ribosomal RNA genes. Nucleic Acids Res 35, 3100–3108 (2007). 10.1093/nar/gkm160

66. Lande, R. Neutral theory of quantitative genetic variance in an island model with local extinction and colonization. Evolution 46, 381–389 (1992). 10.1111/j.1558-5646.1992.tb02046.x

67. Lander, E. S. & Waterman, M. S. Genomic mapping by fingerprinting random clones: A mathematical analysis. Genomics 2, 231–239 (1988). 10.1016/0888-7543(88)90007-9

68. Lefébure, T. et al. Less effective selection leads to larger genomes. Genome Research 27, 1016–1028 (2017). 10.1101/gr.212589.116

69. Leinonen, T., McCairns, R. S., O’hara, R. B. & Merilä, J. Q ST–F ST comparisons: evolutionary and ecological insights from genomic heterogeneity. Nature Reviews Genetics 14, 179–190 (2013). 10.1038/nrg3395

70. Lenth, R. emmeans: Estimated Marginal Means, aka Least-Squares Means_. R package version 1.8.5 (2023). https://cran.r-project.org/package=emmeans

71. Li, H. et al. The Sequence Alignment/Map format and SAMtools. Bioinformatics 25, 2078–2079 (2009). 10.1093/bioinformatics/btp352

72. Li, H. et al. Warming-induced miniaturization of plant community genome size in temperate grasslands over the last four decades. New Phytologist (2026). 10.1111/nph.70948

73. Li H. Aligning sequence reads, clone sequences and assembly contigs with BWA-MEM (2013). 10.48550/arXiv.1303.3997

74. Li, H., Ruan, J. & Durbin, R. Mapping short DNA sequencing reads and calling variants using mapping quality scores. Genome Res 18, 1851–1858 (2008). 10.1101/gr.078212.108

75. Lian, Q. et al. A pan-genome of 69 Arabidopsis thaliana accessions reveals a conserved genome structure throughout the global species range. Nature Genetics 56, 982–991 (2024). 10.1038/s41588-024-01715-9

76. Lisch, D. How important are transposons for plant evolution? Nat Rev Genet 14, 49–61 (2013). 10.1038/nrg3374

77. Lockton, S., Ross-Ibarra, J. & Gaut, B. S. Demography and weak selection drive patterns of transposable element diversity in natural populations of Arabidopsis lyrata. Proceedings of the National Academy of Sciences 105, 13965–13970 (2008). doi:10.1073/pnas.0804671105

78. Long, Q. et al. Massive genomic variation and strong selection in Arabidopsis thaliana lines from Sweden. Nature genetics 45, 884–890 (2013). 10.1038/ng.2678

79. Lopes, S. et al. Genome size variation in Cactaceae and its relationship with invasiveness and seed traits. Biological Invasions 23, 3047–3062 (2021). 10.1007/s10530-021-02557-w

80. Lynch, M. & Walsh, B. Genetics and analysis of quantitative traits. Vol. 1 (Sinauer Sunderland, MA, 1998). https://www.invemar.org.co/redcostera1/invemar/docs/RinconLiterario/2011/febrero/AG_8.pdf

81. Makarevitch, I. et al. Transposable elements contribute to activation of maize genes in response to abiotic stress. PLoS Genet 11, e1004915 (2015). 10.1371/journal.pgen.1004915

82. Marino, A., Debaecker, G., Fiston-Lavier, A.-S., Haudry, A. & Nabholz, B. (eLife Sciences Publications, Ltd, 2024). 10.7554/eLife.100574.1

83. Marshall, C. R. et al. Best practices for the analytical validation of clinical whole-genome sequencing intended for the diagnosis of germline disease. NPJ Genom Med 5, 47 (2020). 10.1038/s41525-020-00154-9

84. McGoey, B. V., Hodgins, K. A. & Stinchcombe, J. R. Parallel flowering time clines in native and introduced ragweed populations are likely due to adaptation. Ecology and Evolution 10, 4595–4608 (2020). 10.1002/ece3.6163

85. McKenna, A. et al. The Genome Analysis Toolkit: a MapReduce framework for analyzing next-generation DNA sequencing data. Genome Res 20, 1297–1303 (2010). 10.1101/gr.107524.110

86. Meisner, J. & Albrechtsen, A. Inferring population structure and admixture proportions in low-depth NGS data. Genetics 210, 719–731 (2018). 10.1534/genetics.118.301336

87. Mérel, V. et al. The Worldwide Invasion of Drosophila suzukii Is Accompanied by a Large Increase of Transposable Element Load and a Small Number of Putatively Adaptive Insertions. Mol Biol Evol 38, 4252–4267 (2021). 10.1093/molbev/msab155

88. Meyerson, L. A. et al. Some like it hot: small genomes may be more prevalent under climate extremes. Biological Invasions 26, 1425–1436 (2024). 10.1007/s10530-024-03253-1

89. Natarajan, S., Gehrke, J. & Pucker, B. Mapping-based genome size estimation. BMC Genomics 26, 482 (2025). 10.1186/s12864-025-11640-8

90. Nilforooshan, M. A. mbend: an R package for bending non-positive-definite symmetric matrices to positive-definite. BMC genetics 21, 97 (2020). 10.1186/s12863-020-00881-z

91. Niu, X. M. et al. Transposable elements drive rapid phenotypic variation in Capsella rubella. Proc Natl Acad Sci U S A 116, 6908–6913 (2019). 10.1073/pnas.1811498116

92. Ou, S. et al. Benchmarking transposable element annotation methods for creation of a streamlined, comprehensive pipeline. Genome Biology 20, 275 (2019). 10.1186/s13059-019-1905-y

93. Parks, M. M. et al. Variant ribosomal RNA alleles are conserved and exhibit tissue-specific expression. Science Advances 4, eaao0665 (2018). doi:10.1126/sciadv.aao0665

94. Pellicer, J., Fay, M. F. & Leitch, I. J. The largest eukaryotic genome of them all? Botanical Journal of the Linnean Society 164, 10–15 (2010). 10.1111/j.1095-8339.2010.01072.x

95. Perrier, C., Delahaie, B. & Charmantier, A. Heritability estimates from genomewide relatedness matrices in wild populations: Application to a passerine, using a small sample size. Molecular ecology resources 18, 838–853 (2018). 10.1111/1755-0998.12886

96. Pfenninger, M., Schönnenbeck, P. & Schell, T. ModEst: Accurate estimation of genome size from next generation sequencing data. Molecular Ecology Resources 22, 1454–1464 (2022). 10.1111/1755-0998.13570

97. Pflug, J. M., Holmes, V. R., Burrus, C., Johnston, J. S. & Maddison, D. R. Measuring Genome Sizes Using Read-Depth, k-mers, and Flow Cytometry: Methodological Comparisons in Beetles (Coleoptera). G3 (Bethesda) 10, 3047–3060 (2020). 10.1534/g3.120.401028

98. Poptsova, M. S. et al. Non-random DNA fragmentation in next-generation sequencing. Scientific reports 4, 4532 (2014). 10.1038/srep04532

99. Potapenko, E., Shermeister, B., Mandel, T. & Hübner, S. Genome size variation is attributed to adaptive purging of transposable elements. bioRxiv, 2025.2009.2026.678839 (2025). 10.1101/2025.09.26.678839

100. Prentis, P. J., Wilson, J. R., Dormontt, E. E., Richardson, D. M. & Lowe, A. J. Adaptive evolution in invasive species. Trends Plant Sci 13, 288–294 (2008). 10.1016/j.tplants.2008.03.004

101. Prout, T. & Barker, J. S. F statistics in Drosophila buzzatii: selection, population size and inbreeding. Genetics 134, 369–375 (1993). 10.1093/genetics/134.1.369

102. Purcell, S. et al. PLINK: a tool set for whole-genome association and population-based linkage analyses. Am J Hum Genet 81, 559–575 (2007). 10.1086/519795

103. Putra, A. R., Hodgins, K. A. & Fournier-Level, A. Assessing the invasive potential of different source populations of ragweed (Ambrosia artemisiifolia L.) through genomically informed species distribution modelling. Evolutionary applications 17, e13632 (2024). 10.1111/eva.13632

104. Pyšek, P. et al. Small genome size and variation in ploidy levels support the naturalization of vascular plants but constrain their invasive spread. New Phytologist 239, 2389–2403 (2023). 10.1111/nph.19135

105. Pyšek, P. et al. Small genome separates native and invasive populations in an ecologically important cosmopolitan grass. Ecology 99, 79–90 (2018). 10.1002/ecy.2068

106. Quinlan, A. R. & Hall, I. M. BEDTools: a flexible suite of utilities for comparing genomic features. Bioinformatics 26, 841–842 (2010). 10.1093/bioinformatics/btq033

107. Rabiner, L. & Juang, B. An introduction to hidden Markov models. IEEE ASSP Magazine 3, 4–16 (1986). 10.1109/MASSP.1986.1165342

108. Richardson, D. M. & Pyšek, P. Plant invasions: merging the concepts of species invasiveness and community invasibility. Progress in physical geography 30, 409–431 (2006). 10.1191/0309133306pp490pr

109. Robinson, D. O. et al. Ploidy and Size at Multiple Scales in the Arabidopsis Sepal. Plant Cell 30, 2308–2329 (2018). 10.1105/tpc.18.00344

110. Roddy, A. B. et al. The scaling of genome size and cell size limits maximum rates of photosynthesis with implications for ecological strategies. International Journal of Plant Sciences 181, 75–87 (2020). 10.1086/706186

111. Sæther, S. A. et al. Inferring local adaptation from QST–FST comparisons: neutral genetic and quantitative trait variation in European populations of great snipe. Journal of Evolutionary Biology 20, 1563–1576 (2007). 10.1111/j.1420-9101.2007.01328.x

112. Schrader, L. et al. Transposable element islands facilitate adaptation to novel environments in an invasive species. Nature Communications 5, 5495 (2014). 10.1038/ncomms6495

113. Schrader, L. & Schmitz, J. The impact of transposable elements in adaptive evolution. Mol Ecol 28, 1537–1549 (2019). 10.1111/mec.14794

114. Simão, F. A., Waterhouse, R. M., Ioannidis, P., Kriventseva, E. V. & Zdobnov, E. M. BUSCO: assessing genome assembly and annotation completeness with single-copy orthologs. Bioinformatics 31, 3210–3212 (2015). 10.1093/bioinformatics/btv351

115. Šímová, I. & Herben, T. Geometrical constraints in the scaling relationships between genome size, cell size and cell cycle length in herbaceous plants. Proceedings of the Royal Society B: Biological Sciences 279, 867–875 (2012). 10.1098/rspb.2011.1284

116. Sims, J., Sestini, G., Elgert, C., von Haeseler, A. & Schlögelhofer, P. Sequencing of the Arabidopsis NOR2 reveals its distinct organization and tissue-specific rRNA ribosomal variants. Nature communications 12, 387 (2021). 10.1038/s41467-020-20728-6

117. Slijepcevic, P. Genome dynamics over evolutionary time: “C-value enigma” in light of chromosome structure. Mutation Research/Genetic Toxicology and Environmental Mutagenesis 836, 22–27 (2018). 10.1016/j.mrgentox.2018.05.005m

118. Sliwinska, E. et al. Application-based guidelines for best practices in plant flow cytometry. Cytometry Part A 101, 749–781 (2022). 10.1002/cyto.a.24499

119. Slovák, M., Vít, P., Urfus, T. & Suda, J. Complex pattern of genome size variation in a polymorphic member of the Asteraceae. Journal of Biogeography 36, 372–384 (2009). 10.1111/j.1365-2699.2008.02005.x

120. Šmarda, P. Understanding intraspecific variation in genome size in plants. Preslia 82, 41–61 (2010). https://www.preslia.cz/article/223

121. Šmarda, P., Horová, L., Bureš, P., Hralová, I. & Marková, M. Stabilizing selection on genome size in a population of Festuca pallens under conditions of intensive intraspecific competition. New Phytologist 187, 1195–1204 (2010). 10.1111/j.1469-8137.2010.03335.x

122. Smit, A., Hubley, R & Green,,P. Repeatmasker Open-4.0. (2013-2015). https://www.repeatmasker.org/

123. Smith, D. R. The mutational hazard hypothesis of organelle genome evolution: 10 years on. Molecular Ecology 25, 3769–3775 (2016). 10.1111/mec.13742

124. Spitze, K. Population structure in Daphnia obtusa: quantitative genetic and allozymic variation. Genetics 135, 367–374 (1993). 10.1093/genetics/135.2.367

125. Stapley, J., Santure, A. W. & Dennis, S. R. Transposable elements as agents of rapid adaptation may explain the genetic paradox of invasive species. Mol Ecol 24, 2241–2252 (2015). 10.1111/mec.13089

126. Stoffel, M. A., Nakagawa, S. & Schielzeth, H. partR2: partitioning R(2) in generalized linear mixed models. PeerJ 9, e11414 (2021). 10.7717/peerj.11414

127. Suda, J. et al. Genome Size Variation and Species Relationships in Hieracium Sub-genus Pilosella (Asteraceae) as Inferred by Flow Cytometry. Annals of Botany 100, 1323–1335 (2007). 10.1093/aob/mcm218

128. Suda, J., Meyerson, L. A., Leitch, I. J. & Pyšek, P. The hidden side of plant invasions: the role of genome size. New Phytologist 205, 994–1007 (2015). 10.1111/nph.13107

129. Sultanov, D. & Hochwagen, A. Varying strength of selection contributes to the intragenomic diversity of rRNA genes. Nature Communications 13, 7245 (2022). 10.1038/s41467-022-34989-w

130. Sun, Y., Bossdorf, O., Grados, R. D., Liao, Z. & Müller-Schärer, H. Rapid genomic and phenotypic change in response to climate warming in a widespread plant invader. Global Change Biology 26, 6511–6522 (2020). 10.1111/gcb.15291

131. Tenaillon, M. I., Manicacci, D., Nicolas, S. D., Tardieu, F. & Welcker, C. Testing the link between genome size and growth rate in maize. PeerJ 4, e2408 (2016). 10.7717/peerj.2408

132. van Boheemen, L. A., Atwater, D. Z. & Hodgins, K. A. Rapid and repeated local adaptation to climate in an invasive plant. New Phytologist 222, 614–627 (2019). 10.1111/nph.15564

133. van Heesch, S. et al. Improving mammalian genome scaffolding using large insert mate-pair next-generation sequencing. BMC genomics 14, 257 (2013). 10.1186/1471-2164-14-257

134. VanRaden, P. M. Efficient Methods to Compute Genomic Predictions. Journal of Dairy Science 91, 4414–4423 (2008). 10.3168/jds.2007-0980

135. Weir, B. & Cockerham, C., Weir BS, Cockerham CC.. Estimating F-Statistics for the Analysis of Population-Structure. Evolution 38: 1358-1370. Evolution 38, 1358–1370 (1984). 10.2307/2408641

136. Whitlock, M. C. Evolutionary inference from QST. Molecular ecology 17, 1885–1896 (2008). 10.1111/j.1365-294x.2008.03712.x

137. Whitlock, M. C. & Guillaume, F. Testing for spatially divergent selection: comparing QST to FST. Genetics 183, 1055–1063 (2009). 10.1534/genetics.108.099812

138. Wicker, T. et al. A unified classification system for eukaryotic transposable elements. Nat Rev Genet 8, 973–982 (2007). 10.1038/nrg2165

139. Wickham, H. in Elegant Graphics for Data Analysis 2197-5744 XVI, 260 (Springer Cham, 2016). 10.1007/978-3-319-24277-4

140. Wilson, A. J. et al. An ecologist’s guide to the animal model. Journal of animal ecology 79, 13–26 (2010). 10.1111/j.1365-2656.2009.01639.x

141. Wilson, J. et al. Copy number variation contributes to parallel local adaptation in an invasive plant. Proceedings of the National Academy of Sciences 122, e2413587122 (2025). doi:10.1073/pnas.2413587122

142. Wu, N. et al. Fall webworm genomes yield insights into rapid adaptation of invasive species. Nature Ecology & Evolution 3, 105–115 (2019). 10.1038/s41559-018-0746-5

143. Xiao, Q.-S. et al. Small genome size ensures adaptive flexibility for an alpine ginger. Genome Biology and Evolution 17, evaf151 (2025). 10.1093/gbe/evaf151

144. Zedek, F. et al. The smallest angiosperm genomes may be the price for effective traps of bladderworts. Annals of Botany 134, 1131–1138 (2024). 10.1093/aob/mcae107

