## Supplementary Materials, Figures and Tables for "Selection shapes the evolution of genome size in a globally invasive plant"

**Affiliations:**
[1]School of Biological Sciences, Monash University, 25 Rainforest Walk, Clayton 3800, Victoria, Australia.
[2]Department of Ecology and Evolutionary Biology, University of Toronto, Toronto, ON M5S3B2, Canada.
[3]Department of Ecological, Plant and Animal Sciences, La Trobe University, Bundoora, VIC 3086, Australia.
[4]Department of Natural History, NTNU University Museum, Norwegian University of Science and Technology (NTNU), Trondheim, Norway.

***Correspondence:**
Byonkesh Nongthongbam
https://orcid.org/0009-0009-0967-147X

**A) Supplementary materials:**

**Suppl. Material (SM1):**

*Quality controls addressing biases in genome size estimation from sequencing data*

We systematically evaluated potential technical sources of error that could influence our results because of the known biases associated with sequencing-based genome size estimation (Poptsova et al, 2014; van Heesch et al, 2013). Given the relatively low average sequencing depth (~7.1×) across our samples, we first assessed whether coverage depth introduced systematic bias into our genome size and TE abundance estimates. We regressed whole-genome average depth against estimated genome size and against the abundance of each TE family. Genome size showed no meaningful association with coverage ($R^2 = 0.00028$; Suppl. Fig. S2), confirming that low-coverage samples did not systematically under- or overestimate genome size. Similarly, the vast majority of TE families showed negligible correlations with coverage ($R^2 < 0.05$; Suppl. Fig. S8), with the exception of rRNA repeat families (18S, 28S, 8S), which showed moderate associations ($R^2 = 0.10$-$0.23$; Suppl. Fig. S8). These rRNA patterns likely reflect genuine biological variation in rRNA copy number rather than technical bias, as rRNA genes are known to vary considerably in copy number across individuals (Parks et al, 2018; Sims et al, 2021; Sultanov et al, 2022). Together, these results indicate that sequencing depth did not systematically bias our genome size or TE abundance estimates

Having established that coverage depth was not a confounding factor, we next assessed a broader set of quality metrics related to read integrity, coverage uniformity, and sequence-dependent fragmentation, all of which are known to introduce bias in depth-based genome size inference. We first examined read insert size (representing the distance between paired-end reads), length and mapping quality as potential indicators of DNA fragmentation and alignment confidence, respectively (Li et al, 2008). Average insert size was calculated using the `IS` option of `samtools stats` in samtools v.1.16.1 (Li et al, 2009). Our analysis revealed no significant linear correlation with average insert size and genome size ($R^2$ = 0.011; Suppl. Fig. S3a). Average read length was calculated using the `RL` option of `samtools stats` and showed no clear linear correlation with genome size ($R^2$ = 0; Suppl. Fig. S3b), indicating that sequencing-induced DNA fragmentation did not bias our estimates. Furthermore, applying various mapping quality thresholds (MAPQ ≥ 2, 5, 7, 10, and 15) and recalculating genome size estimates using `samtools depth -Q` demonstrated consistent and strong relationship with genome size estimated without filters across all thresholds (Suppl. Fig. S4). Thus, alignment confidence did not significantly affect our genome size estimations.

We next assessed the uniformity of genome-wide coverage and the impact of PCR duplication. Coverage depth was evaluated at callable loci, defined as genomic positions with ≥10× read coverage and sufficient quality for reliable consensus sequence generation (Ajay et al, 2011; Marshall et al, 2020), as established by Bieker et al. (2022). By calculating depth at these loci (`samtools depth -b callable_depth.bed`) and comparing it to whole-genome sequencing depth, we observed a strong linear relationship ($R^2$ = 0.99; Suppl. Fig. S3c), suggesting even coverage across the genome, and validating the use of whole-genome depth for genome size estimation. Additionally, we examined BUSCO-based scaling, which was used to normalize genome-wide sequencing depth by the average depth across conserved, single-copy BUSCO genes. To minimize potential biases from structural variation or inconsistent coverage, we computed depth variance at each BUSCO gene site across all samples using `samtools depth`. Only single-copy BUSCO genes with depth variance within the first standard deviation were retained, effectively excluding regions prone to structural variation such as copy number variations (adapted from Gilbert, 2024). Genome size for each sample was then estimated based on these filtered BUSCO site depths. Comparing these BUSCO-filtered genome size estimates to unfiltered estimates revealed a significant relationship, confirming that our scaling approach did not introduce systematic bias ($R^2$ = 0.97; Suppl. Fig. S3d). Finally, we evaluated the effect of clonality (PCR duplicates) on our results. Since our standard pipeline removes duplicates, we used unfiltered aligned reads to calculate clonality (identified via `samtools flagstat`) and re-estimate genome size. We found only a weak correlation between clonality and genome size estimates including duplicates ($R^2$ = 0.069; Suppl. Fig. S3e) suggesting PCR artifacts did not bias the data. Additionally, the geographic patterns of genome size remained consistent when using estimates that included duplicates (Australia > Europe > North America; Suppl. Fig. S3f), also indicating the lack of clonality effect on our estimates. Together, these quality assessments demonstrate that our genome size estimates are robust against known technical and biological sources of sequencing bias.

These controls provide confidence that observed geographic patterns in genome size do not reflect methodological artifacts.

Although short-read-based genome size estimates can be sensitive to coverage and reference choice, multiple independent validation approaches; including flow cytometry confirming that Australian genomes are larger than those from North America and Europe, and sensitivity analyses spanning insert size, read length, mapping quality, callable base depth, clonality, and BUSCO gene depth variance, strongly indicate that the observed range differences and genome size–climate associations reflect genuine biological signal rather than methodological artefacts.

**Suppl. Material (SM2):** *Genome size estimation by flow-cytometry*

To independently validate genome size estimates obtained from the resequencing approach, we performed flow cytometry. Although different individuals were used due to limited tissue availability, all samples originated from the same populations as the sequenced individuals. Sample sizes for flow cytometry per range were determined using power analyses based on effect sizes estimated from sequencing-based genome size differences among ranges, ensuring sufficient power to detect the observed genome size pattern.

Genome size estimation followed established protocols (Sliwinska et al, 2022; Hrabovský et al, 2024). Plants were grown in a common garden at Koffler Science Reserve in Ontario, Canada. Leaf samples were collected in late September 2024, dried with silica gel powder and stored at room temperature until analysis. In total, 40 samples were used (exceeding the minimum sample size per ranges required based on power analyses): 16 from North America, 6 from Europe, and 18 from Australia (Suppl. Table S2). Nuclear suspensions were prepared by co-chopping 60–70 mg of dried common ragweed leaf tissue with 60–70 mg of *Manihot esculenta* (cassava-red stem variety) leaves as an internal control. Cassava was used as internal reference standard for common ragweed because its genome size (~600–772 Mbp/C based on assembly estimates; Bredeson et al, 2016; Awoleye et al, 1994) is smaller yet sufficiently close to the common ragweed genome (~1.02–1.11 Gbp/C; Kubešová et al, 2010; Bai et al, 2012; Battlay et al, 2023) to ensure non-overlapping but adjacent fluorescence peaks during flow cytometry, making it a favourable control. We first estimated the genome size of the cassava individual used, by calibrating it against a standard control, *Solanum lycopersicum* L. (variety: 'Stupické polní rané' (2C = 1.96 pg; Doležel et al. 1992). The calibrated cassava value (1.45 ± 0.01 Gbp/2C) was then used to normalise common ragweed genome size calculation. The homogenate was filtered through 2–3 layers of Miracloth to remove debris. Then, 5 µL of 1 mg/mL RNase was added into the suspensions and incubated at 37℃ for 30 minutes. DNA was then stained with 50 µL of propidium iodide (PI) (1 mg/mL), and fluorescence signals (standard 10,000 nuclei populations) were measured within an hour. For each sample, measurements were also taken from unstained aliquots to assess and correct for background fluorescence. Flow cytometry was performed on a FACSymphony A3 analyzer (BD Biosciences, USA) at FlowCore, Monash University and the data were analysed using FlowJo software (FlowJo LLC, USA). The genome size (2C) was calculated based on the mean fluorescence intensity of PI-stained nuclei, normalized to the cassava reference genome. For comparison with sequence-based estimates, the 2C value (in picograms) was converted to haploid genome size (1C) by dividing by two and then converted to megabase pairs (Mbp) using 1 pg = 978 Mbp. Due to the large number of samples, analyses were performed in two batches. To assess instrument variability (run-to-run consistency), multiple technical replicates were measured for each sample. To further evaluate variability introduced by sample preparation (extraction variability) and potential batch effects, five samples from the first batch were re-extracted from their original dried tissue and re-analyzed on the second day. Adhering to standard thresholds (Greilhuber et al., 2007), G1 peak CVs were <5% and batch-to-batch estimate CVs for genome size estimates ranged from 0.11-1.98%, confirming high technical reproducibility without systematic extraction or batch effects.

**Suppl. Material (SM3):** *Principal Component Analysis*

To control for the effect of population structure on the analyses of genome size and life history traits, we performed principal component analysis (PCA) on samples. Analyses were based on genome-wide SNPs previously identified and validated in Battlay et al. (2025a). Separate PCAs were conducted for the overall analysis (439 samples) and the phenotypic analysis (221 samples). Genotype likelihoods were calculated from BAM files using ANGSD v0.939 (Korneliussen et al, 2014; https://github.com/ANGSD/angsd). Quality filters included a minimum mapping quality of Q30, minimum base quality of Q20, and a minimum minor allele frequency of 0.05, similar to parameters detailed in Battlay et al. (2025a). We modified the site inclusion criterion to require a minimum of 330 individuals with data (`-minInd 330;` 75 % of total 440 samples). Variant sites were then pruned for independence using PLINK v1.9 (Purcell et al, 2007; https://github.com/chrchang/plink-ng/blob/master/1.9/README.md), using a window size of 50 kbp, a step size of 5 SNPs, and an $r^2$ threshold of 0.5 (`-indep-pairwise 50 5 0.5`). Sites were further filtered to include only those that were both independent and outside annotated genes or haploblocks, before being randomly downsampled to 100,000 for computational efficiency. A covariance matrix was then generated from these filtered sites with PCangsd v1.2 (Meisner and Albrechtsen, 2018). The resulting PC scores were included as fixed effects in linear mixed models to account for the effect of population structure.

**Suppl. Material (SM4):** *Detail approaches for selection analysis of genome size*

**SM4a:** Genomic relatedness matrix construction, heritability and $Q_{ST}$ estimation

To estimate $Q_{ST}$, we constructed a genomic relatedness matrix (GRM) from the aforementioned VCF and implemented an animal model to estimate genetic variances in genome size. To ensure the GRM reflected genome-wide additive relatedness without bias from linked regions or rare variants, we applied the following filters: we excluded SNPs located within genes and known haploblocks (regions of high LD), filtered for a Minor Allele Frequency (MAF) ≥ 0.01, and performed LD-pruning to remove SNPs with an $r^2$ = 0.5 within a 50-kb window, using PLINK v1.9 (Purcell et al, 2007; https://github.com/chrchang/plink-ng). Following these quality and linkage filters, we randomly subsampled the remaining SNPs to a final dataset of 50,000 SNPs for GRM construction. The GRM was constructed using the VanRaden (2008) method in the R package `AGHmatrix` (Amadeu et al, 2023). To ensure mathematical stability for the animal model, the matrix was made positive-definite using the heuristic bending method in the R package `mbend` (Nilforooshan, 2020), adding a small positive value ($1 \times 10^{-6}$) to any eigenvalue smaller than $1 \times 10^{-6}$. We then modelled genome size using a Bayesian animal model in `MCMCglmm` (Hadfield, 2010) with the GRM precision matrix supplied via the `ginverse` argument, fitting genome size as a function of a grand mean intercept and an individual-level random effect whose covariance structure was informed by genome-wide SNP relatedness. A parameter-expanded prior was used for variance components, and the chain was run for 7,010,000 iterations with a burn-in of 10,000 and a thinning interval of 7,000, yielding 1,000 posterior samples. Autocorrelations between MCMC samples were below the recommended level of 0.1, yielding high effective sample sizes (mostly ≥ 1,000, minimum = 549) for all estimates. We inspected plots of traces and posterior distributions to ensure that models converged. Other priors were explored and gave similar results to those presented here. Then, heritability was estimated for each of the 1,000 MCMC iterations as:

$$h^2 = \frac{V_A}{V_A + V_R}$$

,where $V_A$ is the posterior additive genetic variance (animal component) and $V_R$ is the residual variance (units component) extracted from the VCV matrix of the fitted MCMCglmm object (Falconer & Mackay, 1996; Lynch & Walsh, 1998; de Villemereuil, 2018; Perrier C et al, 2018). Individual genomic breeding values were extracted directly from the posterior distribution of random effects stored in the Sol matrix of the fitted MCMCglmm object (Hadfield, 2010; Wilson et al, 2010). For each of the 1,000 retained posterior samples, the full vector of individual breeding values was extracted and used to calculate $Q_{ST}$ directly, without collapsing to a single point estimate. Within each geographical range (North America, Europe, Australia), breeding values were partitioned by population and $Q_{ST}$ was calculated as (Spitze, 1993; Lande, 1992; Whitlock, 2008):

$$Q_{ST} = \frac{\sigma^2_{GB}}{\sigma^2_{GB} + 2 \times \sigma^2_{GW})}$$

, where $\sigma^2_{GB}$ is the variance of population mean breeding values, used as an estimate of among-population additive genetic variance and $\sigma^2_{GW}$ the mean within-population variance of breeding values, estimating the within-population additive genetic variance across populations in that range. Applying this calculation across all 1,000 MCMC iterations produced a posterior distribution of $Q_{ST}$ for each range, from which we report the posterior mean and 95% credible interval (2.5th and 97.5th percentiles). Our approach propagates the uncertainty of breeding value estimates sampled from the posterior distribution through to the final $Q_{ST}$ estimate and is more rigorous than approaches based on a single point estimate of heritability, as recommended by Leinonen et al. (2013) and Whitlock (2008).

**SM4b:** $P_{ST}$-$F_{ST}$ comparisons for genome size differentiation

To complement the $Q_{ST}$-$F_{ST}$ analysis, we implemented a $P_{ST}$-$F_{ST}$ comparison as an additional line of evidence for divergent selection on genome size (Spitze, 1993; Prout et al, 1993) and calculated $P_{ST}$

for genome size separately for populations within Australia, Europe, and North America. Following the approach by Sæther et al. (2007) and Whitlock (2008), we estimated $P_{ST}$ as

$$P_{ST} = \frac{(VP, among)}{(VP, among) + 2(VP, within)}$$

where *VP, among* is the among-population variance and *VP, within* is the within-population variance for the trait. Variance components were estimated using linear-mixed models implemented in R\ lme4 (Bates et al, 2015), by fitting the trait (genome size) as the response and population as a random intercept. Genome size was measured from individuals grown in a common-garden experiment, thereby minimising environmental sources of variation. Heritability of genome size was estimated at 0.886 (95% CI: 0.772–0.953; Fig. 5b) from the Bayesian animal model described above, confirming that the vast majority of phenotypic variance reflects additive genetic differences among individuals. Consequently, $P_{ST}$ is expected to closely approximate $Q_{ST}$ (Whitlock, 2008; Brommer, 2011), and any residual non-additive transmission processes would bias $P_{ST}$ downward, making our estimates conservative with respect to detecting genetic differentiation.

To assess whether lower within-population sample sizes could inflate $P_{ST}$ estimates, we generated 2,000 permuted datasets by randomly assigning individuals to populations while preserving overall sample sizes. $P_{ST}$ values from observed data were then compared against this null distribution. A $P_{ST}$ value exceeding the 99$^{th}$ percentile of the neutral $F_{ST}$ distribution was interpreted as evidence of trait divergence exceeding neutral expectations, consistent with divergent selection acting on the trait. Following Sun et al. (2020) and Brommer (2011), we also performed $P_{ST}$ sensitivity across a range of c values (0.1–1.5), where c represents the proportion of total additive genetic variance attributable to among-population differences and heritability values ($h^2$ = 0.25, 0.50, 0.75).

**Result:**

In concordance with $Q_{ST}$-$F_{ST}$ analysis, $P_{ST}$-$F_{ST}$ analysis revealed that genome size differentiation significantly exceeded neutral expectations in North America ($P_{ST}$ = 0.363 > $F_{ST}$ threshold = 0.257 ) and Europe, $P_{ST}$ = 0.454 > $F_{ST}$ threshold = 0.277) (Fig. 1), providing strong evidence that divergent selection is driving genome size differences among populations in these ranges. Sensitivity analysis further confirmed the robustness of the selection signatures in North America and Europe, as $P_{ST}$ remained significantly higher than the neutral $F_{ST}$ threshold across all tested $h^2$ values at biologically realistic c values (Fig. 2). In contrast, populations in Australia exhibited almost no differentiation in genome size. The observed $P_{ST}$ was 0, which fell well below the neutral $F_{ST}$ thresholds, consistent with an absence of divergent selection or the influence of stabilizing selection within this range.

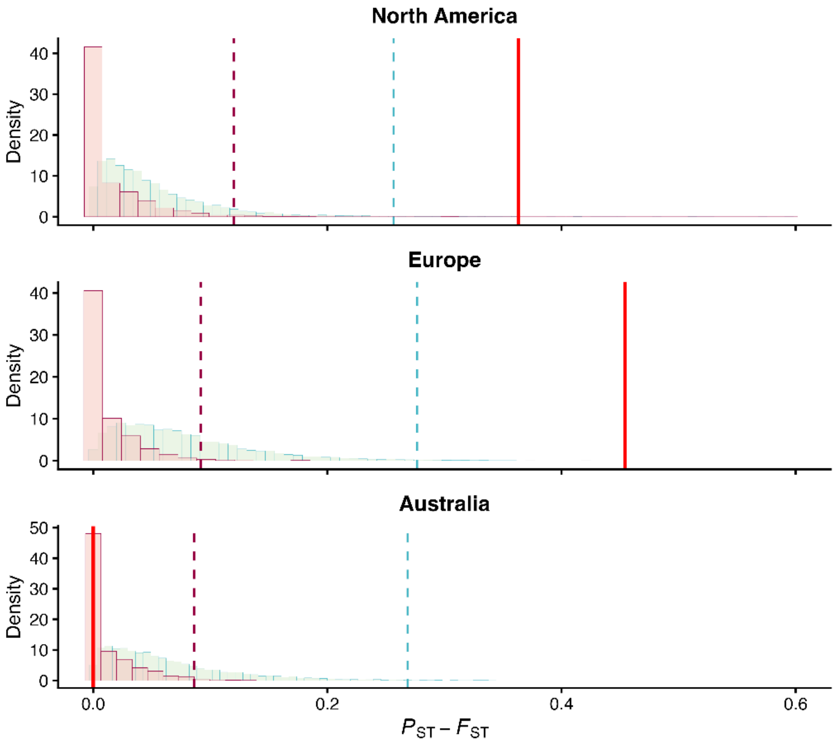


**Fig. 1: Phenotypic differentiation ($P_{ST}$) compared to neutral genetic differentiation ($F_{ST}$) for North America, Europe, and Australia.** The light green background histograms represent the null distribution of 10,000 putatively neutral $F_{ST}$ markers, with the teal dashed line indicating the 99th percentile of this distribution. Pink histograms represent the null distribution generated from 2,000 permutations. The vertical dashed pink line indicates the 99th percentile of the permutation null, while the solid red line indicates the observed $P_{ST}$ value.

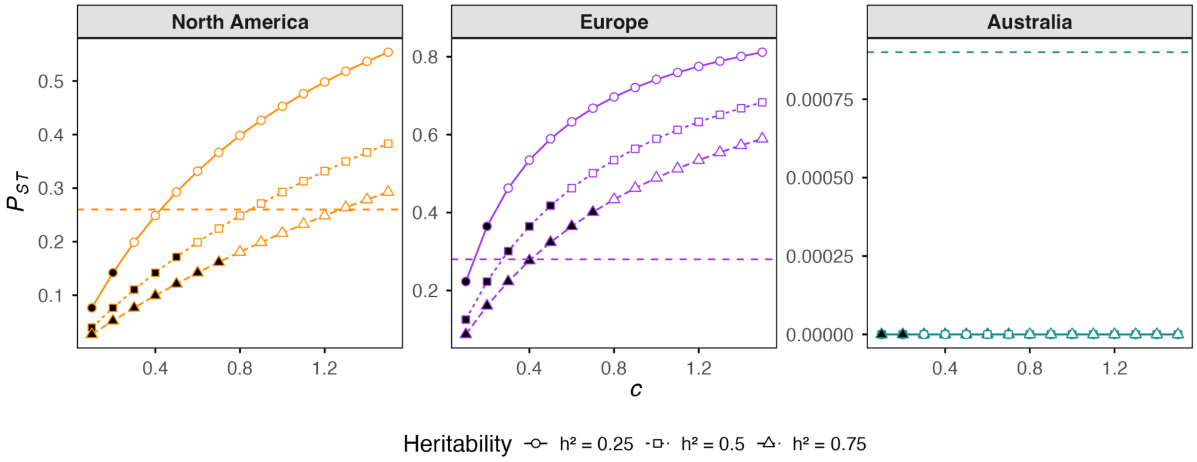


**Fig. 2: $P_{ST}$–$F_{ST}$ sensitivity analysis** across ranges. $P_{ST}$ values are plotted across a range of c values for three heritability assumptions ($h^2$ = 0.25, 0.5, 0.75). The horizontal dashed line indicates the $F_{ST}$ threshold (99th percentile for North America and Europe; 1st percentile for Australia). Filled symbols indicate biologically informative parameter combinations where c ≤ $h^2$; open symbols indicate uninformative combinations where c > $h^2$. Filled symbols exceeding the dashed line indicate $P_{ST} > F_{ST}$

under conservative assumptions, consistent with divergent selection; filled symbols below the threshold indicate differentiation below neutral expectations.

**SM4c:** $Q_{PC}$ framework for testing divergent selection of genome size

To complement the $P_{ST}$–$F_{ST}$ and $Q_{ST}$–$F_{ST}$ comparisons, we additionally applied the $Q_{PC}$ framework (Josephs et al, 2019) to test for divergent selection on genome size while explicitly accounting for genome-wide population structure. Unlike $P_{ST}$–$F_{ST}$ and $Q_{ST}$–$F_{ST}$ comparisons, which treat populations as discrete units, $Q_{PC}$ projects the phenotype onto the eigenvectors of the kinship matrix to identify axes of population structure along which trait divergence exceeds neutral expectations. Following Josephs et al. (2019), a kinship matrix (K) was constructed from the same 50,000 randomly sampled SNPs used for the $Q_{ST}$ analysis, after excluding sites with missing data, using the VanRaden (2008) method implemented in the R/AGHmatrix applied to 289 individuals. Unlike the GRM used in the Bayesian animal model, the kinship matrix was not subjected to positive-definiteness correction, preserving the full eigenspectrum required for $Q_{PC}$. The K matrix was eigendecomposed to obtain principal components (PCs) and their associated eigenvalues. The PC cutoff (pcmax = 52) was defined as the first PC at which cumulative variance explained exceeded 30% of total genetic variance, following Josephs et al. (2019) who used the first PCs cumulatively explaining 30% of variation in K.

For each invasion range independently (North America, Europe, and Australia), genome size was mean-centred and projected onto the eigenvectors of K. For each of the top 52 PCs, the $Q_m$ statistic was computed as:

$$Q_m = \frac{(Z.u_m)^2/\lambda_m}{\frac{1}{144}\sum_{i=146}^{289}(Z.u_i)^2/\lambda_i}$$

Where Z is the mean-centred vector of genome size values for individuals within a given range, $u_m$ is the $m^{th}$ eigenvector of K, and $\lambda_m$ is the corresponding eigenvalue. The numerator captures among-population divergence in genome size along $PC_m$ ($\sigma^b$), and the denominator estimates neutral within-population variance ($\sigma^w$) from the bottom 144 PCs ($\frac{n}{2}$, where n = 289, following Josephs et al, 2019). Under neutrality, $Q_m$ follows an F distribution with 1 and 144 degrees of freedom, from which p-values were derived. A significant positive $Q_m$ indicates that trait divergence along a given PC axis exceeds neutral expectations. For display purposes, 95% confidence intervals were plotted as lines with slope $\pm 1.96\sqrt{(V_a \times \lambda_m)}$ per $PC_m$, where $V_a$ is the neutral additive variance estimated from the bottom 144 PCs, following Josephs et al. (2019). The significance threshold applied was $p < 0.05$.

**Result:**

$Q_{PC}$ analysis provided convergent evidence with the $P_{ST}$–$F_{ST}$ and $Q_{ST}$–$F_{ST}$ results, confirming strong directional selection on genome size in North America, weak signal in Europe, and no signal in Australia (Fig. 1).

In North America, 15 out of 52 tested PCs showed significant associations between genome size and PC score (P < 0.05), with the strongest signal at PC12 ($Q_m$ = 20.28, P = 1.37×10$^{-5}$; Fig. 2). Across all significant PCs, the observed regression of genome size on PC score consistently and substantially exceeded the neutral 95% confidence interval (Fig. 2), indicating that genome size divergence among North American populations is far greater than expected under genetic drift alone. In Europe, three PCs reached the significant threshold (PC17, PC22, PC45; P < 0.05), with observed projections marginally exceeding the neutral confidence interval. In Australia, no PCs showed significance (all P > 0.12), with observed projections falling entirely within the neutral confidence interval (Fig. 2), indicating no detectable divergent selection on genome size.

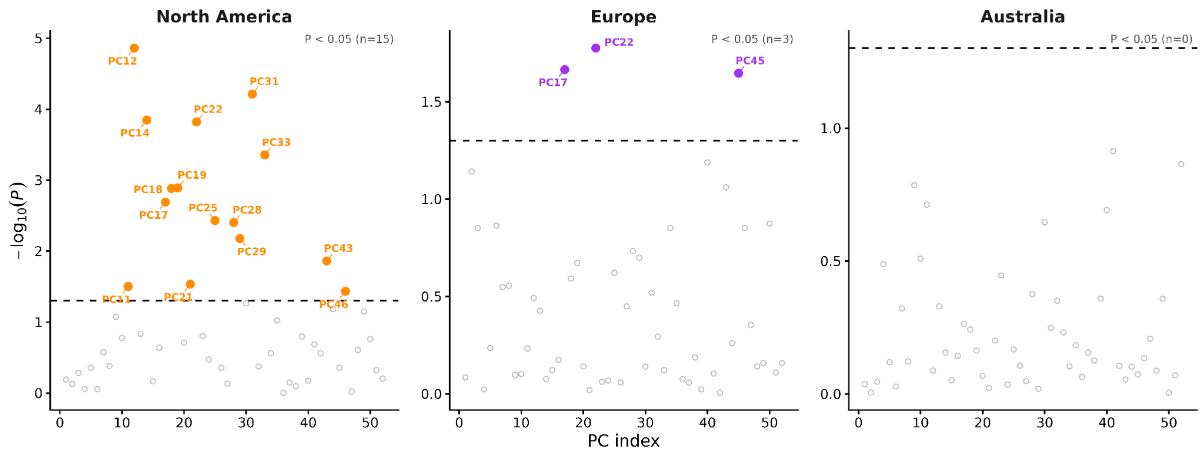


**Fig. 1: $Q_{PC}$ projections of genome size onto principal component axes of the kinship matrix across geographical ranges.** The y-axis shows −log$_{10}$(P-value); the dashed horizontal line marks the significance threshold (P = 0.05). Points above the threshold (P < 0.05) are shown in range-specific colour; non-significant points are shown in grey, with PC labels shown for significant points. North America: 15 significant PCs. Europe: 3 significant PCs. Australia: no significant PCs.

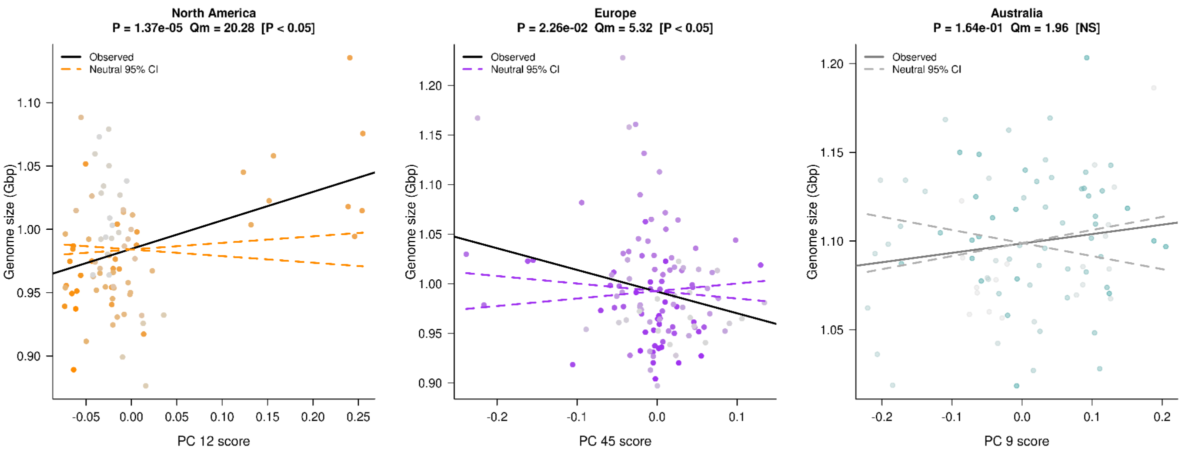


**Fig. 2: *$Q_{PC}$* projections of genome size onto one PC axis per range.** Genome size (Gbp) is plotted against individual scores on the PC axis within each range: North America ($PC_{12}$, $P = 1.37\times10^{-5}$), Europe ($PC_{45}$, $P = 0.023$), and Australia ($PC_9$, $P = 0.164$). The solid line shows the observed regression of genome size on PC score. Dashed lines indicate the expected regression slope under neutrality, derived from drift-based variance estimates ($\pm1.96\sqrt{(Va \times \lambda m)}$); slopes falling outside this envelope indicate divergence beyond neutral expectations. In North America and Europe, the observed regression exceeds the neutral envelope, consistent with divergent selection on genome size. In Australia, the observed regression falls within the neutral envelope, indicating genome size variation consistent with neutral genetic drift.

**Suppl. Material (SM5):** *Polygenic value estimation for flowering time*

Using flowering onset GWAS results from Battlay et al. (2025a), LD clumping was performed in PLINK to identify a single SNP per GWAS peak in each LD block. We considered only SNPs with significant association (Bonferroni-corrected $P < 0.05$), and 'clumped' candidate SNPs in plink v. 1.9 to remove SNPs in LD ($r^2 > 0.3$) with the top candidate in each 1 Mbp window. Following clumping, 380 of 513 associated SNPs were retained. We then inferred genotypes for each of the 380 sites across our samples using ANGSD. We estimated the genotype likelihoods using the GATK model (`-GL 2`), with major and minor allele defined from the candidate SNPs list (`-doMajorMinor 4`) for each site. Reads with mapping quality below $Q_{30}$ were removed (`-minMapQ 30`), only bases with quality scores above $Q_{20}$ were retained (`-minQ 20`), and a minor allele frequency filter of 0.05 was applied (`-minMaf 0.05`). Allele frequencies were estimated assuming a known major allele, with the minor allele frequency inferred from the data (`-doMaf 2`) and only sites with a probability of being variable less than $1 \times 10^{-6}$ were retained (`-SNP_pval 1e-6`). We then used genotype data along with their effect sizes from the GWAS, to estimate the polygenic value for flowering onset ($PGV_{FT}$) of each genotyped individual as described in Kreiner et al. (2023):

$$PGV_{FT} = 2\sum_{i=1}^{L} \alpha p$$

where $L$ represents the set of 380 LD-pruned loci with bonferroni-corrected $P < 0.05$, $\alpha$ is the effect size of the minor allele at locus $i$ derived from the GWAS, and $p$ is the individual's genotype frequency at that locus (coded as 0 for homozygous reference, 0.5 for heterozygotes, and 1 for homozygous alternates).

**Suppl. Material (SM6):** *Identification of primary climatic predictor*

To identify informative environmental predictors of genome size, we first extracted 19 bioclimatic variables (BIO1–BIO19, Suppl. Table S10) from WorldClim (Fick & Hijmans, 2017) for each sampling location using geographic coordinates (latitude and longitude) and year of sample collection. We assessed the potential for redundancy by calculating pairwise Pearson correlation coefficients across all 19 variables (Suppl. Fig. S13,a). Highly correlated variables (absolute Pearson's r > 0.7) were identified and manually reduced by retaining a single representative variable from each correlated cluster. From the resulting uncorrelated clusters, five variables were retained: BIO1 (Mean Annual Temperature, MAT) representing the temperature-related cluster (BIO3–BIO7, BIO9–BIO11); BIO2 (Mean Diurnal Range); BIO8 (Mean Temperature of Wettest Quarter); BIO12 (Annual Precipitation) representing the precipitation-related cluster (BIO13, BIO14, BIO16–BIO19); and BIO15 (Precipitation Seasonality). Then, PCA was subsequently performed on these five variables to quantify the environmental variance explained by each component (Suppl. Fig. S13b). The first principal component explained 48% of the total variance and was strongly correlated with MAT (r = 0.84) (Suppl. Fig. S13,b,c), which has established biological importance for common ragweed. Accordingly, MAT was selected as the primary climatic predictor for subsequent analyses.

**Suppl. Material (SM7):** *Testing the Large Genome Constraint hypothesis*

To test whether genome size constrains developmental speed consistent with the LGC hypothesis, we first fitted a linear mixed model of flowering time on genome size using the `lme4` R package (Bates et al, 2015), controlling for range, polygenic value for flowering time ($PGV_{FT}$), PC1, PC2, and population as a random intercept. This revealed no significant effect of genome size on mean flowering time. Because the LGC hypothesis specifically predicts a lower boundary constraint; larger genomes restrict the minimum time required for development, rather than shifting the mean. Thus, standard mean-based regression is not well suited to detect this signal. We therefore fitted Linear Quantile Mixed Models (LQMM) using the `lqmm` R package (Geraci, 2014) at five quantiles ($\tau$ = 0.10, 0.25, 0.50, 0.75, and 0.90), using data from 221 individuals (van Boheemen et al, 2019). The lower quantiles ($\tau$ = 0.10, 0.25) were of primary interest as they captured the minimum developmental boundary predicted by the LGC hypothesis, while higher quantiles served as comparators to assess whether any effect was specific to the lower tail of the distribution. The fitted R models are:

Across all ranges,

Phenotypic trait (flowering time) ~ Genome size + Range + $PGV_{FT}$ + PC1 + PC2 + (1 | Population)

Within each range,

Phenotypic trait (flowering time) ~ Genome size + $PGV_{FT}$ + PC1 + PC2 + (1 | Population) where Genome size, $PGV_{FT}$, PC1 and PC2 were treated as fixed effects as previously described, and Population as a random intercept. This framework allowed us to isolate the effect of genome size on developmental thresholds while controlling for the additive genetic component of flowering time and population structure.

### B) Supplementary Figures:

**Suppl. Fig. S1: Distribution of genome size estimated by sequence-based method, with mean and standard deviations.**

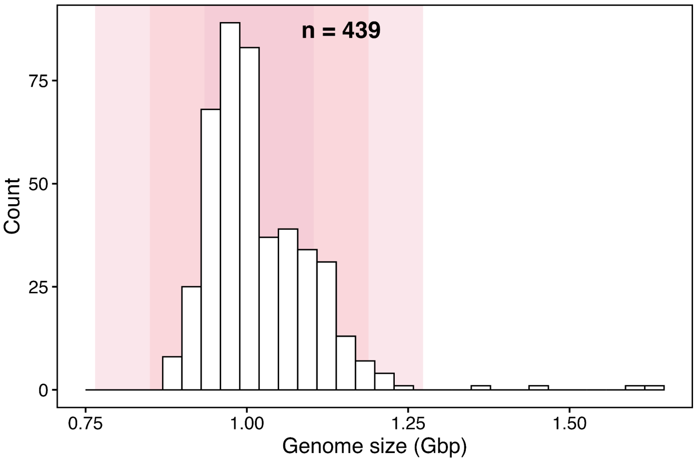


**Suppl. Fig. S2: Assessment of sequencing coverage bias on genome size estimates.**

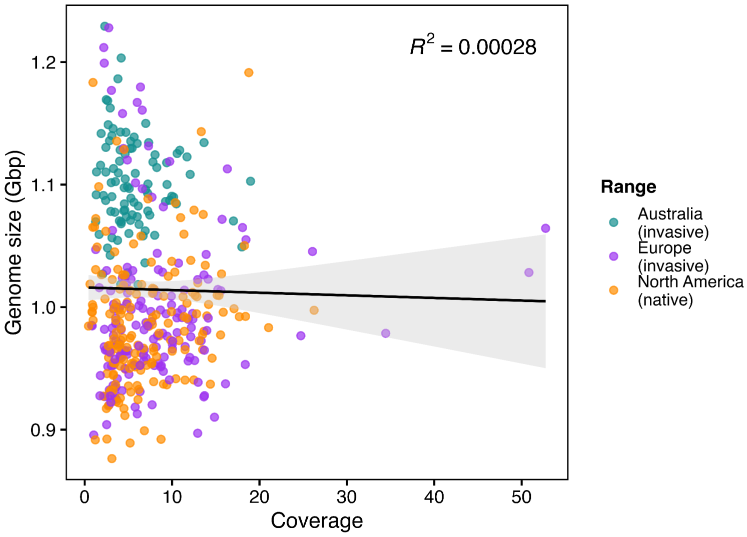


**Suppl. Fig. S3: Relationships between estimated genome size and various genomic metrics.** Scatter plots depict the relationship between: (a) estimated genome size and insert size; (b)

estimated genome size and read length; (c) callable depth and average whole genome depth; (d) estimated genome size and genome size calculated with depth of variance-based filtered BUSCO sites; and (e) estimated genome size and clonality. In these plots, each point represents a sample; the dark blue line shows the regression, with a light blue shaded area indicating the 95% confidence interval. (f) Genome size box plot estimated from sequence data including duplicate reads, to check the consistency of the pattern with estimates without duplicates (Fig. 2B). Above the boxes, we provide sample size (n), and statistical values (model's estimated marginal mean ± standard error). Significance codes: *** < 0.001, ** < 0.01, * < 0.05, ns = not significant.

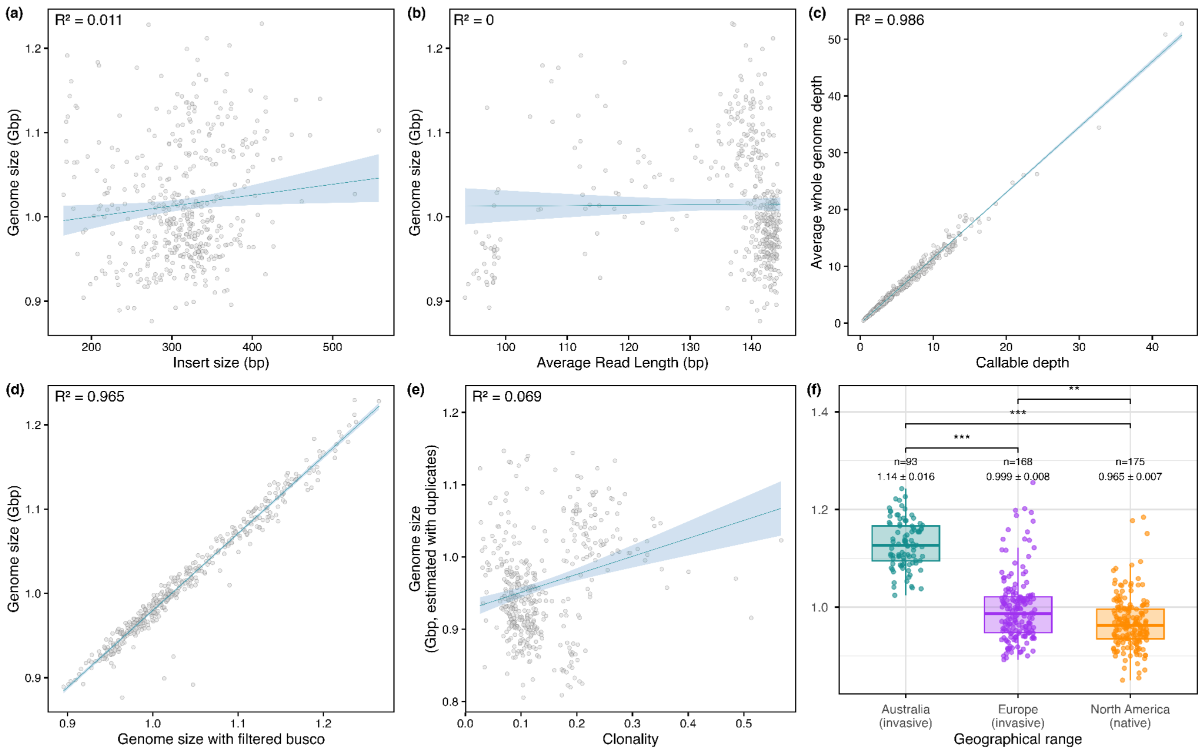


**Suppl. Fig. S4: Relationship between genome size with read mapping quality filters (Q2,5,7,10,15) and genome size without filters.**

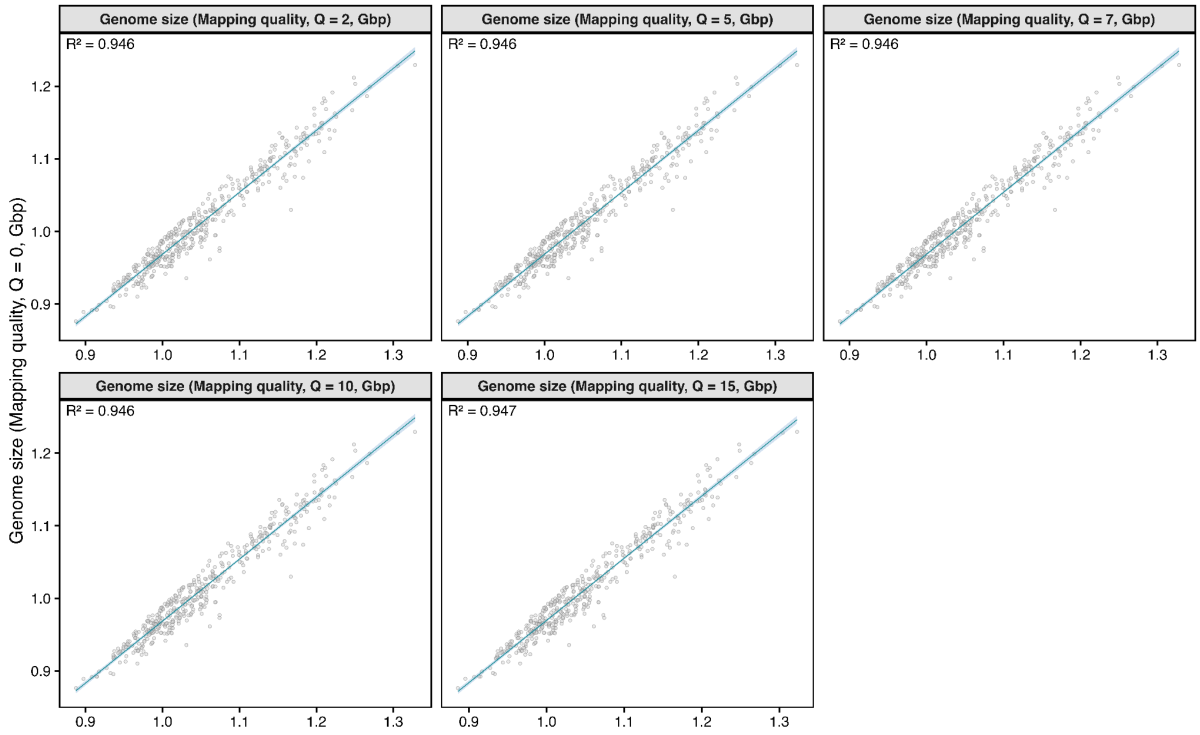


**Suppl. Fig. S5: Estimation and comparison of genome size by flow cytometry.** Genome size was estimated for (a) cassava sample using tomato (*Stupické polní rané*) as an internal standard, and (b) common ragweed using cassava as the internal control. The zoomed-out plots show the histograms of the corresponding gated nuclei populations.

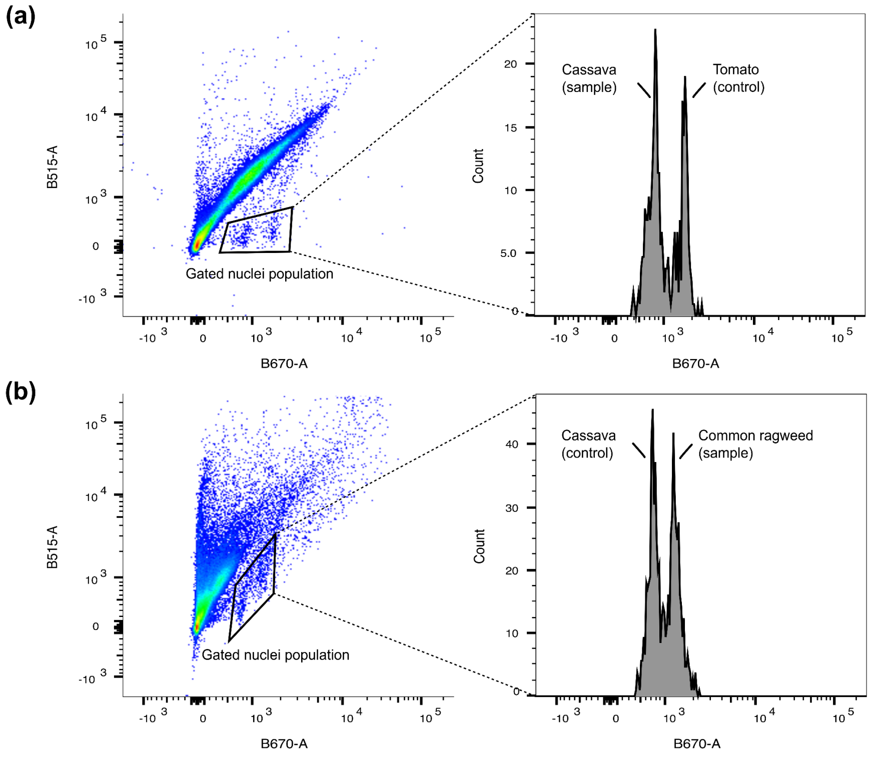

**Suppl. Fig. S6: Distribution of haploid genome size (Gbp) estimated by flow-cytometry.**

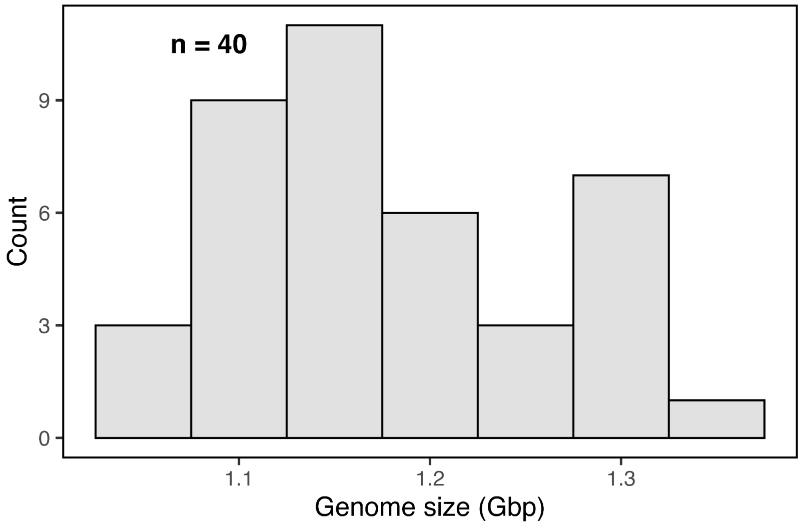


**Suppl. Fig. S7: Comparison between the sequence and flow-cytometry-based genome size estimates** (collected from different samples and times, but from the same locations [Suppl. Table S2]).

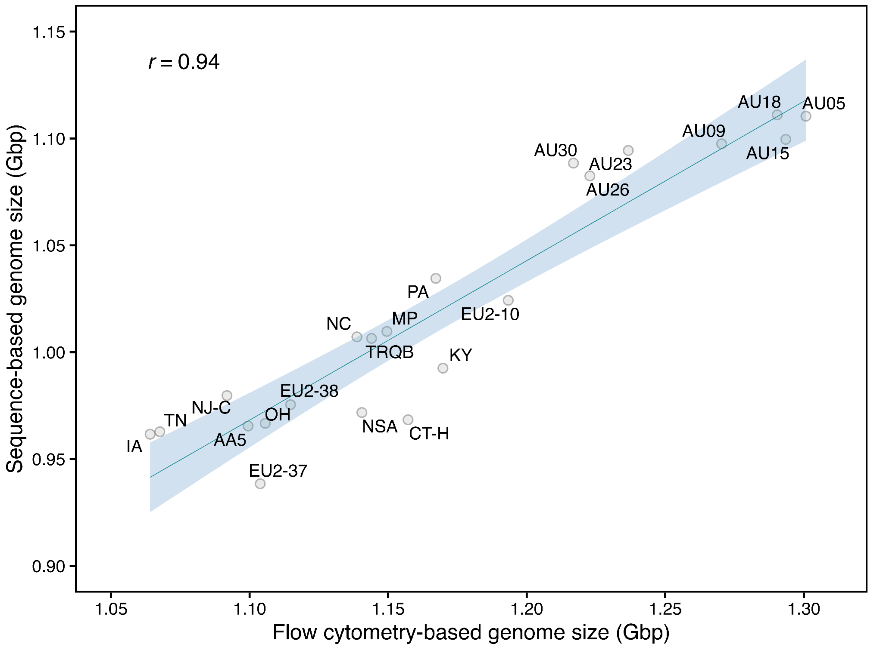


**Suppl. Fig. S8: Assessment of sequencing coverage bias on TE abundance estimates.**

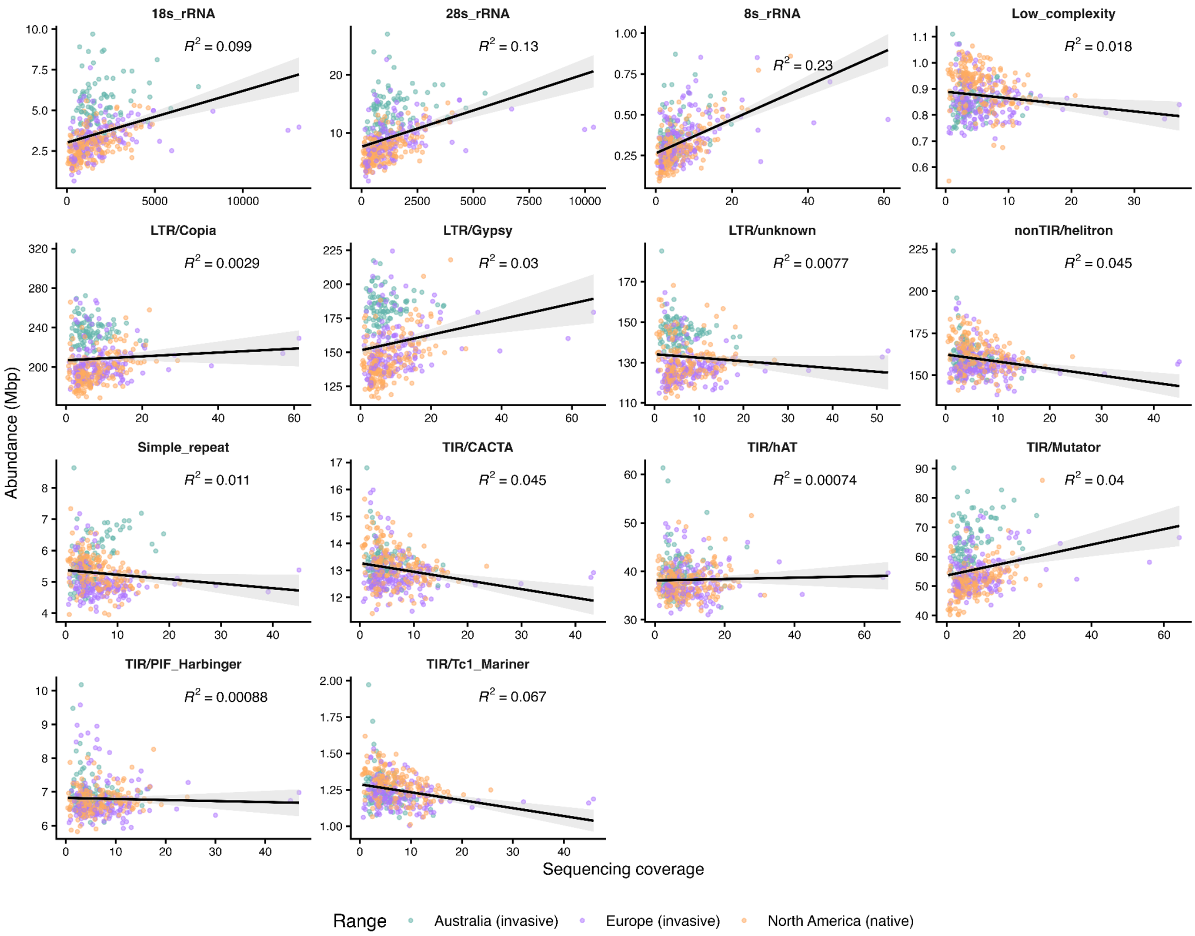


**Suppl. Fig. S9: Comparison of TEs and rRNAs abundances between invasive ranges and their native populations.** Box plots show TE/rRNAs abundance (Mbp) for Australia (AUS), Europe (EUR), North America spatio-genetic clusters mid-east (NAM.ME), south (NAM.S), west (NAM.W) and east (NAM.E). Asterisks denote statistical significance from each pairwise comparisons (e.g., **P* < 0.05, ***P* < 0.01, ****P* < 0.001), and 'ns' indicates non-significant differences.

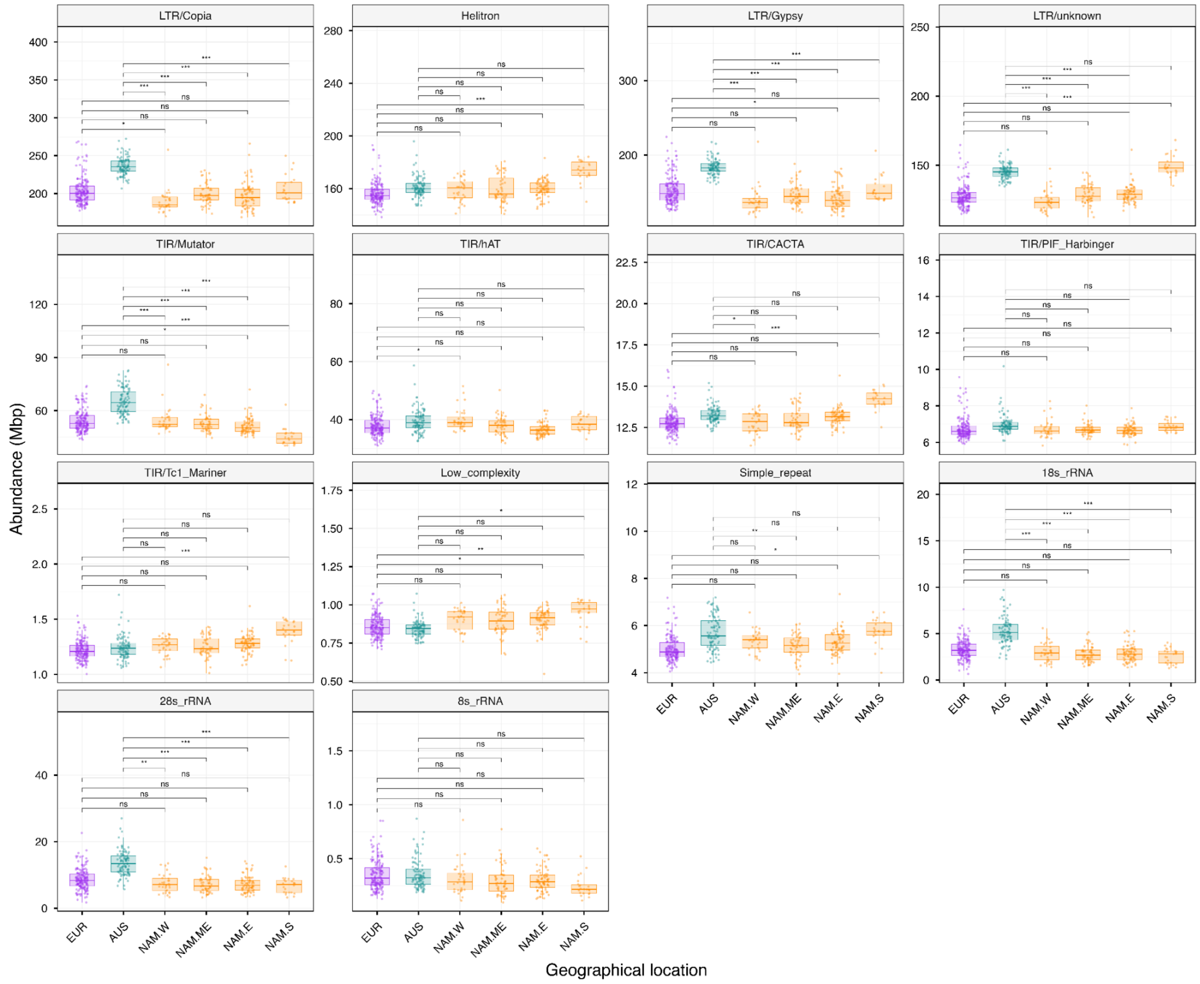


**Suppl. Fig. S10: Comparison of genome size between two Australian clusters (AU01 and rest) and their native populations.** Box plots show genome size (Gbp) for two Australian clusters (AU01 and AUrest), and North America spatio-genetic clusters [mid-east (NAM.ME), south (NAM.S), west (NAM.W) and east (NAM.E)]. Sample sizes (n) and estimated mean genome sizes ± standard error are provided below each box plot. Asterisks denote statistical significance from pairwise comparisons (e.g., **P* < 0.05, ***P* < 0.01, ****P* < 0.001), and 'ns' indicates non-significant differences.

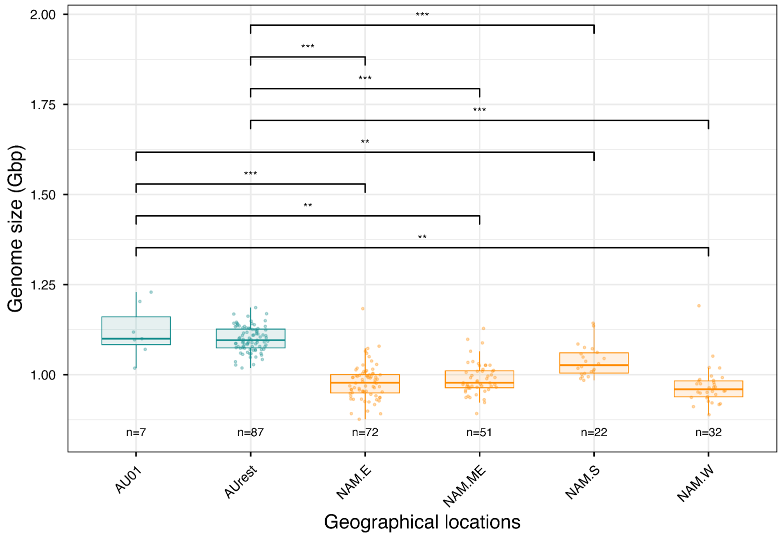


**Suppl. Fig. S11: Proportion of different TE and rRNA families across the individuals from native and invasive ranges.**

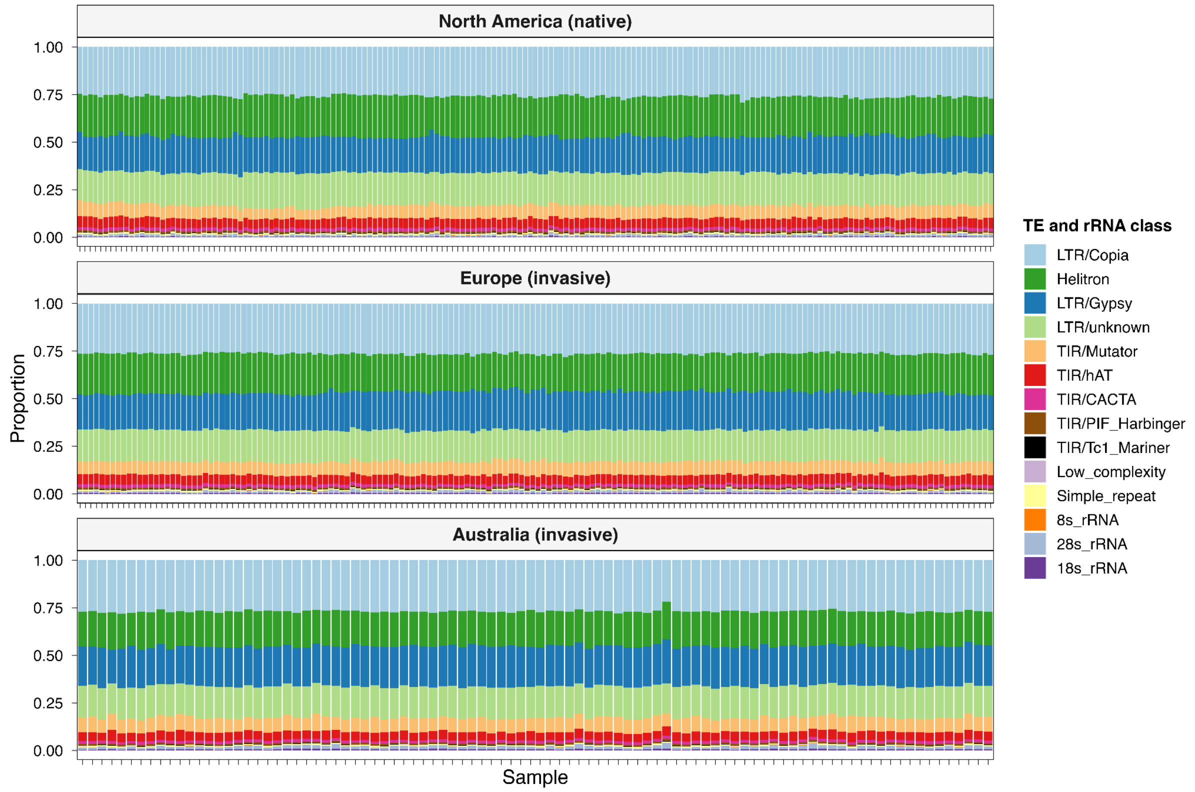


**Suppl. Fig. S12: Abundance of different TE families across geographic ranges**. Asterisks denote statistical significance from pairwise comparisons (e.g., *$P$ < 0.05, **$P$ < 0.01, ***$P$ < 0.001), and 'ns' indicates non-significant differences.

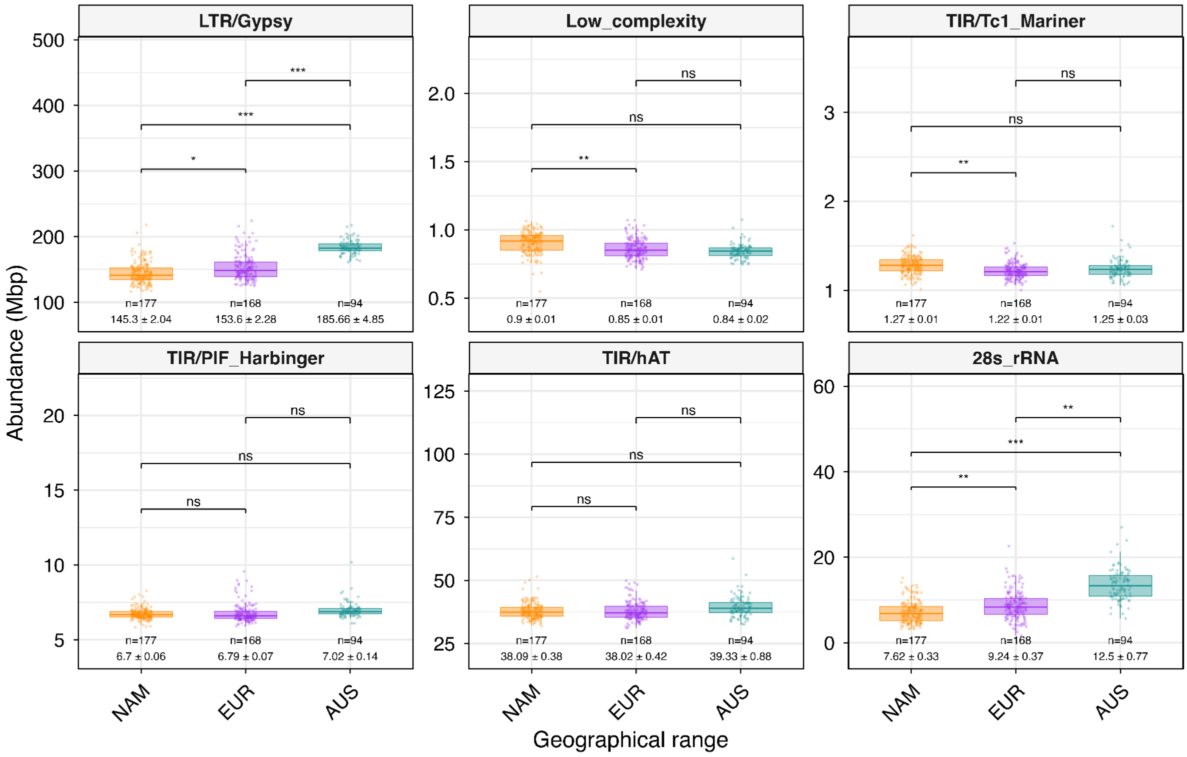


**Suppl. Fig. S13: Identification of primary climatic predictors associated with common ragweed populations.** a) Pearson correlation matrix among 19 bioclimatic variables showing strong collinearity among temperature- and precipitation-related variables. b) Principal Component Analysis (PCA) based on selected variables (R < 0.70) and Mean Annual Temperature (MAT), illustrating the relative contribution of each variable to the first two principal components. c) Pearson correlation between mean annual temperature (MAT) and the first principal component (PC1), showing a strong association (r = 0.84) between MAT and the dominant climatic gradient.

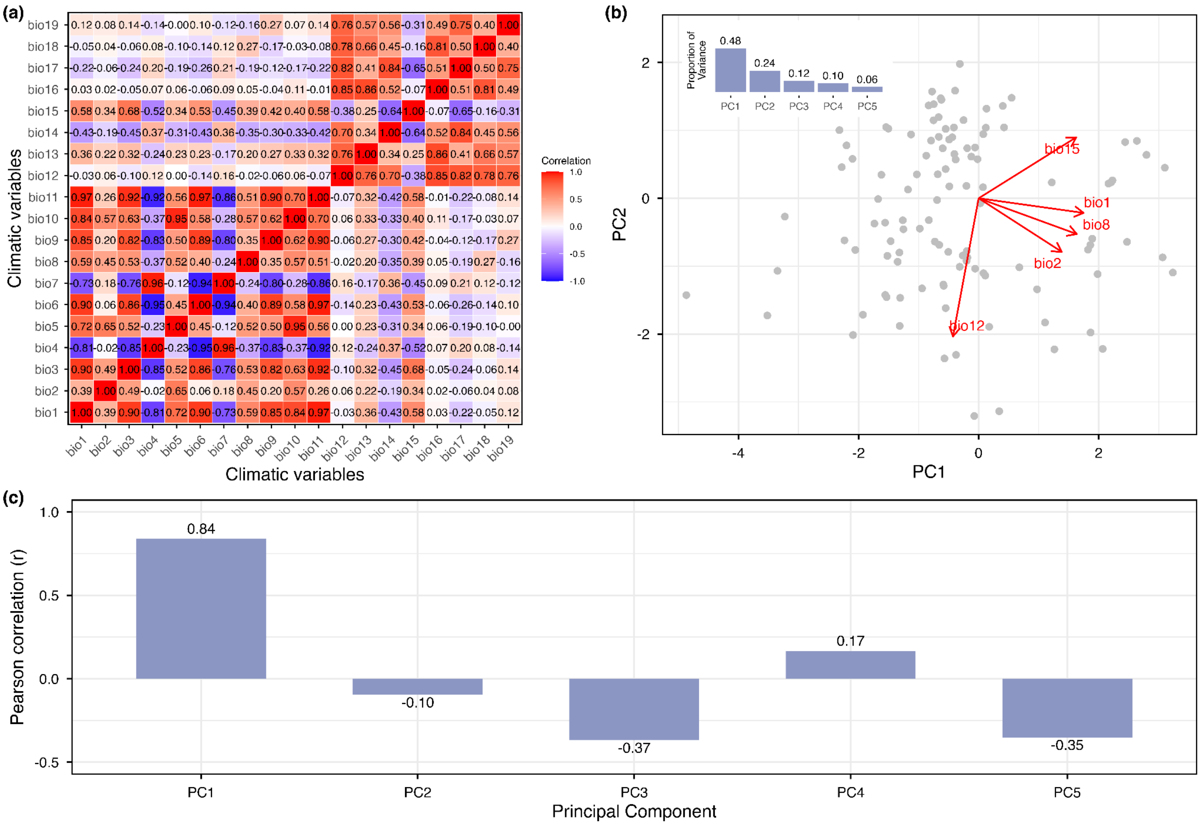


### C) Supplementary Tables:

**Suppl. Table S1: Summary statistics of rRNA and TE annotations,** showing the number of sites, length (mean ± standard error) and distance to nearest gene (mean ± standard error) for each family.

| rRNA and TEs | Number of sites | Total length (Mbp) | Mean_length ± SE (bp) | Mean distance to nearest gene ± SE (Mbp) |
|---|---|---|---|---|
| TIR/PIF_Harbinger | 22782 | 7.3997 | 325.49 ± 2.63 | 0.0489 ± 0.001 |
| Helitron | 473277 | 173.8820 | 371.29 ± 0.83 | 0.0234 ± 0.0001 |
| Low_complexity | 22686 | 1.1171 | 49.31 ± 0.19 | 0.0173 ± 0.0004 |
| LTR/unknown | 201031 | 128.1834 | 637.84 ± 2.33 | 0.0404 ± 0.0001 |
| LTR/Gypsy | 105311 | 134.4367 | 1276.75 ± 6.17 | 0.0447 ± 0.0003 |
| TIR/hAT | 87681 | 29.3837 | 335.31 ± 2.09 | 0.0169 ± 0.0002 |
| LTR/Copia | 154880 | 185.3814 | 1197.04 ± 4.73 | 0.053 ± 0.0003 |
| TIR/Mutator | 133168 | 51.3472 | 385.85 ± 2.57 | 0.1923 ± 0.0009 |
| TIR/CACTA | 44239 | 14.7256 | 333.98 ± 2.15 | 0.0254 ± 0.0004 |
| TIR/Tc1_Mariner | 4859 | 1.2810 | 265.07 ± 3.84 | 0.0156 ± 0.0005 |

| | | | | |
|---|---|---|---|---|
| Simple_repeat | 131708 | 5.9165 | 44.99 ± 0.41 | 0.0176 ± 0.0002 |
| 8s_rRNA | 3324 | 0.3823 | 115 ± 0.01 | 0.2911 ± 0.0026 |
| 18s_rRNA | 9 | 0.0149 | 1652.22 ± 47.04 | 0.03 ± 0.0078 |
| 28s_rRNA | 10 | 0.0523 | 5232.5 ± 285.5 | 0.0263 ± 0.0065 |

**Suppl. Table S2: Flow cytometry–based genome size measurements for 40 common ragweed samples.** Genome sizes in gigabase pairs (Gbp) were calculated by converting picograms (pg) to Gbp using a factor of 0.978 (1 pg DNA = 0.978 Gbp).

| Sample | Population | Range | 2C (pg) | 2C (Gbp) | 1C (Gbp) |
|---|---|---|---|---|---|
| NSA-1-4 | NSA | North America | 2.2207 | 2.1718 | 1.0859 |
| NC-10-2 | NC | North America | 2.3287 | 2.2774 | 1.1387 |
| OH-6-6 | OH | North America | 2.2631 | 2.2133 | 1.1067 |
| NSA-13-2 | NSA | North America | 2.444 | 2.3903 | 1.1951 |
| OH-2-4 | OH | North America | 2.2802 | 2.23 | 1.115 |
| AA5-2-1 | AA5 | North America | 2.2484 | 2.199 | 1.0995 |
| KY-1-1 | KY | North America | 2.3922 | 2.3396 | 1.1698 |
| TRQB-3-1 | TRQB | North America | 2.3395 | 2.288 | 1.144 |
| CTH-2-1 | CT-H | North America | 2.3664 | 2.3144 | 1.1572 |
| IA-7-1 | IA | North America | 2.1761 | 2.1282 | 1.0641 |
| MI-3-4 | MI | North America | 2.198 | 2.1497 | 1.0748 |
| MP-4-2 | MP | North America | 2.3508 | 2.2991 | 1.1495 |
| OH-10-1 | OH | North America | 2.2398 | 2.1905 | 1.0953 |
| PA-5-2 | PA | North America | 2.3869 | 2.3344 | 1.1672 |
| TN-7-3 | TN | North America | 2.1832 | 2.1352 | 1.0676 |
| NJC-11-1 | NJC | North America | 2.2328 | 2.1836 | 1.0918 |
| EU38-13-1 | EU2-38 | Europe | 2.2797 | 2.2296 | 1.1148 |
| EU10-6-4 | EU2-10 | Europe | 2.4011 | 2.3483 | 1.1741 |
| EU37-4-4 | EU2-37 | Europe | 2.2467 | 2.1973 | 1.0986 |
| EU37-8-2 | EU2-37 | Europe | 2.2679 | 2.2181 | 1.109 |
| EU10-10-4 | EU2-10 | Europe | 2.3978 | 2.345 | 1.1725 |

| | | | | | |
|---|---|---|---|---|---|
| EU10-3-1 | EU2-10 | Europe | 2.4796 | 2.425 | 1.2125 |
| AU15-4-1 | AU15 | Australia | 2.7173 | 2.6575 | 1.3288 |
| AU26-2-3 | AU26 | Australia | 2.5034 | 2.4484 | 1.2242 |
| AU23-3-1 | AU23 | Australia | 2.6446 | 2.5864 | 1.2932 |
| AU35-1-2 | AU35 | Australia | 2.3213 | 2.2702 | 1.1351 |
| QLD4-8-2 | QLD4 | Australia | 2.3407 | 2.2892 | 1.1446 |
| AU9-6-6 | AU9 | Australia | 2.6075 | 2.5501 | 1.275 |
| AU1-1-2 | AU15 | Australia | 2.6651 | 2.6064 | 1.3032 |
| AU5-4-2 | AU5 | Australia | 2.66 | 2.6015 | 1.3008 |
| AU9-2-5 | AU9 | Australia | 2.5703 | 2.5137 | 1.2569 |
| AU9-9-3 | AU9 | Australia | 2.6157 | 2.5582 | 1.2791 |
| AU15-2-3 | AU15 | Australia | 2.6003 | 2.5431 | 1.2716 |
| AU15-6-4 | AU15 | Australia | 2.5979 | 2.5408 | 1.2704 |
| AU18-2-5 | AU18 | Australia | 2.6085 | 2.5511 | 1.2755 |
| AU18-3-1 | AU18 | Australia | 2.6693 | 2.6106 | 1.3053 |
| AU23-7-5 | AU23 | Australia | 2.4133 | 2.3602 | 1.1801 |
| AU26-5-6 | AU26 | Australia | 2.4978 | 2.4428 | 1.2214 |
| AU30-3-4 | AU30 | Australia | 2.4885 | 2.4337 | 1.2169 |
| QLD5-5-5 | QLD5 | Australia | 2.3424 | 2.2909 | 1.1454 |

**Suppl. Table S3: Genome size comparison between invasive ranges (Australia, AUS; Europe, EUR) and native populations from North America (NAM.ME = mid-east, NAM.S = south, NAM.W = west, NAM.E = east).** Significance codes: **p < 0.001, *p < 0.01, p < 0.05, ns = not significant.

| Comparison | Estimate | SE | df | t.ratio | p.value | Sig. |
|---|---|---|---|---|---|---|
| EUR vs NAM.W | 0.0315 | 0.0194 | 52.48 | 1.63 | 1 | ns |
| EUR vs NAM.ME | 0.0124 | 0.013 | 83.88 | 0.95 | 0.343 | ns |
| EUR vs NAM.E | 0.028 | 0.0146 | 80.53 | 1.91 | 0.06 | ns |
| EUR vs NAM.S | –0.0342 | 0.0221 | 70.18 | –1.54 | 0.127 | ns |
| AUS vs NAM.W | 0.1437 | 0.0149 | 31.72 | 9.67 | $5.68 \times 10^{-11}$ | *** |
| AUS vs NAM.ME | 0.1171 | 0.0125 | 37.88 | 9.41 | $1.87 \times 10^{-11}$ | *** |

| | | | | | | |
|---|---|---|---|---|---|---|
| AUS vs NAM.E | 0.1402 | 0.0139 | 31.1 | 10.12 | 2.33 x $10^{-11}$ | *** |
| AUS vs NAM.S | 0.0829 | 0.0128 | 34.35 | 6.47 | 2.02 x $10^{-7}$ | *** |

**Suppl. Table S4: Anova results of linear mixed model for TEs and rRNAs abundance with range, PC1, PC2 as fixed effects and population as random effect. Post hoc pairwise comparison for TEs and rRNA in different ranges from the mixed model. Significance codes: *** < 0.001, ** < 0.01, * < 0.05, ns = not significant**.

| TEs and rRNAs | Effect | F | Df | Df.res | p_value | FDR | Sig | FDR_sig |
|---|---|---|---|---|---|---|---|---|
| LTR/Copia | range | 38.18 | 2 | 134.28 | 7.47 x $10^{-14}$ | 1.05 x $10^{-12}$ | *** | *** |
| Helitron | range | 4.03 | 2 | 155.03 | 0.0197 | 0.0229 | * | * |
| LTR/Gypsy | range | 27.03 | 2 | 150.99 | 9.25 x $10^{-11}$ | 4.32 x $10^{-10}$ | *** | *** |
| LTR/unknown | range | 19.24 | 2 | 155.3 | 3.42 x $10^{-8}$ | 1.2 x $10^{-7}$ | *** | *** |
| TIR/Mutator | range | 32.01 | 2 | 130.43 | 4.91 x $10^{-12}$ | 3.43 x $10^{-11}$ | *** | *** |
| TIR/hAT | range | 0.76 | 2 | 120.52 | 0.4696 | 0.4696 | ns | ns |
| TIR/CACTA | range | 4.25 | 2 | 155.66 | 0.016 | 0.0204 | * | * |
| TIR/PIF_Harbinger | range | 2.1 | 2 | 139.25 | 0.1265 | 0.1362 | ns | ns |
| TIR/Tc1_Mariner | range | 4.96 | 2 | 148.26 | 0.0082 | 0.0128 | ** | * |
| Low_complexity | range | 7.08 | 2 | 157.38 | 0.0011 | 0.002 | ** | ** |
| Simple_repeat | range | 8.42 | 2 | 141.4 | 0.0004 | 0.0007 | *** | *** |
| 8s_rRNA | range | 4.28 | 2 | 126.58 | 0.0159 | 0.0204 | * | * |
| 18s_rRNA | range | 19.28 | 2 | 124.78 | 5.04 x $10^{-8}$ | 1.41 x $10^{-7}$ | *** | *** |
| 28s_rRNA | range | 17.72 | 2 | 128.54 | 1.60 x $10^{-7}$ | 3.73 x $10^{-7}$ | *** | *** |

| TEs and rRNAs | contrast | estimate | SE | df | t.ratio | p_value | Sig |
|---|---|---|---|---|---|---|---|
| LTR/Copia | AUS - EUR | 36.9322 | 5.317 | 144.0035 | 6.9461 | 3.59 x $10^{-10}$ | *** |
| | AUS - NAM | 44.5185 | 5.163 | 141.0133 | 8.6226 | 7.24 x $10^{-14}$ | *** |
| | EUR - NAM | 7.5863 | 2.5423 | 126.2924 | 2.984 | 0.0095 | ** |
| Helitron | AUS - EUR | 5.9432 | 3.1559 | 161.8721 | 1.8832 | 0.1468 | ns |
| | AUS - NAM | 1.9478 | 3.076 | 157.3124 | 0.6332 | 0.8021 | ns |
| | EUR - NAM | -3.9955 | 1.5448 | 150.6326 | -2.5865 | 0.0285 | * |

| | | | | | | | |
|---|---|---|---|---|---|---|---|
| LTR/Gypsy | AUS - EUR | 32.0547 | 5.7712 | 161.1381 | 5.5542 | $3.38 \times 10^{-7}$ | *** |
| | AUS - NAM | 40.3526 | 5.6192 | 156.9374 | 7.1812 | $7.82 \times 10^{-11}$ | *** |
| | EUR - NAM | 8.2979 | 2.8121 | 143.1265 | 2.9508 | 0.0103 | * |
| LTR/unknown | AUS - EUR | 18.847 | 3.0497 | 161.8466 | 6.1799 | $1.50 \times 10^{-8}$ | *** |
| | AUS - NAM | 16.8926 | 2.9727 | 157.2636 | 5.6825 | $1.88 \times 10^{-7}$ | *** |
| | EUR - NAM | -1.9544 | 1.4931 | 151.2151 | -1.3089 | 0.3924 | ns |
| TIR/Mutator | AUS - EUR | 12.6649 | 1.9899 | 138.6159 | 6.3646 | $8.00 \times 10^{-9}$ | *** |
| | AUS - NAM | 15.248 | 1.9314 | 135.8364 | 7.8946 | $2.65 \times 10^{-12}$ | *** |
| | EUR - NAM | 2.5832 | 0.9487 | 123.7798 | 2.7228 | 0.0201 | * |
| TIR/hAT | AUS - EUR | 1.3098 | 1.0823 | 124.35 | 1.2101 | 0.4494 | ns |
| | AUS - NAM | 1.2399 | 1.0496 | 122.0617 | 1.1813 | 0.4666 | ns |
| | EUR - NAM | -0.0699 | 0.5136 | 117.9299 | -0.1361 | 0.9898 | ns |
| TIR/CACTA | AUS - EUR | 0.6456 | 0.2426 | 161.7965 | 2.6614 | 0.0232 | * |
| | AUS - NAM | 0.4173 | 0.2365 | 157.1822 | 1.7647 | 0.1849 | ns |
| | EUR - NAM | -0.2283 | 0.1188 | 152.0252 | -1.9215 | 0.1361 | ns |
| TIR/PIF_Harbinger | AUS - EUR | 0.2292 | 0.1699 | 150.5173 | 1.3495 | 0.3702 | ns |
| | AUS - NAM | 0.3164 | 0.165 | 147.2246 | 1.9172 | 0.1374 | ns |
| | EUR - NAM | 0.0872 | 0.0816 | 129.9767 | 1.0686 | 0.5353 | ns |
| TIR/Tc1_Mariner | AUS - EUR | 0.0331 | 0.033 | 159.598 | 1.0034 | 0.5758 | ns |
| | AUS - NAM | -0.0173 | 0.0321 | 155.6329 | -0.54 | 0.8517 | ns |
| | EUR - NAM | -0.0505 | 0.016 | 139.1901 | -3.1487 | 0.0057 | ** |
| Low_complexity | AUS - EUR | -0.0134 | 0.0255 | 161.3113 | -0.5247 | 0.8594 | ns |
| | AUS - NAM | -0.056 | 0.0249 | 156.5534 | -2.2502 | 0.0661 | ns |
| | EUR - NAM | -0.0426 | 0.0125 | 156.2153 | -3.4059 | 0.0024 | ** |
| Simple_repeat | AUS - EUR | 0.7387 | 0.1877 | 153.0841 | 3.9354 | 0.0004 | *** |
| | AUS - NAM | 0.5321 | 0.1824 | 149.647 | 2.917 | 0.0113 | * |
| | EUR - NAM | -0.2065 | 0.0904 | 131.7991 | -2.2851 | 0.0614 | ns |
| 8s_rRNA | AUS - EUR | 0.003 | 0.0408 | 133.0891 | 0.0733 | 0.997 | ns |
| | AUS - NAM | 0.0567 | 0.0396 | 130.5065 | 1.4325 | 0.3273 | ns |

| | | | | | | | |
|---|---|---|---|---|---|---|---|
| | EUR - NAM | 0.0537 | 0.0194 | 121.4426 | 2.7676 | 0.0178 | * |
| 18s_rRNA | AUS - EUR | 1.4617 | 0.3324 | 130.4871 | 4.3974 | 6.66 x $10^{-5}$ | *** |
| | AUS - NAM | 1.9145 | 0.3225 | 127.9933 | 5.9371 | 7.66 x $10^{-8}$ | *** |
| | EUR - NAM | 0.4528 | 0.158 | 120.3861 | 2.866 | 0.0135 | * |
| 28s_rRNA | AUS - EUR | 3.2618 | 0.9466 | 135.9223 | 3.4459 | 0.0022 | ** |
| | AUS - NAM | 4.8848 | 0.9186 | 133.2406 | 5.3177 | 1.29 x $10^{-6}$ | *** |
| | EUR - NAM | 1.6231 | 0.4508 | 122.6201 | 3.6007 | 0.0013 | ** |

**Suppl. Table S5: Linear mixed-model effects of rRNA/TE families abundance on genome size with PC1, PC2 as covariates and population as a random effect. Significance codes: *** < 0.001, ** < 0.01, * < 0.05, ns = not significant.**

| TEs and rRNAs | F_value | p_value | Sig. | FDR | FDR_sig. |
|---|---|---|---|---|---|
| LTR/Copia | 861.1793 | 2.25 x $10^{-102}$ | *** | 3.15 x $10^{-101}$ | *** |
| Helitron | 130.3308 | 2.13 x $10^{-24}$ | *** | 7.44 x $10^{-24}$ | *** |
| LTR/Gypsy | 252.3147 | 1.78 x $10^{-44}$ | *** | 1.25 x $10^{-43}$ | *** |
| LTR/unknown | 0.91862 | 0.339 | ns | 0.365 | ns |
| TIR/Mutator | 124.3392 | 4.57 x $10^{-25}$ | *** | 2.13 x $10^{-24}$ | *** |
| TIR/hAT | 64.3942 | 1.07 x $10^{-14}$ | *** | 2.51 x $10^{-14}$ | *** |
| TIR/CACTA | 99.3991 | 3.87 x $10^{-21}$ | *** | 1.08 x $10^{-20}$ | *** |
| TIR/PIF_Harbinger | 17.5135 | 3.52 x $10^{-5}$ | *** | 5.47 x $10^{-5}$ | *** |
| TIR/Tc1_Mariner | 7.4279 | 0.0068 | ** | 0.0087 | ** |
| Low_complexity | 22.2882 | 3.22 x $10^{-6}$ | *** | 6.43 x $10^{-6}$ | *** |
| Simple_repeat | 0.4649 | 0.496 | ns | 0.496 | ns |
| 8s_rRNA | 15.964 | 7.62 x $10^{-5}$ | *** | 1.07 x $10^{-4}$ | *** |
| 18s_rRNA | 1.81564 | 0.179 | ns | 0.208 | ns |
| 28s_rRNA | 21.4757 | 4.84 x $10^{-6}$ | *** | 8.46 x $10^{-6}$ | *** |

**Suppl. Table S6: Linear mixed model (base and interaction) for genome size with MAT, PC1, PC2 as fixed effect and population as a random intercept. Significance codes: *** < 0.001, ** < 0.01, * < 0.05, ns = not significant.**

| | F | Df | Df.res | Pr(>F) | Sig. |
|---|---|---|---|---|---|
| range | 18.7924 | 2 | 129.73 | 6.81 x $10^{-8}$ | *** |
| MAT | 8.2482 | 1 | 144.86 | 0.0047 | ** |
| PC1 | 0.5362 | 1 | 208.88 | 0.4648 | ns |
| PC2 | 2.3229 | 1 | 214.9 | 0.1289 | ns |

| | F | Df | Df.res | Pr(>F) | Sig. |
|---|---|---|---|---|---|
| range | 0.7576 | 2 | 83.229 | 0.47199 | ns |
| MAT | 0.7225 | 1 | 68.005 | 0.39831 | ns |
| PC1 | 0.1265 | 1 | 254.046 | 0.72234 | ns |
| PC2 | 3.0169 | 1 | 247.501 | 0.08364 | ns |
| **range*MAT** | **0.3827** | **2** | **83.089** | **0.68325** | **ns** |

**Suppl. Table S7: Linear quantile mixed model (LQMM) results for genome size across all ranges, with range, flowering time polygenic score ($PGV_{FT}$), PC1 and PC2 included as fixed effects and population as a random effect.**

| Tau | Coefficient | SE | P-value | Significant? |
|---|---|---|---|---|
| 0.1 | -1.585 | 1.841 | 0.393 | No |
| 0.25 | 1.353 | 1.769 | 0.448 | No |
| 0.5 | 2.307 | 1.142 | 0.049 | Yes* |
| 0.75 | 2.376 | 1.645 | 0.155 | No |
| 0.9 | 3.499 | 1.881 | 0.069 | No |

**Suppl. Table S8: Linear quantile mixed model (LQMM) results for genome size within each range, with flowering time polygenic score ($PGV_{FT}$), PC1 and PC2 included as fixed effects and population as a random effect.**

| Range | Tau | Coefficient | SE | P-value | Significant? |
|---|---|---|---|---|---|
| North America | 0.1 | -1.031 | 6.708 | 0.785 | No |
| North America | 0.25 | 0.9638 | 6.208 | 0.877 | No |
| North America | 0.5 | 0.9212 | 6.009 | 0.878 | No |
| North America | 0.75 | 2.6765 | 5.751 | 0.643 | No |
| North America | 0.9 | -2.1231 | 6.113 | 0.729 | No |
| Europe | 0.1 | 2.59 | 2.714 | 0.345 | No |
| Europe | 0.25 | 1.405 | 1.573 | 0.376 | No |

| | | | | | |
|---|---|---|---|---|---|
| Europe | 0.5 | 1.77 | 1.37 | 0.203 | No |
| Europe | 0.75 | -0.358 | 1.431 | 0.803 | No |
| Europe | 0.9 | 2.311 | 1.796 | 0.204 | No |
| Australia | 0.1 | 1.274 | 2.265 | 0.576 | No |
| Australia | 0.25 | -0.012 | 1.927 | 0.995 | No |
| Australia | 0.5 | 0.684 | 1.916 | 0.723 | No |
| Australia | 0.75 | 2.153 | 1.757 | 0.226 | No |
| **Australia** | **0.9** | **3.613** | **1.59** | **0.027** | **Yes*** |

**Suppl. Table S9. Sampling information: sequencing coverage, data sources, and sequence-based haploid genome size estimates (Gbp) for 443 *Ambrosia artemisiifolia* individuals across native North America (NAM), invasive Europe (EUR), and invasive Australia (AUS).** Samples highlighted in green indicate those filtered samples, based on the genome size standard deviation (SD) threshold.

| Sample ID | Population | Cluster | Range | Latitude | Longitude | Seq. coverage | Genome size (Gbp) | Year | Reference | Accession |
|---|---|---|---|---|---|---|---|---|---|---|
| 280808-1-10 | 280808-1 | EUR.ME | EUR | 44.7249 | 22.4206 | 2.8000 | 0.9202 | 2013 | Bieker et al. 2022 | ERR7445133 |
| 280808-1-11 | 280808-1 | EUR.ME | EUR | 44.7249 | 22.4206 | 3.5736 | 0.9348 | 2013 | Bieker et al. 2022 | ERR7445144 |
| 280808-1-12 | 280808-1 | EUR.ME | EUR | 44.7249 | 22.4206 | 2.6493 | 0.9272 | 2013 | Bieker et al. 2022 | ERR7445115; ERR7445036 |
| 280808-1-16 | 280808-1 | EUR.ME | EUR | 44.7249 | 22.4206 | 11.4939 | 0.9678 | 2013 | Bieker et al. 2022 | ERR7445482 |
| 280808-1-17 | 280808-1 | EUR.ME | EUR | 44.7249 | 22.4206 | 10.5544 | 0.9912 | 2013 | Bieker et al. 2022 | ERR7445380 |
| 280808-1-24 | 280808-1 | EUR.ME | EUR | 44.7249 | 22.4206 | 2.5170 | 0.9041 | 2013 | Bieker et al. 2022 | ERR7445129 |
| 280808-1-3 | 280808-1 | EUR.ME | EUR | 44.7249 | 22.4206 | 3.2393 | 0.9360 | 2013 | Bieker et al. 2022 | ERR7445140 |
| 280808-1-5 | 280808-1 | EUR.ME | EUR | 44.7249 | 22.4206 | 2.5645 | 0.9646 | 2013 | Bieker et al. 2022 | ERR7445117 |
| 280808-1-8 | 280808-1 | EUR.ME | EUR | 44.7249 | 22.4206 | 2.8111 | 0.9324 | 2013 | Bieker et al. 2022 | ERR7445134 |
| A-2010-T1-1 | A-2010-T1 | EUR | EUR | 47.29309 | 11.0475 | 6.3297 | 0.9310 | 2010 | Bieker et al. 2022 | ERR7445199 |
| A-2010-T1-10 | A-2010-T1 | EUR | EUR | 47.29309 | 11.0475 | 6.7935 | 0.9815 | 2010 | Bieker et al. 2022 | ERR7445272 |

| | | | | | | | | | | |
|---|---|---|---|---|---|---|---|---|---|---|
| A-2010-T1-2 | A-2010-T1 | EUR | EUR | 47.29309 | 11.0475 | 14.1224 | 1.0228 | 2010 | Bieker et al. 2022 | ERR7445574 |
| A-2010-T1-3 | A-2010-T1 | EUR | EUR | 47.29309 | 11.0475 | 8.6968 | 0.9512 | 2010 | Bieker et al. 2022 | ERR7445330 |
| A-2010-T1-4 | A-2010-T1 | EUR | EUR | 47.29309 | 11.0475 | 16.1132 | 0.9374 | 2010 | Bieker et al. 2022 | ERR7445580 |
| A-2010-T1-5 | A-2010-T1 | EUR | EUR | 47.29309 | 11.0475 | 18.3702 | 0.9532 | 2010 | Bieker et al. 2022 | ERR7445601 |
| A-2010-T1-6 | A-2010-T1 | EUR | EUR | 47.29309 | 11.0475 | 9.9380 | 1.0018 | 2010 | Bieker et al. 2022 | ERR7445349 |
| A-2010-T1-7 | A-2010-T1 | EUR | EUR | 47.29309 | 11.0475 | 3.5510 | 0.9912 | 2010 | Bieker et al. 2022 | ERR7445121 |
| A-2010-T1-8 | A-2010-T1 | EUR | EUR | 47.29309 | 11.0475 | 9.0957 | 0.9592 | 2010 | Bieker et al. 2022 | ERR7445311 |
| A-2010-T1-9 | A-2010-T1 | EUR | EUR | 47.29309 | 11.0475 | 7.7156 | 0.9203 | 2010 | Bieker et al. 2022 | ERR7445274 |
| AA16-19 | AA16 | NAM.W | NAM | 44.09456 | -102.87 | 12.9417 | 1.0198 | 2013 | Bieker et al. 2022 | ERR7445552 |
| AA16-2 | AA16 | NAM.W | NAM | 44.09456 | -102.87 | 5.0122 | 0.9357 | 2013 | Bieker et al. 2022 | ERR7445215 |
| AA2-1 | AA2 | NAM.W | NAM | 49.83778 | -97.329 | 3.4365 | 0.9262 | 2013 | Bieker et al. 2022 | ERR7445189 |
| AA2-4 | AA2 | NAM.W | NAM | 49.83778 | -97.329 | 2.4861 | 0.9173 | 2013 | Bieker et al. 2022 | ERR7445154 |
| AA20-19 | AA20 | NAM.W | NAM | 48.14689 | -103.57 | 9.1155 | 1.4384 | 2013 | Bieker et al. 2022 | ERR7445475 |
| AA20-26 | AA20 | NAM.W | NAM | 48.14689 | -103.57 | 9.0128 | 0.9372 | 2013 | Bieker et al. 2022 | ERR7445430 |
| AA20-3 | AA20 | NAM.W | NAM | 48.14689 | -103.57 | 5.1965 | 0.8891 | 2013 | Bieker et al. 2022 | ERR7445214 |
| AA3-15 | AA3 | NAM.W | NAM | 48.19453 | -97.329 | 4.6436 | 0.9493 | 2013 | Bieker et al. 2022 | ERR7445217 |
| AA3-17 | AA3 | NAM.W | NAM | 48.19453 | -97.329 | 5.7912 | 0.9512 | 2013 | Bieker et al. 2022 | ERR7445249 |
| AA3-23 | AA3 | NAM.W | NAM | 48.19453 | -97.329 | 9.5638 | 0.9869 | 2013 | Bieker et al. 2022 | ERR7445439 |
| AA5-1 | AA5 | NAM.W | NAM | 46.21708 | -96.05 | 7.8922 | 0.9392 | 2013 | Bieker et al. 2022 | ERR7445491 |
| AA5-16 | AA5 | NAM.W | NAM | 46.21708 | -96.05 | 5.1967 | 0.9747 | 2013 | Bieker et al. 2022 | ERR7445288 |

| | | | | | | | | | | |
|---|---|---|---|---|---|---|---|---|---|---|
| AA5-23 | AA5 | NAM.W | NAM | 46.21708 | -96.05 | 7.3603 | 0.9844 | 2013 | Bieker et al. 2022 | ERR7445467 |
| AA5-9A | AA5 | NAM.W | NAM | 46.21708 | -96.05 | 4.2347 | 0.9636 | 2013 | Bieker et al. 2022 | ERR7445306 |
| AA7-12 | AA7 | NAM.W | NAM | 44.738 | -95.412 | 5.5950 | 1.0516 | 2013 | Bieker et al. 2022 | ERR7445368 |
| AA7-24 | AA7 | NAM.W | NAM | 44.738 | -95.412 | 4.3831 | 0.9749 | 2013 | Bieker et al. 2022 | ERR7445353 |
| AA7-27 | AA7 | NAM.W | NAM | 44.738 | -95.412 | 5.8029 | 0.9556 | 2013 | Bieker et al. 2022 | ERR7445443 |
| AR-2019-1 | AR-2019 | NAM.ME | NAM | 33.84188 | -93.776 | 2.8629 | 0.9307 | 2009/2010 | Bieker et al. 2022 | ERR7445383 |
| AR-2019-2 | AR-2019 | NAM.ME | NAM | 33.84188 | -93.776 | 2.5417 | 0.8923 | 2009/2010 | Bieker et al. 2022 | ERR7445379 |
| AR-6 | AR | NAM.ME | NAM | 33.97552 | -91.413 | 2.7851 | 0.9683 | 2013 | Bieker et al. 2022 | ERR7445351 |
| AR-7 | AR | NAM.ME | NAM | 33.97552 | -91.413 | 2.7351 | 0.9588 | 2013 | Bieker et al. 2022 | ERR7445361 |
| AU01-1A | AU01 | AUS | AUS | -35.6411 | 150.127 | 7.8092 | 1.1185 | 2014 | Battlay et al. 2025 | SRR30012807 |
| AU01-2A | AU01 | AUS | AUS | -35.6411 | 150.127 | 9.4214 | 1.1001 | 2014 | Battlay et al. 2025 | SRR30012806 |
| AU01-3A | AU01 | AUS | AUS | -35.6411 | 150.127 | 6.0965 | 1.0184 | 2014 | Battlay et al. 2025 | SRR30012795 |
| AU01-6A | AU01 | AUS | AUS | -35.6411 | 150.127 | 2.2997 | 1.2293 | 2014 | Battlay et al. 2025 | SRR30012760 |
| AU01-6D | AU01 | AUS | AUS | -35.6411 | 150.127 | 5.0966 | 1.0969 | 2014 | Battlay et al. 2025 | SRR30012749 |
| AU01-8A | AU01 | AUS | AUS | -35.6411 | 150.127 | 4.1771 | 1.2033 | 2014 | Battlay et al. 2025 | SRR30012786 |
| AU01-8C | AU01 | AUS | AUS | -35.6411 | 150.127 | 3.9732 | 1.0704 | 2014 | Battlay et al. 2025 | SRR30012735 |
| AU05-11A | AU05 | AUS | AUS | -28.7668 | 153.397 | 4.6142 | 1.1250 | 2014 | Battlay et al. 2025 | SRR30012780 |
| AU05-14A | AU05 | AUS | AUS | -28.7668 | 153.397 | 3.2988 | 1.1131 | 2014 | Battlay et al. 2025 | SRR30012769 |
| AU05-16A | AU05 | AUS | AUS | -28.7668 | 153.397 | 3.5193 | 1.1025 | 2014 | Battlay et al. 2025 | SRR30012726 |
| AU05-19A | AU05 | AUS | AUS | -28.7668 | 153.397 | 9.3811 | 1.1179 | 2014 | Battlay et al. 2025 | SRR30012805 |

| | | | | | | | | | | |
|---|---|---|---|---|---|---|---|---|---|---|
| AU05-19B | AU05 | AUS | AUS | -28.7668 | 153.397 | 2.6870 | 1.1489 | 2014 | Battlay et al. 2025 | SRR30012804 |
| AU05-1A | AU05 | AUS | AUS | -28.7668 | 153.397 | 3.4504 | 1.0775 | 2014 | Battlay et al. 2025 | SRR30012803 |
| AU05-20A | AU05 | AUS | AUS | -28.7668 | 153.397 | 3.1082 | 1.0902 | 2014 | Battlay et al. 2025 | SRR30012802 |
| AU05-3B | AU05 | AUS | AUS | -28.7668 | 153.397 | 6.9851 | 1.1500 | 2014 | Battlay et al. 2025 | SRR30012801 |
| AU05-4A | AU05 | AUS | AUS | -28.7668 | 153.397 | 7.2379 | 1.1005 | 2014 | Battlay et al. 2025 | SRR30012800 |
| AU05-5A | AU05 | AUS | AUS | -28.7668 | 153.397 | 4.4765 | 1.0787 | 2014 | Battlay et al. 2025 | SRR30012799 |
| AU09-1A | AU09 | AUS | AUS | -28.8688 | 151.167 | 6.6782 | 1.0581 | 2014 | Battlay et al. 2025 | SRR30012798 |
| AU09-21A | AU09 | AUS | AUS | -28.8688 | 151.167 | 3.1120 | 1.1400 | 2014 | Battlay et al. 2025 | SRR30012797 |
| AU09-22B | AU09 | AUS | AUS | -28.8688 | 151.167 | 9.7181 | 1.0872 | 2014 | Battlay et al. 2025 | SRR30012796 |
| AU09-29A | AU09 | AUS | AUS | -28.8688 | 151.167 | 4.6370 | 1.1047 | 2014 | Battlay et al. 2025 | SRR30012794 |
| AU11-11A | AU11 | AUS | AUS | -25.3655 | 152.916 | 8.2515 | 1.1140 | 2014 | Battlay et al. 2025 | SRR30012793 |
| AU11-15D | AU11 | AUS | AUS | -25.3655 | 152.916 | 6.0123 | 1.0739 | 2014 | Battlay et al. 2025 | SRR30012792 |
| AU11-1C | AU11 | AUS | AUS | -25.3655 | 152.916 | 3.8444 | 1.1096 | 2014 | Battlay et al. 2025 | SRR30012767 |
| AU11-24B | AU11 | AUS | AUS | -25.3655 | 152.916 | 10.5748 | 1.1254 | 2014 | Battlay et al. 2025 | SRR30012766 |
| AU11-5A | AU11 | AUS | AUS | -25.3655 | 152.916 | 5.4231 | 1.1297 | 2014 | Battlay et al. 2025 | SRR30012765 |
| AU11-5B | AU11 | AUS | AUS | -25.3655 | 152.916 | 4.0555 | 1.1283 | 2014 | Battlay et al. 2025 | SRR30012764 |
| AU15-10A | AU15 | AUS | AUS | -27.3792 | 152.802 | 4.8975 | 1.0281 | 2014 | Battlay et al. 2025 | SRR30012763 |
| AU15-14B | AU15 | AUS | AUS | -27.3792 | 152.802 | 2.6132 | 1.0483 | 2014 | Battlay et al. 2025 | SRR30012762 |
| AU15-16A | AU15 | AUS | AUS | -27.3792 | 152.802 | 2.1405 | 1.0912 | 2014 | Battlay et al. 2025 | SRR30012761 |
| AU15-17C | AU15 | AUS | AUS | -27.3792 | 152.802 | 5.6532 | 1.1388 | 2014 | Battlay et al. 2025 | SRR30012759 |

| | | | | | | | | | | |
|---|---|---|---|---|---|---|---|---|---|---|
| AU15-21C | AU15 | AUS | AUS | -27.3792 | 152.802 | 4.7134 | 1.1427 | 2014 | Battlay et al. 2025 | SRR30012758 |
| AU15-23C | AU15 | AUS | AUS | -27.3792 | 152.802 | 3.0344 | 1.1016 | 2014 | Battlay et al. 2025 | SRR30012757 |
| AU15-26A | AU15 | AUS | AUS | -27.3792 | 152.802 | 6.0016 | 1.1336 | 2014 | Battlay et al. 2025 | SRR30012756 |
| AU15-2A | AU15 | AUS | AUS | -27.3792 | 152.802 | 5.2124 | 1.1294 | 2014 | Battlay et al. 2025 | SRR30012755 |
| AU15-9A | AU15 | AUS | AUS | -27.3792 | 152.802 | 6.1289 | 1.0826 | 2014 | Battlay et al. 2025 | SRR30012754 |
| AU18-13A | AU18 | AUS | AUS | -27.785 | 153.275 | 1.8823 | 1.1417 | 2014 | Battlay et al. 2025 | SRR30012753 |
| AU18-14B | AU18 | AUS | AUS | -27.785 | 153.275 | 2.4700 | 1.1693 | 2014 | Battlay et al. 2025 | SRR30012752 |
| AU18-19A | AU18 | AUS | AUS | -27.785 | 153.275 | 17.0203 | 1.0703 | 2014 | Battlay et al. 2025 | SRR30012751 |
| AU18-21A | AU18 | AUS | AUS | -27.785 | 153.275 | 6.9691 | 1.0959 | 2014 | Battlay et al. 2025 | SRR30012750 |
| AU18-25A | AU18 | AUS | AUS | -27.785 | 153.275 | 4.1698 | 1.0681 | 2014 | Battlay et al. 2025 | SRR30012748 |
| AU18-29A | AU18 | AUS | AUS | -27.785 | 153.275 | 1.8191 | 1.1158 | 2014 | Battlay et al. 2025 | SRR30012747 |
| AU18-3A | AU18 | AUS | AUS | -27.785 | 153.275 | 3.9962 | 1.1191 | 2014 | Battlay et al. 2025 | SRR30012746 |
| AU18-4A | AU18 | AUS | AUS | -27.785 | 153.275 | 4.5392 | 1.0843 | 2014 | Battlay et al. 2025 | SRR30012745 |
| AU18-5A | AU18 | AUS | AUS | -27.785 | 153.275 | 2.8429 | 1.1358 | 2014 | Battlay et al. 2025 | SRR30012744 |
| AU18-8B | AU18 | AUS | AUS | -27.785 | 153.275 | 2.4166 | 1.1102 | 2014 | Battlay et al. 2025 | SRR30012791 |
| AU19-10A | AU19 | AUS | AUS | -28.0122 | 153.168 | 2.9225 | 1.1625 | 2014 | Battlay et al. 2025 | SRR30012790 |
| AU19-11A | AU19 | AUS | AUS | -28.0122 | 153.168 | 4.5244 | 1.0975 | 2014 | Battlay et al. 2025 | SRR30012789 |
| AU19-14C | AU19 | AUS | AUS | -28.0122 | 153.168 | 17.9820 | 1.0491 | 2014 | Battlay et al. 2025 | SRR30012788 |
| AU19-21A | AU19 | AUS | AUS | -28.0122 | 153.168 | 12.0686 | 1.1083 | 2014 | Battlay et al. 2025 | SRR30012787 |
| AU19-22C | AU19 | AUS | AUS | -28.0122 | 153.168 | 8.0477 | 1.1244 | 2014 | Battlay et al. 2025 | SRR30012785 |

| | | | | | | | | | | |
|---|---|---|---|---|---|---|---|---|---|---|
| AU19-23A | AU19 | AUS | AUS | -28.0122 | 153.168 | 3.2424 | 1.1458 | 2014 | Battlay et al. 2025 | SRR30012784 |
| AU19-25A | AU19 | AUS | AUS | -28.0122 | 153.168 | 2.0696 | 1.0270 | 2014 | Battlay et al. 2025 | SRR30012743 |
| AU19-29C | AU19 | AUS | AUS | -28.0122 | 153.168 | 9.2488 | 1.0862 | 2014 | Battlay et al. 2025 | SRR30012742 |
| AU19-30A | AU19 | AUS | AUS | -28.0122 | 153.168 | 7.4467 | 1.0752 | 2014 | Battlay et al. 2025 | SRR30012741 |
| AU19-3A | AU19 | AUS | AUS | -28.0122 | 153.168 | 10.1767 | 1.0898 | 2014 | Battlay et al. 2025 | SRR30012740 |
| AU23-10A | AU23 | AUS | AUS | -28.9263 | 152.374 | 4.1243 | 1.0187 | 2014 | Battlay et al. 2025 | SRR30012739 |
| AU23-14A | AU23 | AUS | AUS | -28.9263 | 152.374 | 6.9166 | 1.0623 | 2014 | Battlay et al. 2025 | SRR30012738 |
| AU23-21A | AU23 | AUS | AUS | -28.9263 | 152.374 | 7.2025 | 1.0930 | 2014 | Battlay et al. 2025 | SRR30012737 |
| AU23-24A | AU23 | AUS | AUS | -28.9263 | 152.374 | 7.3190 | 1.1343 | 2014 | Battlay et al. 2025 | SRR30012736 |
| AU23-28A | AU23 | AUS | AUS | -28.9263 | 152.374 | 2.6303 | 1.1685 | 2014 | Battlay et al. 2025 | SRR30012734 |
| AU23-30A | AU23 | AUS | AUS | -28.9263 | 152.374 | 6.8927 | 1.0362 | 2014 | Battlay et al. 2025 | SRR30012733 |
| AU23-5A | AU23 | AUS | AUS | -28.9263 | 152.374 | 1.3554 | 1.1104 | 2014 | Battlay et al. 2025 | SRR30012732 |
| AU23-6A | AU23 | AUS | AUS | -28.9263 | 152.374 | 10.8769 | 1.1280 | 2014 | Battlay et al. 2025 | SRR30012731 |
| AU23-8A | AU23 | AUS | AUS | -28.9263 | 152.374 | 4.9003 | 1.0982 | 2014 | Battlay et al. 2025 | SRR30012730 |
| AU26-11A | AU26 | AUS | AUS | -30.0514 | 152.985 | 4.5953 | 1.0793 | 2014 | Battlay et al. 2025 | SRR30012729 |
| AU26-13A | AU26 | AUS | AUS | -30.0514 | 152.985 | 2.9149 | 1.0423 | 2014 | Battlay et al. 2025 | SRR30012728 |
| AU26-16B | AU26 | AUS | AUS | -30.0514 | 152.985 | 1.6488 | 1.0593 | 2014 | Battlay et al. 2025 | SRR30012783 |
| AU26-1A | AU26 | AUS | AUS | -30.0514 | 152.985 | 7.2188 | 1.0784 | 2014 | Battlay et al. 2025 | SRR30012782 |
| AU26-20A | AU26 | AUS | AUS | -30.0514 | 152.985 | 1.2543 | 1.0697 | 2014 | Battlay et al. 2025 | SRR30012781 |
| AU26-22A | AU26 | AUS | AUS | -30.0514 | 152.985 | 1.5148 | 1.3631 | 2014 | Battlay et al. 2025 | SRR30012779 |

| | | | | | | | | | | |
|---|---|---|---|---|---|---|---|---|---|---|
| AU26-28A | AU26 | AUS | AUS | -30.0514 | 152.985 | 13.6542 | 1.1343 | 2014 | Battlay et al. 2025 | SRR30012778 |
| AU26-29A | AU26 | AUS | AUS | -30.0514 | 152.985 | 3.7611 | 1.0545 | 2014 | Battlay et al. 2025 | SRR30012777 |
| AU26-30B | AU26 | AUS | AUS | -30.0514 | 152.985 | 5.2885 | 1.0889 | 2014 | Battlay et al. 2025 | SRR30012776 |
| AU26-8A | AU26 | AUS | AUS | -30.0514 | 152.985 | 5.2690 | 1.1354 | 2014 | Battlay et al. 2025 | SRR30012775 |
| AU27-10A | AU27 | AUS | AUS | -30.2413 | 152.585 | 5.7891 | 1.1092 | 2014 | Battlay et al. 2025 | SRR30012774 |
| AU27-16D | AU27 | AUS | AUS | -30.2413 | 152.585 | 1.3188 | 1.0917 | 2014 | Battlay et al. 2025 | SRR30012773 |
| AU27-21A | AU27 | AUS | AUS | -30.2413 | 152.585 | 4.1800 | 1.0696 | 2014 | Battlay et al. 2025 | SRR30012772 |
| AU27-22C | AU27 | AUS | AUS | -30.2413 | 152.585 | 18.9797 | 1.1028 | 2014 | Battlay et al. 2025 | SRR30012771 |
| AU27-27A | AU27 | AUS | AUS | -30.2413 | 152.585 | 10.4961 | 1.0843 | 2014 | Battlay et al. 2025 | SRR30012770 |
| AU27-2A | AU27 | AUS | AUS | -30.2413 | 152.585 | 4.0966 | 1.1429 | 2014 | Battlay et al. 2025 | SRR30012768 |
| AU27-2B | AU27 | AUS | AUS | -30.2413 | 152.585 | 2.4224 | 1.1304 | 2014 | Battlay et al. 2025 | SRR30012815 |
| AU27-30A | AU27 | AUS | AUS | -30.2413 | 152.585 | 5.4431 | 1.0754 | 2014 | Battlay et al. 2025 | SRR30012814 |
| AU27-4A | AU27 | AUS | AUS | -30.2413 | 152.585 | 5.6376 | 1.0856 | 2014 | Battlay et al. 2025 | SRR30012813 |
| AU27-9A | AU27 | AUS | AUS | -30.2413 | 152.585 | 6.2148 | 1.0675 | 2014 | Battlay et al. 2025 | SRR30012812 |
| AU30-12A | AU30 | AUS | AUS | -30.7406 | 152.915 | 3.8024 | 1.1864 | 2014 | Battlay et al. 2025 | SRR30012811 |
| AU30-13A | AU30 | AUS | AUS | -30.7406 | 152.915 | 5.0562 | 1.0601 | 2014 | Battlay et al. 2025 | SRR30012810 |
| AU30-16A | AU30 | AUS | AUS | -30.7406 | 152.915 | 4.7620 | 1.0738 | 2014 | Battlay et al. 2025 | SRR30012809 |
| AU30-17B | AU30 | AUS | AUS | -30.7406 | 152.915 | 3.1219 | 1.0566 | 2014 | Battlay et al. 2025 | SRR30012808 |
| AU30-18A | AU30 | AUS | AUS | -30.7406 | 152.915 | 2.9875 | 1.0782 | 2014 | Battlay et al. 2025 | SRR30012727 |
| AU30-1A | AU30 | AUS | AUS | -30.7406 | 152.915 | 2.4033 | 1.0871 | 2014 | Battlay et al. 2025 | SRR30012725 |

| | | | | | | | | | | |
|---|---|---|---|---|---|---|---|---|---|---|
| AU30-21B | AU30 | AUS | AUS | -30.7406 | 152.915 | 5.2359 | 1.0708 | 2014 | Battlay et al. 2025 | SRR30012724 |
| AU30-24A | AU30 | AUS | AUS | -30.7406 | 152.915 | 11.7052 | 1.1226 | 2014 | Battlay et al. 2025 | SRR30012723 |
| AU30-28A | AU30 | AUS | AUS | -30.7406 | 152.915 | 9.9673 | 1.0905 | 2014 | Battlay et al. 2025 | SRR30012722 |
| AU30-29A | AU30 | AUS | AUS | -30.7406 | 152.915 | 8.0096 | 1.0586 | 2014 | Battlay et al. 2025 | SRR30012721 |
| B-1 | B | EUR.ME | EUR | 44.90308 | -0.5352 | 3.5525 | 0.9731 | 2019 | Bieker et al. 2022 | ERR7445587 |
| B-2 | B | EUR.W | EUR | 44.90308 | -0.5352 | 2.6415 | 0.9516 | 2019 | Bieker et al. 2022 | ERR7445449 |
| B-3 | B | EUR.ME | EUR | 44.90308 | -0.5352 | 3.2873 | 0.9770 | 2019 | Bieker et al. 2022 | ERR7445568 |
| B-4 | B | EUR.W | EUR | 44.90308 | -0.5352 | 4.8650 | 0.9614 | 2019 | Bieker et al. 2022 | ERR7445618 |
| B1-1 | B1 | EUR | EUR | 47.87951 | 18.1561 | 2.5077 | 1.0298 | 2014 | Bieker et al. 2022 | ERR7445405 |
| B2-1 | B1 | EUR | EUR | 47.87951 | 18.1561 | 5.7722 | 0.9184 | 2014 | Bieker et al. 2022 | ERR7445510 |
| B3-1 | B1 | EUR | EUR | 47.87951 | 18.1561 | 3.8043 | 1.0001 | 2014 | Bieker et al. 2022 | ERR7445424 |
| BN-ON-6 | BN-ON | NAM.ME | NAM | 45 | -77.7 | 15.5509 | 0.9773 | 2013 | Bieker et al. 2022 | ERR7445918 |
| Caluire-26 | Caluire | EUR.ME | EUR | 45.99 | 4.84 | 3.3096 | 1.0024 | 2017 | Bieker et al. 2022 | ERR7445586 |
| Caluire-27 | Caluire | EUR.ME | EUR | 45.99 | 4.84 | 3.7465 | 0.9695 | 2017 | Bieker et al. 2022 | ERR7445595 |
| Caluire-31 | Caluire | EUR.ME | EUR | 45.99 | 4.84 | 4.7851 | 1.0189 | 2017 | Bieker et al. 2022 | ERR7445609 |
| Caluire-32 | Caluire | EUR.ME | EUR | 45.99 | 4.84 | 4.5922 | 0.9500 | 2017 | Bieker et al. 2022 | ERR7445641 |
| Caluire-36 | Caluire | EUR.ME | EUR | 45.99 | 4.84 | 5.2947 | 1.0228 | 2017 | Bieker et al. 2022 | ERR7445702 |
| Caluire-37 | Caluire | EUR.ME | EUR | 45.99 | 4.84 | 2.2943 | 0.9269 | 2017 | Bieker et al. 2022 | ERR7445612 |
| Caluire-41 | Caluire | EUR.ME | EUR | 45.99 | 4.84 | 4.3034 | 1.0238 | 2017 | Bieker et al. 2022 | ERR7445690 |
| Caluire-42 | Caluire | EUR.ME | EUR | 45.99 | 4.84 | 3.3282 | 0.9984 | 2017 | Bieker et al. 2022 | ERR7445643 |

| | | | | | | | | | | |
|---|---|---|---|---|---|---|---|---|---|---|
| Caluire-46 | Caluire | EUR.ME | EUR | 45.99 | 4.84 | 4.8987 | 1.0116 | 2017 | Bieker et al. 2022 | ERR7445685 |
| Caluire-47 | Caluire | EUR.ME | EUR | 45.99 | 4.84 | 3.1805 | 1.0626 | 2017 | Bieker et al. 2022 | ERR7445649 |
| CT-H-1 | CT-H | NAM.E | NAM | 41.2877 | -73.766 | 12.4788 | 0.9406 | 2009 | Bieker et al. 2022 | ERR7445851 |
| CT-H-10 | CT-H | NAM.E | NAM | 41.2877 | -73.766 | 3.9864 | 0.9875 | 2009 | Bieker et al. 2022 | ERR7445507 |
| CT-H-2 | CT-H | NAM.E | NAM | 41.2877 | -73.766 | 4.8670 | 1.0041 | 2009 | Bieker et al. 2022 | ERR7445527 |
| CT-H-3 | CT-H | NAM.E | NAM | 41.2877 | -73.766 | 6.0880 | 0.9690 | 2009 | Bieker et al. 2022 | ERR7445571 |
| CT-H-4 | CT-H | NAM.E | NAM | 41.2877 | -73.766 | 8.5146 | 0.9764 | 2009 | Bieker et al. 2022 | ERR7445599 |
| CT-H-6 | CT-H | NAM.E | NAM | 41.2877 | -73.766 | 4.4905 | 0.9173 | 2009 | Bieker et al. 2022 | ERR7445588 |
| CT-H-7 | CT-H | NAM.E | NAM | 41.2877 | -73.766 | 2.7977 | 0.9687 | 2009 | Bieker et al. 2022 | ERR7445539 |
| CT-H-8 | CT-H | NAM.E | NAM | 41.2877 | -73.766 | 3.9002 | 0.9979 | 2009 | Bieker et al. 2022 | ERR7445563 |
| CT-H-9 | CT-H | NAM.E | NAM | 41.2877 | -73.766 | 5.3828 | 0.9542 | 2009 | Bieker et al. 2022 | ERR7445598 |
| D-2017-4-10 | D-2017-4 | EUR.E | EUR | 52.8243 | 13.7691 | 12.1407 | 0.9759 | 2017 | Bieker et al. 2022 | ERR7446018 |
| D-2017-4-2 | D-2017-4 | EUR.E | EUR | 52.8243 | 13.7691 | 12.3416 | 0.9948 | 2017 | Bieker et al. 2022 | ERR7446048 |
| D-2017-4-3 | D-2017-4 | EUR.E | EUR | 52.8243 | 13.7691 | 12.5795 | 0.9564 | 2017 | Bieker et al. 2022 | ERR7446052 |
| D-2017-4-4 | D-2017-4 | EUR.E | EUR | 52.8243 | 13.7691 | 3.7315 | 0.9415 | 2017 | Bieker et al. 2022 | ERR7445696 |
| D-2017-4-5 | D-2017-4 | EUR.E | EUR | 52.8243 | 13.7691 | 7.5742 | 0.9965 | 2017 | Bieker et al. 2022 | ERR7445883 |
| D-2017-4-6 | D-2017-4 | EUR.E | EUR | 52.8243 | 13.7691 | 6.7693 | 0.9563 | 2017 | Bieker et al. 2022 | ERR7445759 |
| D-2017-4-7 | D-2017-4 | EUR.E | EUR | 52.8243 | 13.7691 | 8.3414 | 0.9770 | 2017 | Bieker et al. 2022 | ERR7445900 |
| D-2017-4-9 | D-2017-4 | EUR.E | EUR | 52.8243 | 13.7691 | 9.3081 | 0.9946 | 2017 | Bieker et al. 2022 | ERR7445944 |
| EU2-01-12 | EU2-01 | EUR.ME | EUR | 46.03608 | 15.2961 | 1.0341 | 0.8956 | 2014 | Bieker et al. 2022 | ERR7445630 |

| | | | | | | | | | | |
|---|---|---|---|---|---|---|---|---|---|---|
| EU2-02-1 | EU2-02 | EUR | EUR | 47.13087 | 16.9031 | 4.8009 | 1.0026 | 2014 | Bieker et al. 2022 | ERR7445722 |
| EU2-02-10 | EU2-02 | EUR | EUR | 47.13087 | 16.9031 | 9.7312 | 1.1189 | 2014 | Bieker et al. 2022 | ERR7451270; ERR7445700 |
| EU2-03-9C | EU2-03 | EUR.ME | EUR | 47.3278 | 19.7307 | 12.3793 | 0.9902 | 2014 | Bieker et al. 2022 | ERR7446032 |
| EU2-04-11 | EU2-04 | EUR | EUR | 47.87992 | 18.1545 | 3.1462 | 0.9829 | 2014 | Bieker et al. 2022 | ERR7445676 |
| EU2-04-12 | EU2-04 | EUR | EUR | 47.87992 | 18.1545 | 5.2942 | 0.9978 | 2014 | Bieker et al. 2022 | ERR7445760 |
| EU2-05-18 | EU2-05 | EUR.ME | EUR | 45.46996 | 12.1092 | 2.3676 | 0.9650 | 2014 | Bieker et al. 2022 | ERR7445672 |
| EU2-05-23 | EU2-05 | EUR.ME | EUR | 45.46996 | 12.1092 | 6.6578 | 0.9990 | 2014 | Bieker et al. 2022 | ERR7445823 |
| EU2-05-25 | EU2-05 | EUR.ME | EUR | 45.46996 | 12.1092 | 3.9722 | 1.5958 | 2014 | Bieker et al. 2022 | ERR7448827; ERR7445768 |
| EU2-06-15D | EU2-06 | EUR.ME | EUR | 45.57073 | 8.78546 | 14.8513 | 0.9102 | 2014 | Bieker et al. 2022 | ERR7446171 |
| EU2-06-18 | EU2-06 | EUR.ME | EUR | 45.57073 | 8.78546 | 11.8556 | 0.9900 | 2014 | Bieker et al. 2022 | ERR7445725 |
| EU2-08-13 | EU2-08 | EUR.ME | EUR | 45.93087 | 8.98377 | 13.3247 | 0.9725 | 2014 | Bieker et al. 2022 | ERR7446122 |
| EU2-08-21 | EU2-08 | EUR.ME | EUR | 45.93087 | 8.98377 | 8.6689 | 0.9977 | 2014 | Bieker et al. 2022 | ERR7445996 |
| EU2-08-4 | EU2-08 | EUR.ME | EUR | 45.93087 | 8.98377 | 50.8313 | 1.0283 | 2014 | Bieker et al. 2022 | ERR7446626 |
| Eu2-09-1 | EU2-09 | EUR.W | EUR | 45.06541 | 7.59229 | 8.9294 | 0.9787 | 2014 | Bieker et al. 2022 | ERR7446514 |
| EU2-09-10 | EU2-09 | EUR.W | EUR | 45.06541 | 7.59229 | 5.4996 | 1.0298 | 2014 | Bieker et al. 2022 | ERR7445820 |
| EU2-09-13 | EU2-09 | EUR.W | EUR | 45.06541 | 7.59229 | 0.8566 | 0.9847 | 2014 | Bieker et al. 2022 | ERR7445744 |
| EU2-10-10 | EU2-10 | EUR.ME | EUR | 43.93235 | 4.32049 | 8.9118 | 1.0286 | 2014 | Bieker et al. 2022 | ERR7451305 |
| EU2-10-11 | EU2-10 | EUR.ME | EUR | 43.93235 | 4.32049 | 8.9988 | 1.0819 | 2014 | Bieker et al. 2022 | ERR7451254; ERR7445819 |
| EU2-10-12 | EU2-10 | EUR.ME | EUR | 43.93235 | 4.32049 | 13.7131 | 0.9278 | 2014 | Bieker et al. 2022 | ERR7451241; ERR7445891 |
| EU2-10-18 | EU2-10 | EUR.ME | EUR | 43.93235 | 4.32049 | 8.2960 | 1.0439 | 2014 | Bieker et al. 2022 | ERR7448828; ERR7445989 |

| | | | | | | | | | | |
|---|---|---|---|---|---|---|---|---|---|---|
| EU2-10-2 | EU2-10 | EUR.ME | EUR | 43.93235 | 4.32049 | 10.7402 | 1.0359 | 2014 | Bieker et al. 2022 | ERR7448846; ERR7446012 |
| EU2-10-3 | EU2-10 | EUR.ME | EUR | 43.93235 | 4.32049 | 4.8717 | 1.0325 | 2014 | Bieker et al. 2022 | ERR7448849 |
| EU2-10-4 | EU2-10 | EUR.ME | EUR | 43.93235 | 4.32049 | 15.2390 | 1.0144 | 2014 | Bieker et al. 2022 | ERR7451306; ERR7459478 |
| EU2-10-5 | EU2-10 | EUR.ME | EUR | 43.93235 | 4.32049 | 18.4520 | 1.0550 | 2014 | Bieker et al. 2022 | ERR7451323; ERR7446128 |
| EU2-10-7 | EU2-10 | EUR.ME | EUR | 43.93235 | 4.32049 | 8.3755 | 1.0092 | 2014 | Bieker et al. 2022 | ERR7451219; ERR7445878 |
| EU2-10-9 | EU2-10 | EUR.ME | EUR | 43.93235 | 4.32049 | 15.5886 | 1.0134 | 2014 | Bieker et al. 2022 | ERR7451257; ERR7446103 |
| EU2-12-13 | EU2-12 | EUR.ME | EUR | 46.66434 | 4.3278 | 6.8227 | 0.9650 | 2014 | Bieker et al. 2022 | ERR7445959 |
| EU2-14-12 | EU2-14 | EUR.ME | EUR | 47.4556 | 5.21213 | 11.1351 | 0.9982 | 2014 | Bieker et al. 2022 | ERR7446267 |
| EU2-14-13 | EU2-14 | EUR.ME | EUR | 47.4556 | 5.21213 | 6.3861 | 1.1796 | 2014 | Bieker et al. 2022 | ERR7448877; ERR7445939 |
| EU2-16-1 | EU2-16 | EUR.ME | EUR | 46.1622 | 6.0094 | 8.4323 | 1.0306 | 2014 | Bieker et al. 2022 | ERR7446143 |
| EU2-16-12 | EU2-16 | EUR.ME | EUR | 46.1622 | 6.0094 | 11.4039 | 1.0138 | 2014 | Bieker et al. 2022 | ERR7446240 |
| EU2-16-13 | EU2-16 | EUR.ME | EUR | 46.1622 | 6.0094 | 7.9713 | 0.9523 | 2014 | Bieker et al. 2022 | ERR7446153 |
| EU2-17-1 | EU2-17 | EUR | EUR | 51.12003 | 5.8403 | 13.8939 | 1.0430 | 2014 | Bieker et al. 2022 | ERR7448842; ERR7446254 |
| EU2-17-10 | EU2-17 | EUR | EUR | 51.12003 | 5.8403 | 18.0630 | 1.0650 | 2014 | Bieker et al. 2022 | ERR7448838; ERR7446424 |
| EU2-17-11 | EU2-17 | EUR | EUR | 51.12003 | 5.8403 | 52.7390 | 1.0643 | 2014 | Bieker et al. 2022 | ERR7448883; ERR7448735 |
| EU2-17-2 | EU2-17 | EUR | EUR | 51.12003 | 5.8403 | 26.0648 | 1.0454 | 2014 | Bieker et al. 2022 | ERR7448845; ERR7446484 |
| EU2-17-3 | EU2-17 | EUR | EUR | 51.12003 | 5.8403 | 15.7182 | 1.0717 | 2014 | Bieker et al. 2022 | ERR7448841; ERR7446403 |
| EU2-17-5 | EU2-17 | EUR | EUR | 51.12003 | 5.8403 | 11.8416 | 0.9560 | 2014 | Bieker et al. 2022 | ERR7448851; ERR7446234 |
| EU2-17-6 | EU2-17 | EUR | EUR | 51.12003 | 5.8403 | 13.6380 | 0.9268 | 2014 | Bieker et al. 2022 | ERR7446431 |
| EU2-17-7 | EU2-17 | EUR | EUR | 51.12003 | 5.8403 | 13.6515 | 1.0263 | 2014 | Bieker et al. 2022 | ERR7448850; ERR7446300 |

| | | | | | | | | | | |
|---|---|---|---|---|---|---|---|---|---|---|
| EU2-17-8 | EU2-17 | EUR | EUR | 51.12003 | 5.8403 | 7.3054 | 1.1315 | 2014 | Bieker et al. 2022 | ERR7448858; ERR7446151 |
| EU2-17-9 | EU2-17 | EUR | EUR | 51.12003 | 5.8403 | 7.6998 | 1.0897 | 2014 | Bieker et al. 2022 | ERR7448859; ERR7446146 |
| EU2-19-1 | EU2-19 | EUR | EUR | 56.16811 | 15.8904 | 2.1869 | 1.2119 | 2014 | Bieker et al. 2022 | ERR7446125 |
| EU2-19-2 | EU2-19 | EUR.ME | EUR | 56.16811 | 15.8904 | 1.6688 | 1.0159 | 2014 | Bieker et al. 2022 | ERR7446159 |
| EU2-20-1 | EU2-20 | EUR.E | EUR | 51.63281 | 14.1844 | 4.4941 | 1.0168 | 2014 | Bieker et al. 2022 | ERR7446204 |
| EU2-20-10 | EU2-20 | EUR.E | EUR | 51.63281 | 14.1844 | 6.5836 | 1.1608 | 2014 | Bieker et al. 2022 | ERR7448869; ERR7446195 |
| EU2-20-11 | EU2-20 | EUR.E | EUR | 51.63281 | 14.1844 | 7.8623 | 1.0114 | 2014 | Bieker et al. 2022 | ERR7446281 |
| EU2-21-13 | EU2-21 | EUR | EUR | 50.19005 | 15.0604 | 10.6246 | 0.9953 | 2014 | Bieker et al. 2022 | ERR7446433 |
| EU2-21-14 | EU2-21 | EUR | EUR | 50.19005 | 15.0604 | 10.5727 | 0.9792 | 2014 | Bieker et al. 2022 | ERR7446435 |
| EU2-21-16 | EU2-21 | EUR | EUR | 50.19005 | 15.0604 | 34.4461 | 0.9786 | 2014 | Bieker et al. 2022 | ERR7446589 |
| EU2-21-18 | EU2-21 | EUR | EUR | 50.19005 | 15.0604 | 10.0232 | 0.9698 | 2014 | Bieker et al. 2022 | ERR7446430 |
| EU2-21-20 | EU2-21 | EUR | EUR | 50.19005 | 15.0604 | 11.1300 | 0.9915 | 2014 | Bieker et al. 2022 | ERR7446439 |
| EU2-21-22 | EU2-21 | EUR | EUR | 50.19005 | 15.0604 | 4.6874 | 0.9640 | 2014 | Bieker et al. 2022 | ERR7446288 |
| EU2-21-24 | EU2-21 | EUR | EUR | 50.19005 | 15.0604 | 9.9167 | 1.0089 | 2014 | Bieker et al. 2022 | ERR7446438 |
| EU2-21-27 | EU2-21 | EUR | EUR | 50.19005 | 15.0604 | 5.8702 | 0.9853 | 2014 | Bieker et al. 2022 | ERR7446349 |
| EU2-21-28 | EU2-21 | EUR | EUR | 50.19005 | 15.0604 | 16.3208 | 1.1128 | 2014 | Bieker et al. 2022 | ERR7451426; ERR7446451 |
| EU2-21-29 | EU2-21 | EUR | EUR | 50.19005 | 15.0604 | 7.0101 | 0.9874 | 2014 | Bieker et al. 2022 | ERR7446411 |
| EU2-22-1 | EU2-22 | EUR | EUR | 49.41803 | 17.9615 | 8.7722 | 0.9522 | 2014 | Bieker et al. 2022 | ERR7446450 |
| EU2-22-10 | EU2-22 | EUR | EUR | 49.41803 | 17.9615 | 24.7317 | 0.9767 | 2014 | Bieker et al. 2022 | ERR7446528 |
| EU2-22-13 | EU2-22 | EUR | EUR | 49.41803 | 17.9615 | 9.4278 | 0.9763 | 2014 | Bieker et al. 2022 | ERR7446454 |

| | | | | | | | | | | |
|---|---|---|---|---|---|---|---|---|---|---|
| EU2-22-15 | EU2-22 | EUR | EUR | 49.41803 | 17.9615 | 4.2722 | 0.9318 | 2014 | Bieker et al. 2022 | ERR7446402 |
| EU2-22-18 | EU2-22 | EUR | EUR | 49.41803 | 17.9615 | 6.4411 | 1.0050 | 2014 | Bieker et al. 2022 | ERR7446440 |
| EU2-22-19 | EU2-22 | EUR | EUR | 49.41803 | 17.9615 | 8.9962 | 0.9476 | 2014 | Bieker et al. 2022 | ERR7446475 |
| EU2-22-22 | EU2-22 | EUR | EUR | 49.41803 | 17.9615 | 4.1665 | 0.9615 | 2014 | Bieker et al. 2022 | ERR7446434 |
| EU2-22-24 | EU2-22 | EUR | EUR | 49.41803 | 17.9615 | 4.6104 | 1.0241 | 2014 | Bieker et al. 2022 | ERR7446437 |
| EU2-22-25 | EU2-22 | EUR | EUR | 49.41803 | 17.9615 | 6.0168 | 0.9129 | 2014 | Bieker et al. 2022 | ERR7446446 |
| EU2-22-26 | EU2-22 | EUR | EUR | 49.41803 | 17.9615 | 7.8802 | 0.9739 | 2014 | Bieker et al. 2022 | ERR7446481 |
| EU2-23-10 | EU2-23 | EUR.W | EUR | 50.44297 | 18.8634 | 4.6617 | 1.0162 | 2014 | Bieker et al. 2022 | ERR7446445 |
| EU2-23-12 | EU2-23 | EUR.W | EUR | 50.44297 | 18.8634 | 0.7851 | 0.9851 | 2014 | Bieker et al. 2022 | ERR7446436 |
| EU2-23-17 | EU2-23 | EUR.W | EUR | 50.44297 | 18.8634 | 7.9307 | 1.0145 | 2014 | Bieker et al. 2022 | ERR7446492 |
| EU2-24-1 | EU2-24 | EUR.ME | EUR | 48.48919 | 21.8062 | 4.4369 | 1.1295 | 2014 | Bieker et al. 2022 | ERR7448892 |
| EU2-24-10 | EU2-24 | EUR.ME | EUR | 48.48919 | 21.8062 | 7.0198 | 0.9585 | 2014 | Bieker et al. 2022 | ERR7446458 |
| EU2-25-14 | EU2-25 | EUR.ME | EUR | 47.9773 | 23.0443 | 4.5275 | 0.9649 | 2014 | Bieker et al. 2022 | ERR7446447 |
| EU2-25-15 | EU2-25 | EUR.W | EUR | 47.9773 | 23.0443 | 2.2453 | 0.9677 | 2014 | Bieker et al. 2022 | ERR7446441 |
| EU2-26-1 | EU2-26 | EUR.ME | EUR | 46.23696 | 24.854 | 3.4625 | 0.9341 | 2014 | Bieker et al. 2022 | ERR7446443 |
| EU2-26-12 | EU2-26 | EUR.ME | EUR | 46.23696 | 24.854 | 3.0609 | 1.1769 | 2014 | Bieker et al. 2022 | ERR7448874; ERR7446442 |
| EU2-30-1 | EU2-30 | EUR.W | EUR | 49.86851 | 23.0118 | 3.3174 | 0.9526 | 2014 | Bieker et al. 2022 | ERR7446444 |
| EU2-30-12 | EU2-30 | EUR.W | EUR | 49.86851 | 23.0118 | 5.6438 | 0.9525 | 2014 | Bieker et al. 2022 | ERR7446457 |
| EU2-31-1 | EU2-31 | EUR.ME | EUR | 45.37402 | 27.072 | 6.2095 | 1.0012 | 2014 | Bieker et al. 2022 | ERR7446483 |
| EU2-31-11 | EU2-31 | EUR.ME | EUR | 45.37402 | 27.072 | 3.3948 | 0.9287 | 2014 | Bieker et al. 2022 | ERR7446449 |

| | | | | | | | | | | |
|---|---|---|---|---|---|---|---|---|---|---|
| EU2-32-12 | EU2-32 | EUR.ME | EUR | 44.16268 | 28.5098 | 5.0553 | 0.9452 | 2014 | Bieker et al. 2022 | ERR7446464 |
| EU2-32-13 | EU2-32 | EUR.ME | EUR | 44.16268 | 28.5098 | 4.8814 | 1.1201 | 2014 | Bieker et al. 2022 | ERR7448875; ERR7446452 |
| EU2-33-1 | EU2-33 | EUR.ME | EUR | 44.40457 | 26.1348 | 4.1748 | 0.9816 | 2014 | Bieker et al. 2022 | ERR7446455 |
| EU2-33-10 | EU2-33 | EUR.ME | EUR | 44.40457 | 26.1348 | 4.9777 | 0.9643 | 2014 | Bieker et al. 2022 | ERR7446468 |
| EU2-35-1 | EU2-35 | EUR.ME | EUR | 43.09184 | 21.938 | 4.2050 | 0.9587 | 2014 | Bieker et al. 2022 | ERR7446471 |
| EU2-35-14 | EU2-35 | EUR.ME | EUR | 43.09184 | 21.938 | 3.5622 | 0.9866 | 2014 | Bieker et al. 2022 | ERR7446469 |
| EU2-36-1 | EU2-36 | EUR.ME | EUR | 43.31519 | 24.2598 | 1.2617 | 1.0470 | 2014 | Bieker et al. 2022 | ERR7446456 |
| EU2-36-11 | EU2-36 | EUR.ME | EUR | 43.31519 | 24.2598 | 3.9894 | 0.9995 | 2014 | Bieker et al. 2022 | ERR7446470 |
| EU2-37-1 | EU2-37 | EUR.W | EUR | 43.91788 | 20.7331 | 3.6060 | 0.9249 | 2014 | Bieker et al. 2022 | ERR7446472 |
| EU2-37-11 | EU2-37 | EUR | EUR | 43.91788 | 20.7331 | 3.0739 | 0.9519 | 2014 | Bieker et al. 2022 | ERR7446473 |
| EU2-38-12D | EU2-38 | EUR.ME | EUR | 45.71531 | 15.6538 | 8.1826 | 0.9676 | 2014 | Bieker et al. 2022 | ERR7446511 |
| EU2-38-13D | EU2-38 | EUR.ME | EUR | 45.71531 | 15.6538 | 12.1958 | 0.9833 | 2014 | Bieker et al. 2022 | ERR7446526 |
| FL-26 | FL | NAM.S | NAM | 30.4063 | -83.14 | 13.3367 | 1.1432 | 2013 | Bieker et al. 2022 | ERR7451287; ERR7446520 |
| FL-9 | FL | NAM.S | NAM | 30.4063 | -83.14 | 11.2490 | 1.0084 | 2013 | Bieker et al. 2022 | ERR7451432; ERR7446477 |
| FL-A-1 | FLA | NAM.S | NAM | 25.54179 | -80.413 | 5.9216 | 1.0107 | 2009 | Bieker et al. 2022 | ERR7446494 |
| FL-B-1 | FLB | NAM.S | NAM | 27.76424 | -81.595 | 0.8935 | 1.0021 | 2009 | Bieker et al. 2022 | ERR7446474 |
| FL-B-2 | FLB | NAM.S | NAM | 27.76424 | -81.595 | 2.8742 | 0.9945 | 2009 | Bieker et al. 2022 | ERR7446478 |
| FL-D-2018-1 | FLD | NAM.S | NAM | 27.37437 | -80.416 | 4.1269 | 0.9944 | 2009 | Bieker et al. 2022 | ERR7446489 |
| FL-D-2018-2 | FLD | NAM.S | NAM | 27.37437 | -80.416 | 4.4383 | 1.0179 | 2009 | Bieker et al. 2022 | ERR7446488 |
| FL-D-2018-3 | FLD | NAM.S | NAM | 27.37437 | -80.416 | 4.0206 | 1.0147 | 2009 | Bieker et al. 2022 | ERR7446490 |

| | | | | | | | | | | |
|---|---|---|---|---|---|---|---|---|---|---|
| FL-E-1 | FLE | NAM.S | NAM | 29.35716 | -81.16 | 0.9599 | 0.9893 | 2009 | Bieker et al. 2022 | ERR7446479 |
| FL-E-2 | FLE | NAM.S | NAM | 29.35716 | -81.16 | 0.4387 | 0.9846 | 2009 | Bieker et al. 2022 | ERR7446476 |
| FL-F-1 | FL-F | NAM.ME | NAM | 30.69589 | -86.751 | 3.7919 | 1.0308 | 2009 | Bieker et al. 2022 | ERR7446486 |
| FL-F-2 | FL-F | NAM.ME | NAM | 30.69589 | -86.751 | 3.4098 | 0.9996 | 2009 | Bieker et al. 2022 | ERR7446491 |
| FLD-1 | FLD | NAM.S | NAM | 27.37437 | -80.416 | 13.5614 | 1.0756 | 2009 | Bieker et al. 2022 | ERR3597488 |
| FLD-4 | FLD | NAM.S | NAM | 27.37437 | -80.416 | 1.2682 | 1.0723 | 2009 | Bieker et al. 2022 | ERR3602084 |
| FLD-5 | FLD | NAM.S | NAM | 27.37437 | -80.416 | 1.2529 | 1.0306 | 2009 | Bieker et al. 2022 | ERR3597473 |
| FLD-6 | FLD | NAM.S | NAM | 27.37437 | -80.416 | 1.4214 | 1.0491 | 2009 | Bieker et al. 2022 | ERR3597475 |
| FLD-7 | FLD | NAM.S | NAM | 27.37437 | -80.416 | 3.6621 | 1.1355 | 2009 | Bieker et al. 2022 | ERR3597480 |
| FLG-7 | FLG | NAM.S | NAM | 27.08522 | -81.795 | 9.9631 | 1.0374 | 2009 | Bieker et al. 2022 | ERR3597486 |
| FLG-8 | FLG | NAM.S | NAM | 27.08522 | -81.795 | 10.3597 | 1.0853 | 2009 | Bieker et al. 2022 | ERR3597487 |
| FLH-2018-1 | FLH-2018 | NAM.S | NAM | 29.95931 | -82.712 | 3.1566 | 1.0225 | 2009 | Bieker et al. 2022 | ERR7446487 |
| FLH-2018-2 | FLH-2018 | NAM.S | NAM | 29.95931 | -82.712 | 2.4126 | 1.0619 | 2009 | Bieker et al. 2022 | ERR7446485 |
| FLH-2018-3 | FLH-2018 | NAM.S | NAM | 29.95931 | -82.712 | 3.6050 | 1.0035 | 2009 | Bieker et al. 2022 | ERR7446493 |
| FLH-2018-5 | FLH-2018 | NAM.S | NAM | 29.95931 | -82.712 | 3.4336 | 1.0452 | 2009 | Bieker et al. 2022 | ERR7446496 |
| FLH-2018-6 | FLH-2018 | NAM.S | NAM | 29.95931 | -82.712 | 6.4303 | 1.0580 | 2009 | Bieker et al. 2022 | ERR7446504 |
| FR-5 | FR | EUR.ME | EUR | 45.67878 | 4.97993 | 4.3333 | 1.1580 | 2008 | Bieker et al. 2022 | ERR7446501 |
| FR1-26 | FR | EUR.ME | EUR | 45.08023 | 4.75744 | 3.9832 | 0.9699 | 2008 | Bieker et al. 2022 | ERR7446500 |
| FR2-1 | FR | EUR.ME | EUR | 45.67878 | 4.97993 | 6.0073 | 1.1671 | 2008 | Bieker et al. 2022 | ERR7446508 |
| FR2-7 | FR | EUR.ME | EUR | 45.67878 | 4.97993 | 2.7561 | 1.2280 | 2008 | Bieker et al. 2022 | ERR7446499 |

| | | | | | | | | | | |
|---|---|---|---|---|---|---|---|---|---|---|
| FR7-3 | FR7 | EUR.ME | EUR | 47.17582 | 3.01463 | 5.8207 | 1.1015 | 2008 | Bieker et al. 2022 | ERR7446507 |
| FR7-6 | FR7 | EUR.ME | EUR | 47.17582 | 3.01463 | 2.2410 | 1.1991 | 2008 | Bieker et al. 2022 | ERR7446497 |
| FR8-19 | FR8 | EUR.ME | EUR | 44.21666 | 4.26401 | 12.0241 | 0.9589 | 2008 | Bieker et al. 2022 | ERR7446540 |
| FR9-5 | FR9 | EUR.ME | EUR | 44.75302 | 4.87052 | 6.5927 | 1.0966 | 2008 | Bieker et al. 2022 | ERR7446513 |
| GA-1 | GA | NAM.ME | NAM | 31.67867 | -83.196 | 0.8414 | 0.9963 | 2009 | Bieker et al. 2022 | ERR7446495 |
| GA-2 | GA | NAM.ME | NAM | 31.67867 | -83.196 | 4.2582 | 0.9978 | 2009 | Bieker et al. 2022 | ERR7446502 |
| IA-1 | IA | NAM.W | NAM | 41.87498 | -93.483 | 5.5944 | 0.9539 | 2009 | Bieker et al. 2022 | ERR7446512 |
| IA-10 | IA | NAM.W | NAM | 41.87498 | -93.483 | 3.2975 | 0.9452 | 2009 | Bieker et al. 2022 | ERR7446503 |
| IA-2 | IA | NAM.W | NAM | 41.87498 | -93.483 | 3.7093 | 0.9659 | 2009 | Bieker et al. 2022 | ERR7446506 |
| IA-3 | IA | NAM.W | NAM | 41.87498 | -93.483 | 3.2523 | 0.9545 | 2009 | Bieker et al. 2022 | ERR7446505 |
| IA-4 | IA | NAM.W | NAM | 41.87498 | -93.483 | 5.0982 | 0.9922 | 2009 | Bieker et al. 2022 | ERR7446516 |
| IA-5 | IA | NAM.W | NAM | 41.87498 | -93.483 | 2.9989 | 0.9653 | 2009 | Bieker et al. 2022 | ERR7446509 |
| IA-6 | IA | NAM.W | NAM | 41.87498 | -93.483 | 9.9222 | 0.9963 | 2009 | Bieker et al. 2022 | ERR7446544 |
| IA-7 | IA | NAM.W | NAM | 41.87498 | -93.483 | 4.6000 | 0.9115 | 2009 | Bieker et al. 2022 | ERR7446522 |
| IA-8 | IA | NAM.W | NAM | 41.87498 | -93.483 | 3.6117 | 0.9794 | 2009 | Bieker et al. 2022 | ERR7446517 |
| IA-9 | IA | NAM.W | NAM | 41.87498 | -93.483 | 4.0651 | 0.9519 | 2009 | Bieker et al. 2022 | ERR7446518 |
| IL-1 | IL | NAM.ME | NAM | 39.23775 | -88.151 | 3.5282 | 0.9368 | 2009 | Bieker et al. 2022 | ERR7446515 |
| IL-2 | IL | NAM.ME | NAM | 39.23775 | -88.151 | 13.6389 | 0.9802 | 2009 | Bieker et al. 2022 | ERR7446653 |
| IW-10 | IW | NAM.W | NAM | 42.67797 | -96.502 | 18.7807 | 1.1914 | 2013 | Bieker et al. 2022 | ERR7451325; ERR7446648 |
| IW-11 | IW | NAM.W | NAM | 42.67797 | -96.502 | 4.1507 | 0.9730 | 2013 | Bieker et al. 2022 | ERR7446524 |

| | | | | | | | | | | |
|---|---|---|---|---|---|---|---|---|---|---|
| K12-1 | K | EUR.ME | EUR | 45.25472 | 19.9123 | 7.6658 | 0.9882 | 2017 | Bieker et al. 2022 | ERR7446537 |
| K13-1 | K | EUR | EUR | 45.25472 | 19.9123 | 13.9745 | 0.9809 | 2017 | Bieker et al. 2022 | ERR7446702 |
| K16-1 | K | EUR | EUR | 45.25472 | 19.9123 | 12.9292 | 0.8971 | 2017 | Bieker et al. 2022 | ERR7446633 |
| K2-1 | K | EUR | EUR | 45.25472 | 19.9123 | 8.4464 | 0.9540 | 2017 | Bieker et al. 2022 | ERR7446538 |
| KY-14 | KY | NAM.ME | NAM | 38.62588 | -85.007 | 12.3392 | 1.0120 | 2013 | Bieker et al. 2022 | ERR7446641 |
| KY-29 | KY | NAM.ME | NAM | 38.62588 | -85.007 | 9.8306 | 0.9731 | 2013 | Bieker et al. 2022 | ERR7446590 |
| MA-A-1 | MA-A | NAM.E | NAM | 42.31109 | -71.161 | 4.8235 | 0.9672 | 2009 | Bieker et al. 2022 | ERR7446529 |
| MA-A-10 | MA-A | NAM.E | NAM | 42.31109 | -71.161 | 6.0982 | 0.9245 | 2009 | Bieker et al. 2022 | ERR7446531 |
| MA-A-2 | MA-A | NAM.E | NAM | 42.31109 | -71.161 | 4.0472 | 0.9578 | 2009 | Bieker et al. 2022 | ERR7446527 |
| MA-A-3 | MA-A | NAM.E | NAM | 42.31109 | -71.161 | 7.7527 | 1.0113 | 2009 | Bieker et al. 2022 | ERR7446588 |
| MA-A-4 | MA-A | NAM.E | NAM | 42.31109 | -71.161 | 6.4091 | 0.9485 | 2009 | Bieker et al. 2022 | ERR7446543 |
| MA-A-5 | MA-A | NAM.E | NAM | 42.31109 | -71.161 | 6.2008 | 0.9328 | 2009 | Bieker et al. 2022 | ERR7446546 |
| MA-A-6 | MA-A | NAM.E | NAM | 42.31109 | -71.161 | 5.2797 | 0.9307 | 2009 | Bieker et al. 2022 | ERR7446533 |
| MA-A-7 | MA-A | NAM.E | NAM | 42.31109 | -71.161 | 5.2108 | 1.0136 | 2009 | Bieker et al. 2022 | ERR7446534 |
| MA-A-8 | MA-A | NAM.E | NAM | 42.31109 | -71.161 | 7.6604 | 0.9536 | 2009 | Bieker et al. 2022 | ERR7446623 |
| MA-A-9 | MA-A | NAM.E | NAM | 42.31109 | -71.161 | 5.1472 | 0.9501 | 2009 | Bieker et al. 2022 | ERR7446536 |
| ME-C-1 | ME-C | NAM.E | NAM | 44.01332 | -70.069 | 8.7372 | 0.8921 | 2009 | Bieker et al. 2022 | ERR7446639 |
| ME-C-2 | ME-C | NAM.E | NAM | 44.01332 | -70.069 | 10.4276 | 0.9416 | 2009 | Bieker et al. 2022 | ERR7447157 |
| ME-E-1 | ME-E | NAM.E | NAM | 45.15784 | -67.289 | 2.5485 | 0.9457 | 2009 | Bieker et al. 2022 | ERR7446539 |
| ME-E-2 | ME-E | NAM.E | NAM | 45.15784 | -67.289 | 14.5752 | 0.9583 | 2009 | Bieker et al. 2022 | ERR7448117 |

| | | | | | | | | | | |
|---|---|---|---|---|---|---|---|---|---|---|
| MI-2 | MI | NAM.ME | NAM | 46.35814 | -84.881 | 0.9342 | 1.0652 | 2013 | Bieker et al. 2022 | ERR7446542 |
| MI-3 | MI | NAM.ME | NAM | 46.35814 | -84.881 | 3.6440 | 1.6317 | 2013 | Bieker et al. 2022 | ERR7450296; ERR7446545 |
| MI-B-2019-1 | MI-B-2019 | NAM.ME | NAM | 43.15405 | -84.504 | 3.2586 | 0.9443 | 2009 | Bieker et al. 2022 | ERR7446611 |
| MI-B-2019-2 | MI-B-2019 | NAM.ME | NAM | 43.15405 | -84.504 | 2.9574 | 0.9627 | 2009 | Bieker et al. 2022 | ERR7446610 |
| MN2-1 | MN2 | NAM.ME | NAM | 44.44716 | -79.804 | 7.7749 | 0.9917 | 2013 | Bieker et al. 2022 | ERR7447078 |
| MN2-3 | MN2 | NAM.ME | NAM | 44.44716 | -79.804 | 13.4924 | 0.9777 | 2013 | Bieker et al. 2022 | ERR7448167 |
| MO-A-2019-1 | MO-A-2019 | NAM.W | NAM | 38.96843 | -93.879 | 2.7041 | 0.9194 | 2009 | Bieker et al. 2022 | ERR7446608 |
| MO-A-2019-2 | MO-A-2019 | NAM.W | NAM | 38.96843 | -93.879 | 3.8987 | 0.9219 | 2009 | Bieker et al. 2022 | ERR7446654 |
| MO-B-2018-3 | MO-B-2018 | NAM.ME | NAM | 38.38635 | -90.259 | 6.4414 | 0.9627 | 2009 | Bieker et al. 2022 | ERR7446703 |
| MO-B-2018-4 | MO-B-2018 | NAM.ME | NAM | 38.38635 | -90.259 | 4.5954 | 1.0263 | 2009 | Bieker et al. 2022 | ERR7446665 |
| MO-B-2018-5 | MO-B-2018 | NAM.ME | NAM | 38.38635 | -90.259 | 7.9265 | 1.0148 | 2009 | Bieker et al. 2022 | ERR7447534 |
| MO-B-2018-6 | MO-B-2018 | NAM.ME | NAM | 38.38635 | -90.259 | 4.0190 | 0.9691 | 2009 | Bieker et al. 2022 | ERR7446679 |
| MO-B-2018-7 | MO-B-2018 | NAM.ME | NAM | 38.38635 | -90.259 | 3.3254 | 0.9741 | 2009 | Bieker et al. 2022 | ERR7446666 |
| MO-E-2019-1 | MO-E-2019 | NAM.ME | NAM | 36.51766 | -90.249 | 7.0992 | 0.9525 | 2009 | Bieker et al. 2022 | ERR7448103 |
| MO-E-2019-2 | MO-E-2019 | NAM.ME | NAM | 36.51766 | -90.249 | 3.6192 | 0.9228 | 2009 | Bieker et al. 2022 | ERR7446748 |
| MO-G-1 | MO-G | NAM.W | NAM | 36.66569 | -94.347 | 0.6099 | 1.0188 | 2009 | Bieker et al. 2022 | ERR7446640 |
| MO-G-2 | MO-G | NAM.W | NAM | 36.66569 | -94.347 | 4.1773 | 0.9825 | 2009 | Bieker et al. 2022 | ERR7446722 |
| MOB-4 | MO-B-2018 | NAM.ME | NAM | 38.38635 | -90.259 | 1.6081 | 1.0983 | 2009 | Bieker et al. 2022 | ERR3602086 |
| MOB-7 | MO-B-2018 | NAM.ME | NAM | 38.38635 | -90.259 | 1.9132 | 1.0265 | 2009 | Bieker et al. 2022 | ERR3602089 |
| MOB-8 | MO-B-2018 | NAM.ME | NAM | 38.38635 | -90.259 | 7.2387 | 1.0883 | 2009 | Bieker et al. 2022 | ERR3597485 |

| | | | | | | | | | | |
|---|---|---|---|---|---|---|---|---|---|---|
| MP-16 | MP | NAM.ME | NAM | 31.20781 | -89.066 | 12.6345 | 1.0097 | 2013 | Bieker et al. 2022 | ERR7451248; ERR7447519 |
| MP-18 | MP | NAM.ME | NAM | 31.20781 | -89.066 | 15.1605 | 1.0098 | 2013 | Bieker et al. 2022 | ERR7451347; ERR7448094 |
| MS-1 | MS | NAM.E | NAM | 42.0883 | -72.096 | 12.8406 | 1.0085 | 2013 | Bieker et al. 2022 | ERR7448186 |
| MS-3 | MS | NAM.E | NAM | 42.0883 | -72.096 | 8.2205 | 0.9596 | 2013 | Bieker et al. 2022 | ERR7447941 |
| NB-A-1 | NB-A | NAM.ME | NAM | 46.03306 | -65.041 | 9.3975 | 0.9366 | 2009 | Bieker et al. 2022 | ERR7448100 |
| NB-A-2 | NB-A | NAM.ME | NAM | 46.03306 | -65.041 | 13.1043 | 0.9726 | 2009 | Bieker et al. 2022 | ERR7448220 |
| NC-10-F | NC | NAM.ME | NAM | 35.60699 | -83.022 | 3.5650 | 0.9892 | 2013 | Bieker et al. 2022 | ERR7446716 |
| NC-9D | NC | NAM.ME | NAM | 35.60699 | -83.022 | 9.5657 | 1.0252 | 2013 | Bieker et al. 2022 | ERR7448102 |
| NH-A-1 | NH-A | NAM.E | NAM | 42.89069 | -72.528 | 6.5310 | 1.0184 | 2009 | Bieker et al. 2022 | ERR7447211 |
| NH-E-1 | NH-E | NAM.E | NAM | 43.1863 | -71.14 | 7.7417 | 0.9858 | 2009 | Bieker et al. 2022 | ERR7447499 |
| NH-E-2 | NH-E | NAM.E | NAM | 43.1863 | -71.14 | 6.2738 | 0.9523 | 2009 | Bieker et al. 2022 | ERR7447176 |
| NJ-C-1 | NJ-C | NAM.E | NAM | 39.62068 | -75.506 | 7.2094 | 0.9867 | 2009 | Bieker et al. 2022 | ERR7448053 |
| NJ-C-2 | NJ-C | NAM.E | NAM | 39.62068 | -75.506 | 7.6935 | 0.9548 | 2009 | Bieker et al. 2022 | ERR7447502 |
| NJ-C-3 | NJ-C | NAM.E | NAM | 39.62068 | -75.506 | 5.2191 | 1.0000 | 2009 | Bieker et al. 2022 | ERR7447174 |
| NJ-C-4 | NJ-C | NAM.E | NAM | 39.62068 | -75.506 | 5.3718 | 0.9768 | 2009 | Bieker et al. 2022 | ERR7447760 |
| NJ-C-6 | NJ-C | NAM.E | NAM | 39.62068 | -75.506 | 11.3707 | 0.9923 | 2009 | Bieker et al. 2022 | ERR7448262 |
| NJ-C-7 | NJ-C | NAM.E | NAM | 39.62068 | -75.506 | 7.7610 | 1.0154 | 2009 | Bieker et al. 2022 | ERR7448182 |
| NJ-C-8 | NJ-C | NAM.E | NAM | 39.62068 | -75.506 | 3.2436 | 0.9320 | 2009 | Bieker et al. 2022 | ERR7447517 |
| NJ-G-1 | NJ-G | NAM.E | NAM | 40.77276 | -74.504 | 2.1328 | 0.9274 | 2009 | Bieker et al. 2022 | ERR7447527 |
| NJ-G-2 | NJ-G | NAM.E | NAM | 40.77276 | -74.504 | 6.3335 | 0.9525 | 2009 | Bieker et al. 2022 | ERR7448239 |

| | | | | | | | | | | |
|---|---|---|---|---|---|---|---|---|---|---|
| NS-A-2018-1 | NSA | NAM.E | NAM | 44.9088 | -65.177 | 3.1109 | 0.8764 | 2009 | Bieker et al. 2022 | ERR7447865 |
| NS-A-2018-2 | NSA | NAM.E | NAM | 44.9088 | -65.177 | 2.9942 | 0.9323 | 2009 | Bieker et al. 2022 | ERR7447940 |
| NS-A-2018-3 | NSA | NAM.E | NAM | 44.9088 | -65.177 | 4.0830 | 0.9257 | 2009 | Bieker et al. 2022 | ERR7448012 |
| NSA-1 | NSA | NAM.E | NAM | 44.9088 | -65.177 | 1.1376 | 1.0655 | 2009 | Bieker et al. 2022 | ERR3597474 |
| NSA-3 | NSA | NAM.E | NAM | 44.9088 | -65.177 | 0.9488 | 1.1834 | 2009 | Bieker et al. 2022 | ERR3597476 |
| NSA-6 | NSA | NAM.E | NAM | 44.9088 | -65.177 | 6.8288 | 0.8991 | 2009 | Bieker et al. 2022 | ERR3597483 |
| NSA-7 | NSA | NAM.E | NAM | 44.9088 | -65.177 | 0.9618 | 1.0000 | 2009 | Bieker et al. 2022 | ERR3602088 |
| NSA-8 | NSA | NAM.E | NAM | 44.9088 | -65.177 | 1.2159 | 0.8917 | 2009 | Bieker et al. 2022 | ERR3602090 |
| NSB-6 | NSB | NAM.E | NAM | 44.75472 | -63.672 | 5.7601 | 0.9622 | 2009 | Bieker et al. 2022 | ERR3597482 |
| NSB-7 | NSB | NAM.E | NAM | 44.75472 | -63.672 | 0.8587 | 0.9958 | 2009 | Bieker et al. 2022 | ERR3597472 |
| OH-1 | OH | NAM.ME | NAM | 40.48763 | -82.727 | 3.8546 | 0.9681 | 2013 | Bieker et al. 2022 | ERR7447938 |
| OH-6 | OH | NAM.ME | NAM | 40.48763 | -82.727 | 13.4975 | 0.9653 | 2013 | Bieker et al. 2022 | ERR7448731 |
| OH-B-1 | OH-B | NAM.ME | NAM | 38.90957 | -82.728 | 3.4613 | 0.9543 | 2009 | Bieker et al. 2022 | ERR7447947 |
| OH-B-2 | OH-B | NAM.ME | NAM | 38.90957 | -82.728 | 11.4776 | 0.9653 | 2009 | Bieker et al. 2022 | ERR7448288 |
| ON3-7 | ON3 | NAM.ME | NAM | 45.32971 | -74.893 | 2.3187 | 1.0067 | 2013 | Bieker et al. 2022 | ERR7447993 |
| ON3-8 | ON3 | NAM.E | NAM | 45.32971 | -74.893 | 8.5030 | 0.9787 | 2013 | Bieker et al. 2022 | ERR7448263 |
| PA-1 | PA | NAM.ME | NAM | 40.96589 | -78.175 | 4.5162 | 1.1286 | 2013 | Bieker et al. 2022 | ERR7448022 |
| PA-5 | PA | NAM.ME | NAM | 40.96589 | -78.175 | 11.4825 | 0.9404 | 2013 | Bieker et al. 2022 | ERR7448510 |
| PA-D-1 | PA-D | NAM.E | NAM | 41.31407 | -75.54 | 7.8595 | 0.9669 | 2009 | Bieker et al. 2022 | ERR7448214 |
| PA-D-2 | PA-D | NAM.E | NAM | 41.31407 | -75.54 | 11.6978 | 0.9909 | 2009 | Bieker et al. 2022 | ERR7448733 |

| | | | | | | | | | | |
|---|---|---|---|---|---|---|---|---|---|---|
| PA-E-1 | PA-E | NAM.E | NAM | 40.20361 | -75.906 | 12.8023 | 0.9946 | 2009 | Bieker et al. 2022 | ERR7448742 |
| PA-E-2 | PA-E | NAM.E | NAM | 40.20361 | -75.906 | 14.1769 | 1.0022 | 2009 | Bieker et al. 2022 | ERR7448755 |
| PA-F-1 | PA-F | NAM.ME | NAM | 41.8322 | -80.116 | 21.0436 | 0.9833 | 2009 | Bieker et al. 2022 | ERR7448829 |
| PA-F-2 | PA-F | NAM.ME | NAM | 41.8322 | -80.116 | 10.7888 | 0.9728 | 2009 | Bieker et al. 2022 | ERR7448750 |
| pop2-1 | pop2 | EUR.ME | EUR | 45.40917 | -0.5294 | 1.7380 | 0.9277 | 2019 | Bieker et al. 2022 | ERR7451245 |
| pop2-2 | pop2 | EUR.ME | EUR | 45.40917 | -0.5294 | 2.9742 | 1.0134 | 2019 | Bieker et al. 2022 | ERR7451289 |
| pop2-3 | pop2 | EUR.ME | EUR | 45.40917 | -0.5294 | 3.4648 | 0.9573 | 2019 | Bieker et al. 2022 | ERR7451315 |
| pop3-1 | pop3 | EUR.ME | EUR | 45.47806 | -0.3731 | 2.8549 | 0.9634 | 2019 | Bieker et al. 2022 | ERR7451336 |
| pop3-2 | pop3 | EUR.ME | EUR | 45.47806 | -0.3731 | 6.7457 | 0.9609 | 2019 | Bieker et al. 2022 | ERR7451586 |
| pop3-3 | pop3 | EUR.ME | EUR | 45.47806 | -0.3731 | 4.0197 | 0.9600 | 2019 | Bieker et al. 2022 | ERR7451499 |
| QC-2-1 | QC-2 | NAM.E | NAM | 46.88667 | -70.868 | 15.6584 | 1.0271 | 2013 | Bieker et al. 2022 | ERR7451329; ERR7448740 |
| QC-2-17 | QC-2 | NAM.E | NAM | 46.88667 | -70.868 | 17.6918 | 0.9922 | 2013 | Bieker et al. 2022 | ERR7448790 |
| QC-2-19 | QC-2 | NAM.E | NAM | 46.88667 | -70.868 | 11.3993 | 0.9803 | 2013 | Bieker et al. 2022 | ERR7451307; ERR7448215 |
| QC-2-20 | QC-2 | NAM.E | NAM | 46.88667 | -70.868 | 11.3528 | 1.0596 | 2013 | Bieker et al. 2022 | ERR7451423; ERR7448253 |
| QC-2-22 | QC-2 | NAM.E | NAM | 46.88667 | -70.868 | 10.3134 | 1.0286 | 2013 | Bieker et al. 2022 | ERR7451252; ERR7448286 |
| QC-2-25 | QC-2 | NAM.E | NAM | 46.88667 | -70.868 | 13.9760 | 0.9929 | 2013 | Bieker et al. 2022 | ERR7451297; ERR7448744 |
| QC-2-26 | QC-2 | NAM.E | NAM | 46.88667 | -70.868 | 12.5079 | 1.0792 | 2013 | Bieker et al. 2022 | ERR7451285; ERR7448754 |
| QC-2-28 | QC-2 | NAM.ME | NAM | 46.88667 | -70.868 | 13.7966 | 1.0342 | 2013 | Bieker et al. 2022 | ERR7451417; ERR7448737 |
| QC-2-29 | QC-2 | NAM.E | NAM | 46.88667 | -70.868 | 18.2830 | 1.0502 | 2013 | Bieker et al. 2022 | ERR7451446; ERR7448803 |
| QC-2-30 | QC-2 | NAM.E | NAM | 46.88667 | -70.868 | 26.2481 | 0.9974 | 2013 | Bieker et al. 2022 | ERR7451480; ERR7451456 |

| | | | | | | | | | | |
|---|---|---|---|---|---|---|---|---|---|---|
| QC3-20 | QC-3 | NAM.E | NAM | 47.67876 | -69.022 | 15.8518 | 0.9958 | 2013 | Bieker et al. 2022 | ERR7448889 |
| RI-B-1 | RI-B | NAM.E | NAM | 41.48481 | -71.558 | 6.0052 | 0.9360 | 2009 | Bieker et al. 2022 | ERR7448734 |
| RI-B-2 | RI-B | NAM.E | NAM | 41.48481 | -71.558 | 14.3824 | 0.9371 | 2009 | Bieker et al. 2022 | ERR7448847 |
| SC-1 | SC | NAM.E | NAM | 34.79252 | -80.415 | 17.0874 | 0.9987 | 2009 | Bieker et al. 2022 | ERR7451327 |
| SC-2 | SC | NAM.ME | NAM | 34.79252 | -80.415 | 18.4031 | 0.9924 | 2009 | Bieker et al. 2022 | ERR7451414 |
| SC-K-1 | SC-K | NAM.ME | NAM | 34.22524 | -81.343 | 4.3607 | 1.0264 | 2013 | Bieker et al. 2022 | ERR7448736 |
| SC-K-2 | SC-K | NAM.ME | NAM | 34.22524 | -81.343 | 7.6215 | 1.0367 | 2013 | Bieker et al. 2022 | ERR7448761 |
| StGalmier-1 | StGalmier | EUR.ME | EUR | 45.59 | 4.4 | 14.1354 | 0.9911 | 2017 | Bieker et al. 2022 | ERR7451338 |
| StGalmier-11 | StGalmier | EUR.ME | EUR | 45.59 | 4.4 | 9.6901 | 0.9406 | 2017 | Bieker et al. 2022 | ERR7448872 |
| StGalmier-12 | StGalmier | EUR.ME | EUR | 45.59 | 4.4 | 7.1991 | 0.9848 | 2017 | Bieker et al. 2022 | ERR7448826 |
| StGalmier-16 | StGalmier | EUR.ME | EUR | 45.59 | 4.4 | 10.0433 | 0.9659 | 2017 | Bieker et al. 2022 | ERR7448886 |
| StGalmier-17 | StGalmier | EUR.ME | EUR | 45.59 | 4.4 | 12.9315 | 1.0225 | 2017 | Bieker et al. 2022 | ERR7448881 |
| StGalmier-2 | StGalmier | EUR.ME | EUR | 45.59 | 4.4 | 12.9776 | 0.9387 | 2017 | Bieker et al. 2022 | ERR7451425 |
| StGalmier-21 | StGalmier | EUR.ME | EUR | 45.59 | 4.4 | 6.3308 | 0.9525 | 2017 | Bieker et al. 2022 | ERR7448785 |
| StGalmier-22 | StGalmier | EUR.ME | EUR | 45.59 | 4.4 | 10.0684 | 0.9722 | 2017 | Bieker et al. 2022 | ERR7448878 |
| StGalmier-6 | StGalmier | EUR.ME | EUR | 45.59 | 4.4 | 13.6943 | 0.9648 | 2017 | Bieker et al. 2022 | ERR7451437 |
| TN-1 | TN | NAM.ME | NAM | 36.11943 | -85.059 | 4.4479 | 0.9572 | 2009 | Bieker et al. 2022 | ERR7448786 |
| TN-2019-1 | TN | NAM.ME | NAM | 36.11943 | -85.059 | 2.6949 | 0.9684 | 2009 | Bieker et al. 2022 | ERR7448793 |
| TRH-5 | TRH | EUR.ME | EUR | 58.08278 | 6.75278 | 2.2637 | 0.9819 | 2007 | Bieker et al. 2022 | ERR7460255 |
| TRH-6 | TRH | EUR | EUR | 58.08278 | 6.75278 | 1.7922 | 0.9442 | 2007 | Bieker et al. 2022 | ERR7460286 |

| | | | | | | | | | | |
|---|---|---|---|---|---|---|---|---|---|---|
| TRQB-1 | TRQB | NAM.E | NAM | 45.5 | -72.5 | 9.0505 | 1.0089 | 2013 | Bieker et al. 2022 | ERR7451343 |
| TRQB-10 | TRQB | NAM.E | NAM | 45.5 | -72.5 | 10.9787 | 1.0731 | 2013 | Bieker et al. 2022 | ERR7451326 |
| TRQB-2 | TRQB | NAM.E | NAM | 45.5 | -72.5 | 12.1401 | 1.0382 | 2013 | Bieker et al. 2022 | ERR7451441 |
| TRQB-4 | TRQB | NAM.E | NAM | 45.5 | -72.5 | 8.1634 | 0.9938 | 2013 | Bieker et al. 2022 | ERR7451278 |
| TRQB-7 | TRQB | NAM.E | NAM | 45.5 | -72.5 | 7.8946 | 0.9639 | 2013 | Bieker et al. 2022 | ERR7450221 |
| TRQB-8 | TRQB | NAM.E | NAM | 45.5 | -72.5 | 11.7018 | 1.0032 | 2013 | Bieker et al. 2022 | ERR7451447 |
| TRQB-9 | TRQB | NAM.E | NAM | 45.5 | -72.5 | 10.0110 | 0.9640 | 2013 | Bieker et al. 2022 | ERR7451419 |
| WI-1 | WI | NAM.ME | NAM | 44.87934 | -89.424 | 15.3697 | 1.0095 | 2013 | Bieker et al. 2022 | ERR7451468 |
| WI-3 | WI | NAM.ME | NAM | 44.87934 | -89.424 | 6.7306 | 0.9665 | 2013 | Bieker et al. 2022 | ERR7451282 |
| WU-15 | WU | EUR | EUR | 47.98833 | 16.2831 | 6.2109 | 0.9683 | 2010 | Bieker et al. 2022 | ERR7451424 |
| WU-16 | WU | EUR.ME | EUR | 47.99444 | 16.3294 | 2.9881 | 0.9224 | 2016 | Bieker et al. 2022 | ERR7451249 |
| WU-17 | WU | EUR.ME | EUR | 48.20028 | 16.455 | 2.9390 | 0.9274 | 2013 | Bieker et al. 2022 | ERR7451269 |

**Suppl. Table S10: Descriptions of Bioclimatic Variables (BIO1-BIO19) from WorldClim.**

| Bioclimatic Variable | Description | Unit |
|---|---|---|
| BIO1 | Annual Mean Temperature | °C |
| BIO2 | Mean Diurnal Range (Mean of monthly (max temp - min temp)) | °C |
| BIO3 | Isothermality (BIO2/BIO7) (*100) | Dimensionless (%) |
| BIO4 | Temperature Seasonality (Standard deviation *100) | °C |
| BIO5 | Max Temperature of Warmest Month | °C |
| BIO6 | Min Temperature of Coldest Month | °C |
| BIO7 | Temperature Annual Range (BIO5-BIO6) | °C |
| BIO8 | Mean Temperature of Wettest Quarter | °C |

| | | |
|---|---|---|
| BIO9 | Mean Temperature of Driest Quarter | °C |
| BIO10 | Mean Temperature of Warmest Quarter | °C |
| BIO11 | Mean Temperature of Coldest Quarter | °C |
| BIO12 | Annual Precipitation | mm |
| BIO13 | Precipitation of Wettest Month | mm |
| BIO14 | Precipitation of Driest Month | mm |
| BIO15 | Precipitation Seasonality (Coefficient of Variation) | Dimensionless (%) |
| BIO16 | Precipitation of Wettest Quarter | mm |
| BIO17 | Precipitation of Driest Quarter | mm |
| BIO18 | Precipitation of Warmest Quarter | mm |
| BIO19 | Precipitation of Coldest Quarter | mm |
